# Modular Design and Scaffold-Synthesis of Multi-Functional Fluorophores for Targeted Cellular Imaging and Pyroptosis

**DOI:** 10.1101/2024.05.24.595706

**Authors:** Yuanyuan Li, Chunhui Wang, Haiyang Wang, Kunhui Sun, Siyu Lu, Yahui Wang, Lei Zhang, Su Jing, Thorben Cordes

## Abstract

Modified commercial fluorophores are essential tools for optical imaging and biomedical research. Their synthetic modification to incorporate new functions, however, remains a challenging task. Conventional strategies rely on linear synthesis in which a parent framework is gradually extended. We here designed and synthesized a versatile library of functional fluorophores via a scaffold-based Ugi four-component reaction (U-4CR). The adaptability of the scaffold is achieved through modification of starting materials. This allows to use a small range of starting materials for the creation of fluorogenic probes that can detect reactive-oxygen species and where the localization into subcellular organelles or membranes can be controlled. We present reaction yields ranging from 60% to 90% and discovered that some compounds can even function as imaging and therapeutic agents via Fenton chemistry inducing pyroptosis in living cancer cells. Our study underlines the potential of scaffold-based synthesis for versatile creation of functional fluorophores and their applications.

## INTRODUCTION

The development of advanced fluorescent materials with diverse functional properties has facilitated progress in bioimaging, molecular biology, biochemistry and biomedicine.^[1]^ The used water-soluble fluorescent dyes or fluorophores are not just signal units, but can act as local probes or might even participate in the regulation of biochemical processes.^[2]^ They are capable of imaging specific molecules in living cells or tissue, greatly assisting in the ability to intervene diseases.^[3]^ Recently, the development of theranostic systems that allow simultaneous diagnostic imaging and therapy has been presented.^[4]^ For many applications, a fluorophore with multiple functions, e.g., signal generation and modulation by specfic biochemical events, target binding, or theranostic activity, is required.^[5]^ Thus, preparing fluorophores with excellent optical properties, targetability, and specific functional properties is critical for applications, e.g., in disease identification or treatment.^[6]^

The synthesis and modifications of such multifunctional fluorophores, however, remains a challenging task.^[7]^ Many conventional strategies rely on linear synthesis^[8]^, in which a parent framework (often a fluorophore) is gradually extended through the addition of functional groups (**Figure 1**, top) to allow bio-recognition (targeting), signal regulation or biochemical interactions, e.g., via drug units. By using this strategy, several multifunctional fluorogenic probes (fluorescent turn-on probes) have been developed for bioimaging and diagnostics.^[9]^ Examples are fluorescent probes with near-infrared II emission and high brightness for high signal-to-noise imaging-guided diagnosis based on a cyanine dyes backbone.^[10]^ Another example involves fluorogenic acedan dyes, which were coupled to triphenylphosphonium and morpholine as targeting groups.^[11]^ These modifications require separate and complex synthetic procedures, both starting from the fluorophore, to obtain probes that allow for capable of performing functions in a unidirectional manner within application contexts. There are many other examples for biological applications of similar strategies^[12]^ and it should be stressed that extensive optimization of reaction conditions is required before a functional dye, can be obtained which often hinders the application of such materials beyond basic research. An ideal solution is a modular synthesis^[13]^ of a framework that connects the fluorophore and all desired functional groups (**Figure 1**, middle). Importantly, the reaction towards such a scaffold is much more direct compared to linear approaches, it can provide the functional fluorophore in just a few synthetic steps and allows for the specific exchange of functionalities in the framework. Such reactions have been used to built-up of many different functional light-absorbing chromophores using a scaffolding- or chomorophore-approach.^[14]^ Its use for the design of functional fluorescent dyes, however, is still scarce.^[15]^ The Urano group has established a modular platform for the detection of carboxypeptidases utilizing activatable fluorescent probes.^[15]^ A linear synthesis was used with the necessity to subsequently tailor the fluorophore to match specific requirements. Here, we introduce an “editable” platform for scaffolded synthesis and customization of target-specific fluorogenic dyes via the Ugi four-component reaction (U-4CR), which can be extended easily to other dye classes and functionalities (**Figure 1**, bottom).

**Figure 1.**
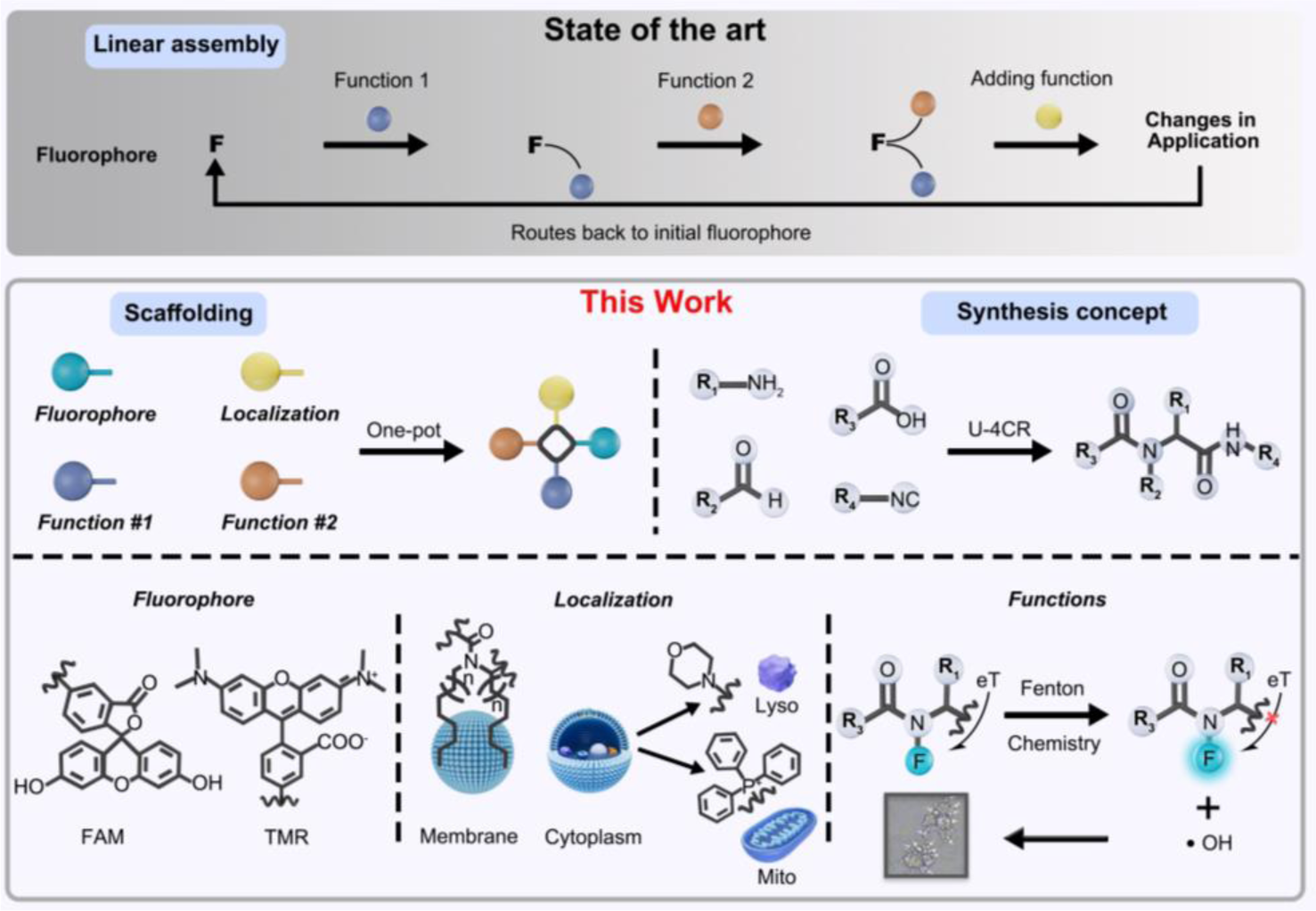
Comparison of linear (top) and scaffold (middle) synthesis routes towards multi-functional fluorescent dyes with fluorogenic properties and the ability for cellular localization, imaging and pyroptosis (bottom).

The U-4CR offers superb scaffold diversity to obtain complex chromophore structures.^[16]^ It combines four reaction partners in a single step and reaction vessel in an atom-efficient manner without the need for catalysts.^[17]^ Based on the U-4CR fluorescent peptidomimetics,^[18]^ pharmacophores,^[19]^ and functional chromophores were prepared^[13b]^. Müller and co-workers pioneered the application of the U-4CR for various chromophores including highly functional donor-acceptor dyads.^[14b]^ Westermann and colleagues used the U-4CR to synthesize rhodamine tags with clickable bioorthogonal groups for biolabeling.^[20]^ Instead of utilizing four components, Vendrell et al. proposed a synthesis based on the Passerini three-component reaction (P-3CR) to create a BODIPY-containing isonitrile fluorescent probe for low-pH sensing in activated macrophages.^[21]^ We recently utilized the U-4CR to create a small library of linker molecules for labeling of biomacromolecules with fluorescent dyes in a two-step protocol, an approach that allowed to convert commercial fluorophores into functional probes with high photostability and metal sensing ability, directly on the biological target.^[22]^

Building on the rising interest and success of scaffolded functional fluorophores^[23]^, we here introduce a modular synthesis platform for fluorogenic probes based on the U-4CR. The obtained probes combine three distinct moeities, i.e., a fluorophore, a signal modulator and a unit used for cell labelling of specific targets (the cell membrane, mitochondria and lysosomes). Our approach uses the redox properties of ferrocene (Fc) ^[24]^, and its influence on organic dyes via photoinduced electron transfer (PeT) ^[25]^ for signal modulation. Furthermore, Fc is involved in a Fenton-like reaction (the generation of hydroxyl radicals from hydrogen peroxide) that is stimulated by the microenvironment of cancer cells.^[26]^ These hydroxyl radicals are reactive oxygen species capable of inducing cell death.^[27]^ More specifically, in cancer cells, the presence of mild acidic conditions triggers the initiation of a Fenton-like reaction, leading to the excessive consumption of hydrogen peroxide in order to generate hydroxyl radicals. This reaction provides a certain degree of safety to normal cells, as the Fenton reaction is significantly inhibited under slightly alkaline conditions and when there is an insufficient amount of hydrogen peroxide in a normal environment.^[26]^ We demonstrate here that fluorogenic probes synthesized via the U-4CR scafold strategy, which target either cytoplasmic lysosomes or cytoplasm mitochondria, can actually activate pyroptosis^[27]^ pathways through photo-Fenton chemistry, evidenced by the activation of caspase-1 and -3 and the cleavage of gasdermin D and E. This underscores the versatility of the modular scaffold strategy, for modifying fluorescent probes not only for imaging but also for biomedical applications.

## RESULTS

### Scaffolding of Multi-Functional Fluorogenic Dyes via U-4CR

The U-4CR method involves the one-pot reaction of an aldehyde, an amine, an isocyanide, and a carboxylic acid to produce *α*-*N*-acetoamido carboxamide derivatives as shown in **Scheme 1**. This approach allows to combine up to four functional groups within the carboxamide scaffold. We employed this approach to obtain fluorogenic probes where fluorescence becomes activated (off→on) under specific chemical conditions in cellular organelles. Our goal was to obtain a series of fluorogenic probes with a scaffold that can be readily modified by altering the starting materials to achieve distinct cellular localization. The probes included a localization moiety comprising either triphenylphosphonium, morpholine, or a hydrophobic aliphatic chain and a fluorophore.

**Scheme 1.**
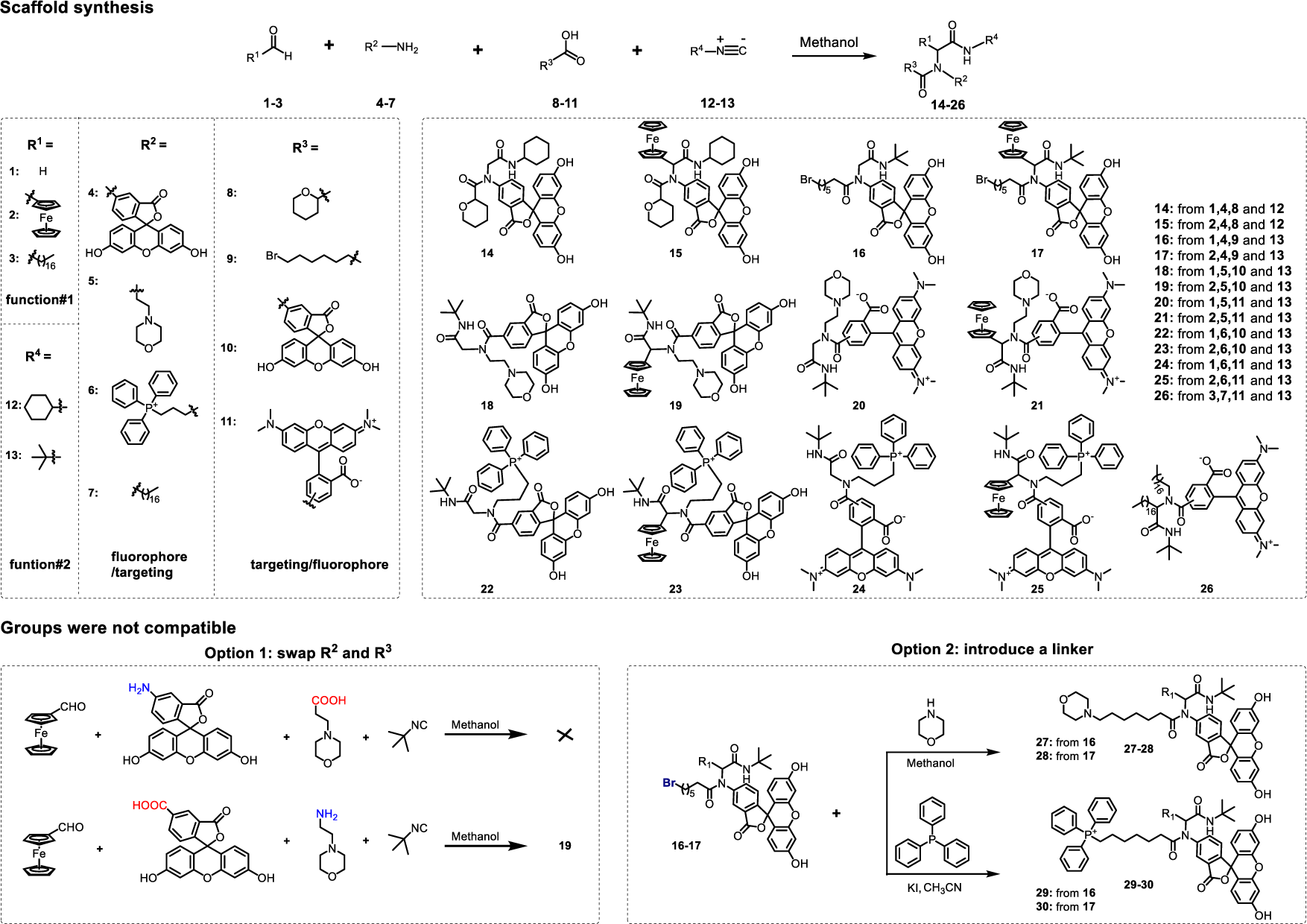
Synthesis route of multi-functional fluorogenic dyes starting with the fluorophore (**4,10,11**), targeting localization group (**3, 5–7**), Fc-PeT units (**2**) and isocyanide (**12** and **13**). Compound **1** lacks the Fc moiety and was used as a control. Compounds **14–26** were obtained by one-pot 4-UCR. Compound **16–17** was used as a linker for compounds **27–30**.

This enabled the fluorogenic probes to specifically anchor on the phospholipid membrane through hydrophobic interactions between alkane chain^[28]^, or to localize to mitochondria via triphenylphosphonium ^[29]^ and lysosomes via morpholine^[30]^, respectively. Additionally, a xanthene fluorophore and a functional ferrocenyl module were incorporated to allow fluorogenic imaging upon ferrocenyl oxidation or photo-Fenton triggered pyroptosis in cancer cells. We used a hydrogen atom (compounds **14, 16, 18, 20, 22, 24, 27** and **29**) as non-functional negative control. Importantly, all starting materials used here were obtained commercially and did not have to be synthesized. For this, the reaction of four components (one of **1–3**, one of **4–7**, one of **8–11** with either **12** or **13**) was performed in methanol for 48 hours, resulting in the formation of *α*-N-acylamino amides **14–26**. The reaction yields for compounds **14** and **19** were high and exceeded 80%, while those for compounds **15–18, 20, 21, 23–26** were between 60–80%, with on exception of compound **22** (∼50%). Compounds **14–17** were purified by silica gel column chromatography, **18–25** were purified using a preparative HPLC, and **26** was purified by recrystallization. All compounds were subsequently characterized (Supplementary Information, Part I/II, **Figures S1–5, S10–13, S18–32**). While some functional dyes could be synthesized in one step (**Scheme 1, 14–26**), others (**Scheme 1, 27–30**) required additional steps, since some functional groups were not compatible with the U-4CR reaction. For example, we observed that a mixture of a fluorophore with an amino group (compound **4**), a cell-targeting morpholine with a carboxylic acid, ferrocenecarboxaldehyde (compound **2**), and *n*-Butyl isocyanide (compound **13**) did not undergo the U-4CR reaction. The modularity of our approach, however, allows to swap the functional moieties in the precursors, illustrated in **Scheme 1** (bottom, left), which shows an optimization of the conditions to obtain compounds such as **18** and **19** in a single step. Using a fluorophore with a carboxylic acid (compound **10**) and a morpholino amino group (compound **5**), under otherwise identical conditions, allowed the one-pot reaction to proceed with good yields. Secondly, we used a more complex scheme that started with compounds **16** and **17** and then introduced a linker allowing the isolation and purification of products **27–30** (**Scheme 1**, bottom, right) via column chromatography. The yields for these were 50–80% (**Figures S6–9, S14–17, S33–40**). These variations show that the scaffold synthesis based on U-4CR allows a straightforward and effective optimization of reaction conditions by variation of the starting precursors.

### Spectroscopic Properties of Scaffolding-Based Fluorogenic Dyes

Prior to the U-4CR conjugation of relevant ferrocene (Fc) moieties to fluorophores, we investigated the quenching of dyes via hybridization in a double-stranded DNA (**Figure 2A**) in bulk fluorescence measurements. The proximity of ferrocene induces fluorescence quenching for blue/green-absorbing fluorophores (**Figures 2B, 2C**). The electron-rich ferrocene moiety serves as an electron donor from the low-spin iron (II) center for the photoexcited fluorophore^[31]^ and efficiently suppresses fluorescence emission via photoinduced electron transfer (PeT)^[32]^. Upon oxidation of the ferrocence moiety, the PeT mechanism is disrupted, leading to a recovery of fluorescence (**Figures 2D, 2E**). This effect was most pronounced for fluorescein (FAM) and tetramethylrhodamine (TMR). Therefore, FAM and TMR were selected as signal units in our dye U-4CR conjugates (**14–30**). Among these, the conjugated triphenylphosphonium-FAM-Fc compound **30** showed significantly lower fluorescence than the triphenylphosphonium-FAM-control (compound **29, Figure 2I**). This trend was also observed in the morpholine-FAM probes (compound **27–28, Figure 2F**). Further support for the removal of PeT quenching is provided by fluorescence recovery of compounds **28** and **30** upon addition of H_2_O_2_ (**Figures 2H, 2K**). A similar effect did not occur in the absence of Fc conjugation for compounds **27** and **29** (**Figures 2G, 2J**).

**Figure 2.**
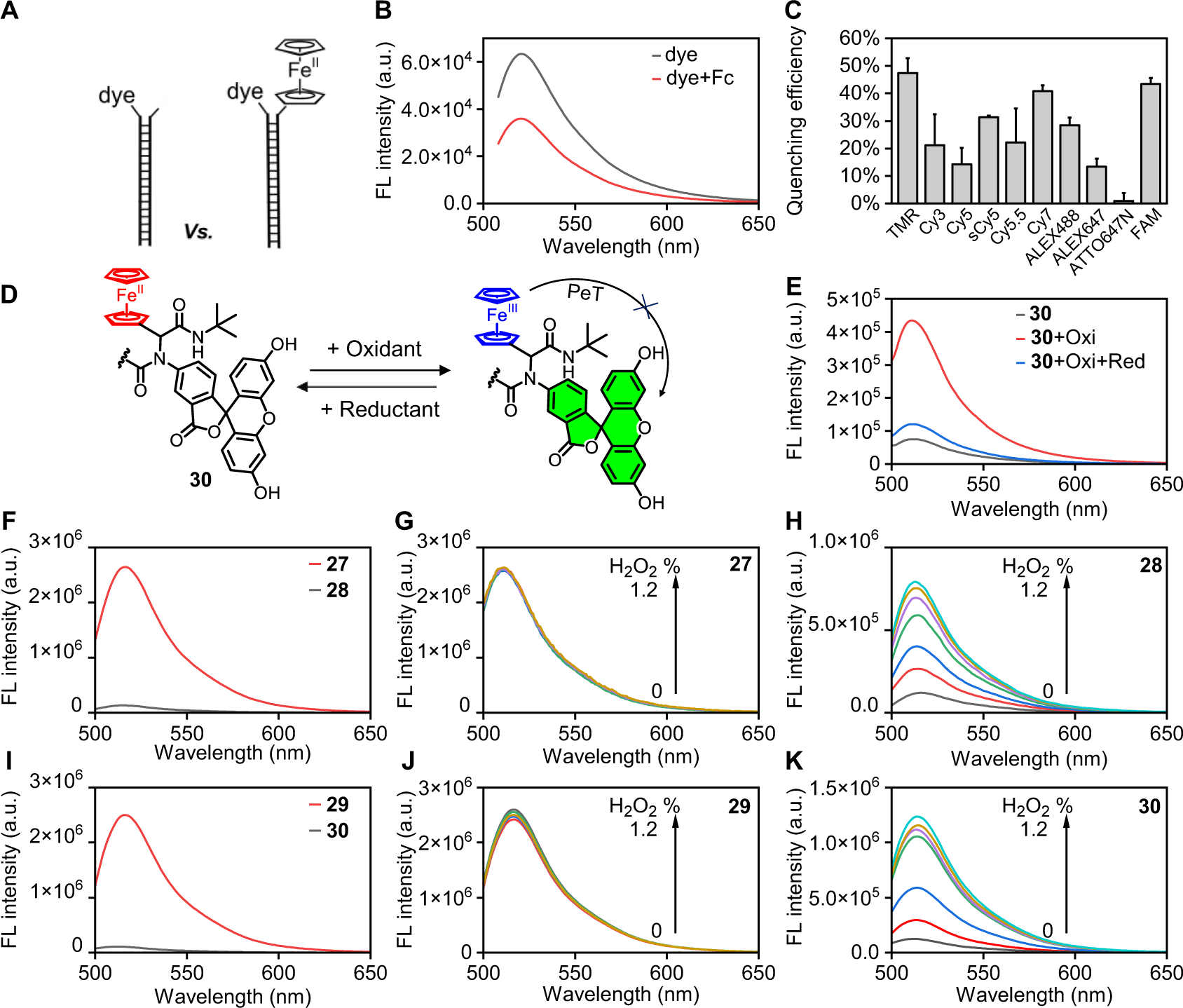
Spectroscopic characterization of compounds **27–30**. (A) The influence of ferrocene on various fluorophores when terminally attached to dsDNA in (B, C) bulk fluorescence measurements. (D) Schematic illustration of PeT quenching and (E) fluorescence spectra of compound **30**, showing the reversible nature of PeT quenching by the sequential addition of 1.0 equivalent of an oxidant and followed by a reductant. (F–K) Fluorescence spectra of compounds **27–30** at identical concentration and under varying concentrations of hydrogen peroxide (0, 0.03, 0.06, 0.12, 0.3, 0.6 and 1.2%) in pH 6.5 PBS buffer solution.

Direct evidence for PeT quenching was obtained from pump-probe transient absorption spectroscopy (TAS) with picosecond time resolution.^[33]^ For compounds **27–30** (**Figure 3A**), the fluorescein fluorophore was consistently positioned at the N-terminal end of the U-4CR scaffold. Ferrocene was located at the carbon atom of the aldehyde functional group in compounds **28** and **30**. Compounds **27** (Fc-free control for **28**) and **28** are characterized by the presence of a morpholine group, while compounds **29** (Fc-free control for **30**) and **30** incorporate a triphenylphosphonium group, respectively. According to the steady state absorption spectra (**Figure S41**), the pump wavelength was set to 490 nm. These synthesized U-4CR dyes based on fluorescein (**Figures 3B, 3C, 3F, 3G** and **Figures S42, S43**) showed similar TAS absorbance difference spectra (ΔA) and clear spectral signatures of excited-state absorption (ESA, positive signal around 410 nm) and two overlapping negative contributions of ground-state bleaching (GSB) and stimulated emission (SE) with a minimum at around 498 nm. For **28** and **30** the complete ΔA signal vanished with a dominant decay time of 30 ps (single exponential fit y(t) = a*exp(-t/τ) ^[34]^ indicating that the excited state is rapidly quenched by PeT. For **27** and **29**, we used a double exponential fit function of the form y(t) = a_1_*exp(-t/τ_1_)+a_2_*exp(-t/τ_2_)^[35]^ showing a fast 30 ps component and a slow decay with a time constant of 4 ns. The latter corresponds to the decay of the relaxed excited state matching the fluorescence lifetime of free fluorescein.^[36]^ The appearance of the shorter-lived component in the initial tens of picoseconds for **29** and **30** can be attributed to the presence of the anionic form of the fluorescein fluorophore, which coexists in a mixture with its dianion at pH 7.4.^[37]^

**Figure 3.**
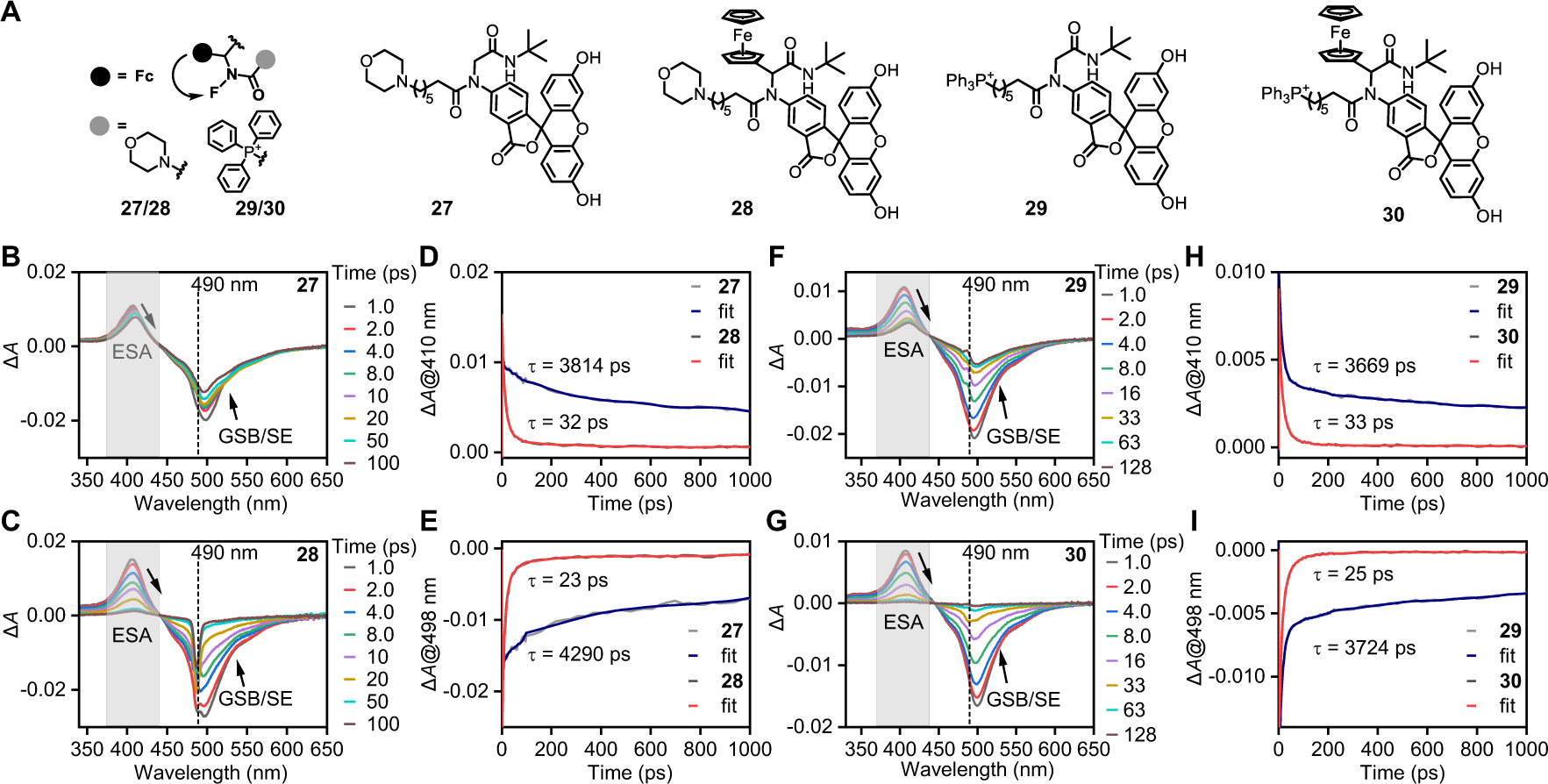
Transient absorption spectroscopy of dye conjugates **27-30** with broad-band probing and excitation at 490 nm.

A comparison of the normalized kinetics of compound **27**/**29** (without ferrocene conjugation) and **28**/**30** (with ferrocene conjugation), as shown in **Figures 3D, 3E, 3H** and **3I**, allow an assessment of the electron transfer rate *k*_eT_. The rate *k*_eT_ was calculated according to *k*_eT_=1/τ_Fc_-1/τ based on both ESA and GSB/SE kinetics. We found values of *k*_eT_ of 3.7 × 10^10^ s^-1^ for **28** and 3.5 × 10^10^ s^-1^ for **30**. These high rates are indicative of fast electron transfer and are compatible with the large negative free energy difference ΔG for charge separation of -0.84 eV, calculated with the Rehm-Weller equation^[38]^: ΔG = e*[E_ox_ − E_red_] − E_0,0_ + C; with the unit charge e^-^, the one-electron oxidation potential of donor (E_ox_ = 0.45 V) and reduction potential of acceptor (E_red_ = -1.23V) and the zero–zero energy (E_0,0_ = 2.52 eV). For the calculation of ΔG we neglected the solvent-dependent coulombic attraction energy (C) due to the experiments being conducted in a polar solvent.

The comparison of electron transfer rates in different functional fluorophore constructs highlights the possibilities of the scaffold-based strategy to optimize their photophysical properties. We could show that the U-4CR scaffold allows for a simple exchange of the biotargeting modules, i.e., switching from morpholine to triphenylphosphonium, without significant alterations of the electron transfer rate (**Figure 3**). We can demonstrate, however, that the scaffold strategy also allows for the adjustment of the relative positions between the PeT donor and the fluorophore. Our TAS analysis of compounds **17**/**28/30**, where the fluorophore is positioned at the N-terminal, exhibit higher PeT rates compared to compound **19** (7.3 × 10^8^ s^-1^), which has its fluorophore located at the C-terminal of the scaffold (**Figures S42–S45**).

### Cellular Imaging with Fluorogenic Dyes

The resulting compounds **26–30** comprise distinct cellular localization groups on the scaffold. These include hydrophobic alkane chains, morpholine, and triphenylphosphonium moieties. Each of them is known for its selective cellular localization, ranging from the membrane to lysosomal targeting (morpholine group), and mitochondrial localization via the triphenylphosphonium group (**Figure 4A**). To verify the specific cellular localization of our compounds, we incubated them with living HeLa cells and imaged those with confocal microscopy. **Figure 4B** illustrates that HeLa cells treated with **26** showed uniform staining of the cytomembrane. This indicates a hydrophobic interaction between **26** and the plasma membrane. In contrast, **27–30** showed a distinct intracellular localization in HeLa cells. Since our compounds exhibit green fluorescence, we checked their co-localization with the red fluorescence of commercially available Lyso- and Mito-Tracker dyes (**Figures 4C–F)**. These experiments show that compounds **27** and **28** specifically targeted lysosomal compartment, while compounds **29** and **30** are found in mitochondrial compartments. Importantly, the integration of the ferrocenyl module into these compounds does not affect their localization characteristics.

**Figure 4.**
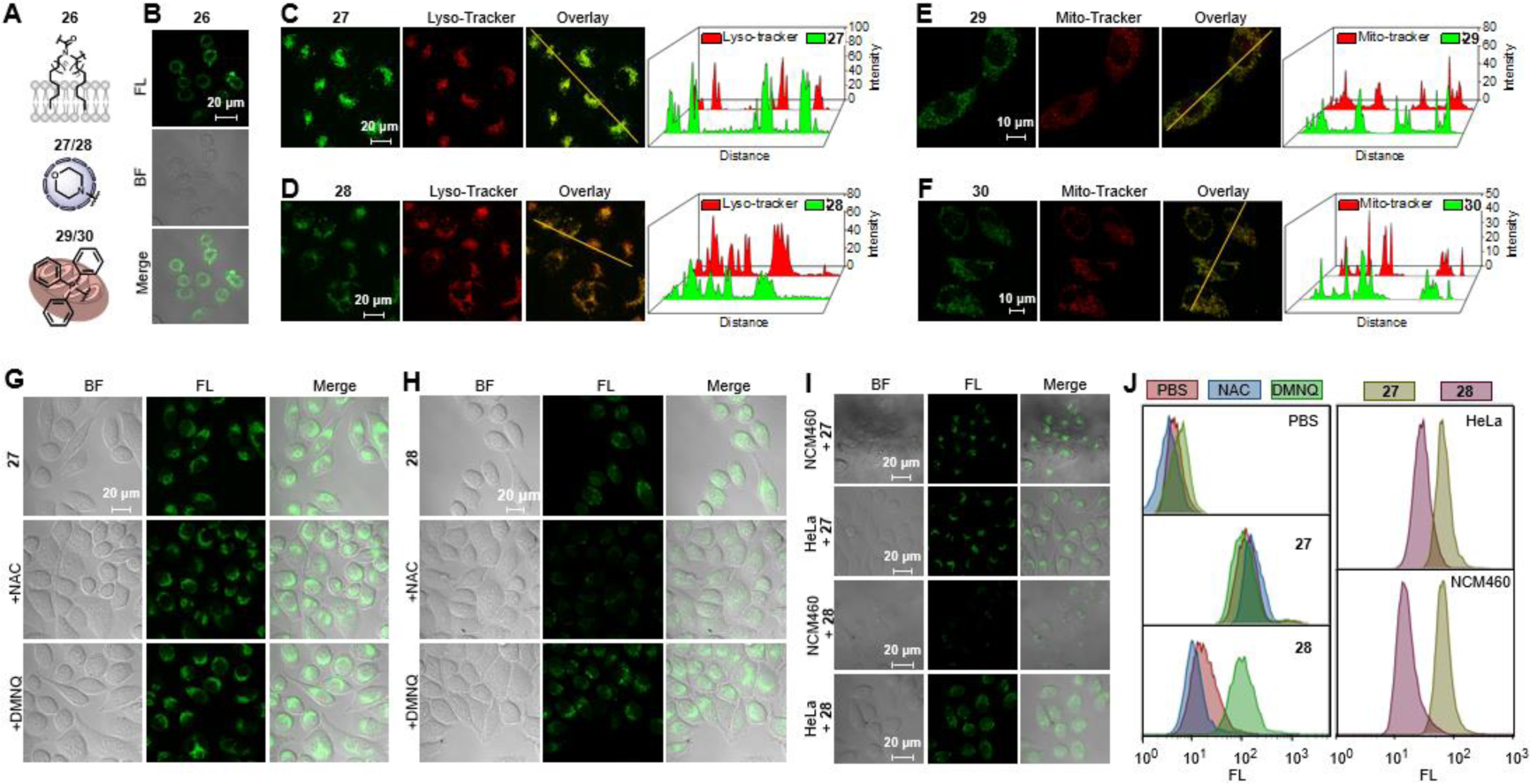
Targeted localization of HeLa cells achieved by staining with compounds **26–30** via confocal microscopy. (A) An illustration of distinct compounds with customized cellular localization capabilities. (B) Confocal microscopy imaging of HeLa cells stained with compound **26** (10 μM). (C, D) Confocal microscopy images of HeLa cells labeled with **27**/**28** (1.0 μM); colocalization with Tracker Red (Lyso-Tracker), Scale bar = 20 μm. (E, F) Confocal microscopy images of HeLa cells labeled with **29**/**30** (1.0 μM); colocalization with Tracker Red (Mito-Tracker), Scale bar = 10 μm. (G, H) Confocal microscopy images of **27** (G) and **28** (H)-incubated HeLa cells pretreated with different chemical agents: NAC (H_2_O_2_ inhibitor), DMNQ (H_2_O_2_ agonist), Scale bar: 20 μm. (I) Confocal microscopy images of **27**- and **28**-incubated HeLa cancer and NCM460 cells, Scale bar: 20 μm. (J) Flow cytometry analysis of HeLa cells incubated with compound **27** and **28** after pre-treatment with NAC or DMNQ, and comparative analysis between the treated HeLa cancer cells and NCM460 cells.

Furthermore, the inclusion of a ferrocenyl group in the main structure promotes a Fenton-like response to intracellular hydrogen peroxide, from which we expect a removal of the PeT effect and subsequent recovery of fluorescence. To validate the specificity and responsiveness of this system towards hydrogen peroxide under cancerous conditions, comparative studies were conducted using NCM460, a normal cell line, and HeLa, a cancer cell line. Typically, the H_2_O_2_ concentration in normal cells is around 20 nM, whereas cancer cells can have values up to 0.4 mM.^[39]^ Our analysis, conducted using confocal microscopy and flow cytometry (**Figures 4I, 4J**), demonstrated a distinct behaviour of the compounds when tested across the two cell lines. Compound **27**, lacking the ferrocene conjugation, exhibited uniform fluorescence across both cell types, indicating a lack of selective response to varying concentrations of H_2_O_2_. In stark contrast, compound **28**, which contains the ferrocene moiety, demonstrated reduced fluorescence in the low-H_2_O_2_ environment characteristic of normal cells. Conversely, it displayed significantly increased fluorescence in HeLa cells, where H_2_O_2_ levels are notably elevated. This differential fluorescence response underscores the sensitivity of **28** to H_2_O_2_, highlighting its potential for cancer cell-specific detection. Data obtained from flow cytometry analysis, after comprehensive cell staining, further corroborated the distinct behaviour of compounds **27** and **28** within cellular environments characterized by varying levels of H_2_O_2_. Additionally, the fluorescence of **28-**stained HeLa cells were markedly reduced upon pre-incubation with the H_2_O_2_ inhibitor N-acetylcysteine (NAC) ^[40]^, while pre-treatment with an H_2_O_2_ oxidative stress inducer, 2,3-dimethoxy-1,4-naphthoquinone (DMNQ) ^[41]^, enhanced the fluorescence of **28-**stained cells (**Figure 4H**). Notably, this oxidative stress treatment did not alter the fluorescence of **27**-stained HeLa cells (**Figure 4G**). Importantly, independent application of either NAC or DMNQ directly to the cell culture medium did not affect the fluorescence of compounds **27** and **28** (**Figure S46**). These results suggest that compound **28** is capable of responding specifically to H_2_O_2_ through Fenton-like chemistry within cancer cells.

### Biomedical Applications

Hydroxyl radicals produced by Fenton-like chemistry from H_2_O_2_ have potential applications in impeding cancer cell proliferation.^[26, 43]^ We here employed a CCK-8 assay as a benchmark method for evaluating cell viability in the presence of our compounds (**Figure 5A**). Specifically, HeLa cells were co-incubated with compound **30** for 24 hours, resulting in a substantial reduction in cell viability to approximately 44%. The half-maximal inhibitory concentration (IC_50_) value of **30** against the HeLa cell line was 22±2 μM, demonstrating a moderate inhibitory effect. Exposing **30**-incubated cells to a blue LED for 30 minutes, however, decreased cell viability significantly, yielding a lower IC₅₀ value of approximately 8.5±0.5 μM. In contrast, under control conditions, either with light irradiation in the absence of any compound or with the inclusion of compound **29** (without Fc conjugation), we observed no cytotoxicity on the timescale of imaging. This indicates that light exposure, in combination with **30**, enhances Fenton-like reactions, thereby generating an increased number of hydroxyl radicals.^[43]^ This increase of intracellular hydroxyl radicals was directly observed using the Cell Meter™ detection kit, featuring a commercially available fluorescent probe designed for the specific detection of intracellular hydroxyl radicals. **Figure 5B** shows the red signal of the Cell Meter™ dye, which is indicative on the presence of hydroxyl radicals in the cytoplasm. To substantiate that these radicals were indeed localized in the cytoplasm, DAPI^[44]^, a near-UV dye, was employed to stain and define the cytoplasmic area. Notably, the Cell Meter™ signal increased for prolonged light exposure, in the presence of compound **30**. Conversely, compound **29** did not show similar effects.

**Figure 5.**
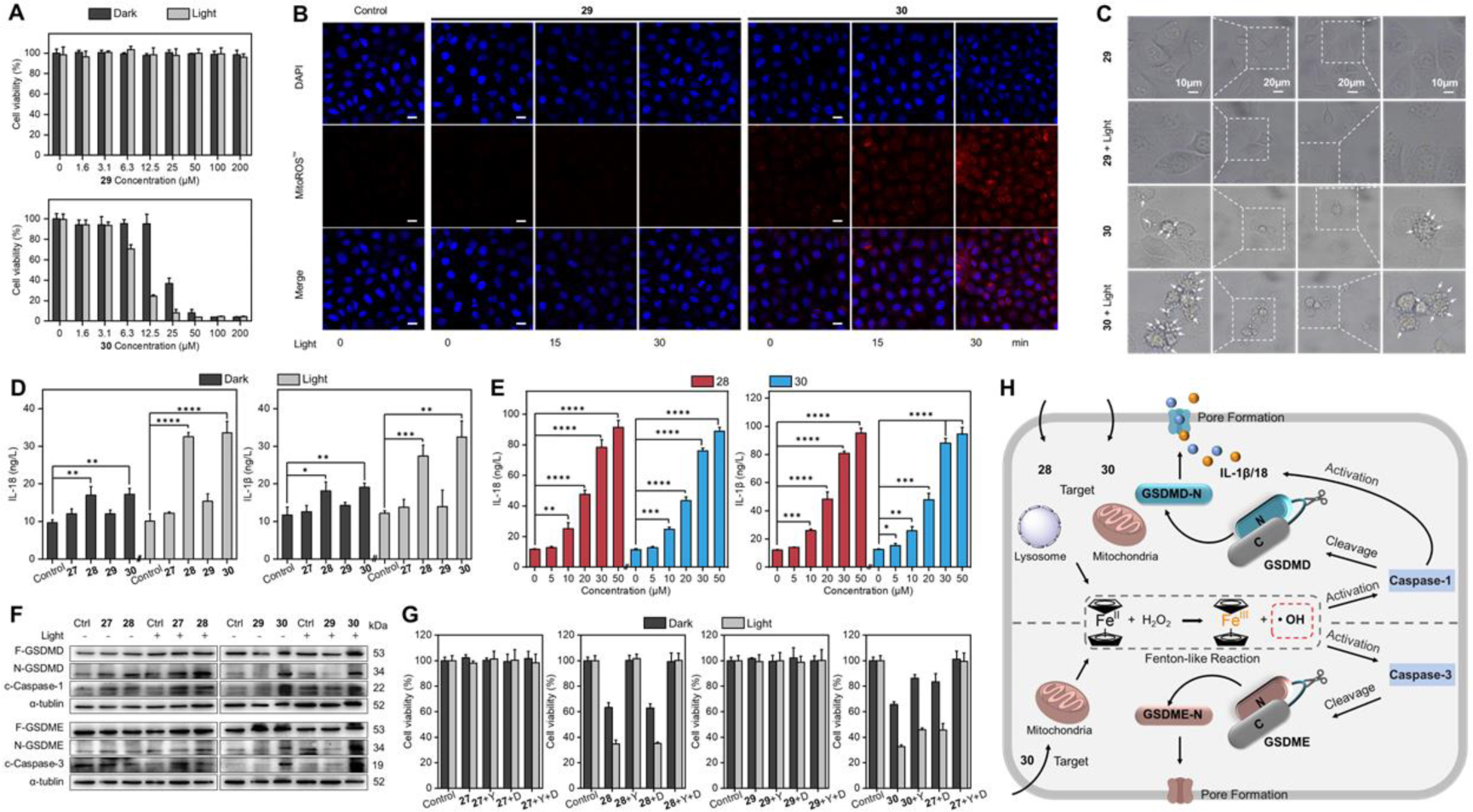
Cancer cellular lysosome-/mitochondria-targeted pyroptosis induced by compounds **28** and **30**. (A) Cell viability analysis of compound **29**- and **30**-mediated toxicity against HeLa cells measured by CCK-8. The data are expressed as mean ± standard deviation, based on three independent experiments for each group. (B) Fluorescent imaging of intracellular hydroxyl radicals using the Cell Meter™ hydroxyl radical detection kit in the absence and presence of light irradiation. (C) Representative phase-contrast images of HeLa cells after treatment with **29** and **30**. White arrows indicate cells with pyroptotic morphology. (D) ELISA analysis of IL-1β and IL-18 secretion into the supernatant of HeLa cells treated with **27,28,29** and **30** with and without light exposure. (E) Correlation between the concentrations of compounds **28** and **30** and the secretion levels of IL-1β and IL-18 into the supernatant of treated HeLa cells. (F) Western blots of cleavage of GSDM and Caspase in response to **27,28,29** and **30**-treated HeLa cells. (G) Relative cell viability of HeLa cells treated with **27–30** in presence of either YVAD (caspase-1 inhibitor) or/and DEVD (caspase-3 inhibitor). (H) A schematic representation elucidates the pyroptosis pathway induced in HeLa cells by treatment with compound **28** and **30**. Statistical significance was determined using one-way analysis of variance (ANOVA) followed by Tukey’s test. *p<0.05, **p< 0.01, ***p<0.001, ****p <0.0001.

Cell viability was further refined through phase-contrast imaging (**Figure 5C**). Cells treated with **29**, designated as the positive control, uniformly presented intact cell membranes, indicating no discernible damage. Treatment with **30** induced morphological alterations, characterized by significant swelling and a unique cellular morphology: a centralized nucleus encircled by an expansive cytoplasmic zone. After 15 minutes of light exposure, these morphological changes were clearly visible with multiple protrusions on the cell membrane with diameters ranging from approximately 2 to 5 μm. Such morphological features are reminiscent of those observed in pyroptosis^[45]^, a form of programmed cell death recently delineated by the caspase-mediated cleavage of proteins from the gasdermin family.

To substantiate the hypothesis that our compounds can induce pyroptosis, we quantified key biomarkers associated with this form of cell death, specifically the inflammatory cytokines interleukin-1β (IL-1β) and interleukin-18 (IL-18). This quantification was conducted on cancer cells treated with compounds **27–30**, utilizing ELISA kits (**Figure 5D**). The results showed an increased release of IL-1β and IL-18, especially when exposed to compounds **28** and **30** in conjunction with light irradiation. Importantly, a positive correlation between the levels of IL-1β and IL-18 and the concentrations of the compounds was observed (**Figure 5E**), reinforcing the potential link between these compounds and the activation of pyroptotic pathways. In stark contrast, cells treated with compounds **27** and **29**, which lack Fc, exhibited negligible release of these cytokines, even when exposed to light. This supports the role of Fc present in compounds **28** and **30** for pyroptosis.

A Western blot analysis further supported these findings, offering insights into the mechanism by which these compounds induce pyroptosis (**Figure 5F**). In SDS-PAGE, a significant alteration in gasdermin D (GSDMD) dynamics was observed in cells treated with compound **28** or **30**. Specifically, there was a noticeable reduction in the levels of full-length GSDMD and a concurrent increase in the GSDMD-N fragment, which was cleaved by activated caspase-1. This change was corroborated by the upregulation of cleaved caspase-1 (c-caspase-1). Both compounds **28** (lysosome-targeting) and **30** (mitochondria-targeting) were shown to induce pyroptosis via a classic caspase-1-dependent pathway. However, in the case of pyroptosis induced by compound **30** with mitochondrial targeting, caspase-3 activation was also responsible for the cleavage of gasdermin E (GSDME). Such complexity underscores the potential of differential regulatory mechanisms in pyroptosis initiated at these subcellular sites (**Figure 5H**). Importantly, the modular design of our experimental approach offers an advantage. It enables the preservation of a unique functional while permitting alterations of cellular localization groups. This methodology not only underscores the importance of precise molecular control in elucidating complex cellular processes, but also enhances our understanding of the role of the caspase family in pyroptosis.

To elucidate the role of the caspase family in the pyroptosis pathway initiated by compounds **27–30**, we conducted caspase inhibition studies. Specifically, HeLa cells underwent pretreatment with z-YVAD-FMK (YVAD), a caspase-1 inhibitor, and/or DEVD, a selective inhibitor of caspase-3, before being exposed to these compounds (**Figure 5G**). Cells pretreated with YVAD exhibited a significant enhancement in resistance to cell death following exposure to compounds **28** and **30**. In contrast, DEVD pretreatment afforded a protective effect specifically in response to compound **30**. The concurrent administration of YVAD and DEVD yielded notable cytoprotective efficacy, substantially attenuating cytotoxicity induced by compound **30**. These results indicate the critical involvement of caspase-1 in the pyroptosis pathway triggered by compound **28** and delineate the synergistic roles of caspase-1 and caspase-3 in the cell death process elicited by compound **30**. Conversely, in control experiments, neither inhibitor improved cell viability in the presence of compound **27**, which targets lysosomes yet lacks a functional ferrocene unit, or compound **29**, which targets mitochondria without incorporating a functional ferrocene unit. This lack of a protective effect from the inhibitors against the effects of compounds **27** and **29** suggests the specificity of the caspase-mediated pyroptosis pathway for compounds equipped with a functional ferrocene unit. Our observations suggest a nuanced relationship between the molecule localization and their ability to modulate cell death pathways, emphasizing the potential for targeted therapeutic strategies in the modulation of pyroptosis.

## DISCUSSION & CONCLUSION

The synthesis of new fluorophores with multiple functionalities has become an essential aspect in bioimaging and biomedical research. Traditional synthesis methods of such fluorophores rely on linear synthesis routes, which require extensive optimization of reaction conditions before a functional dye can be obtained. In previous work^[22]^, we developed ‘linker’ molecules that can serve as a scaffold for connecting a biological target, a fluorophore and a functional group. Such linkers provide a new means for biolabeling, facilitating the attachment of the linker, followed by the incorporation of the fluorophore in a second step. The synthesis of these linkers was done using a scaffold-based synthesis using the U-4CR.

We here considered the possibility of directly integrating the fluorophore as a moiety into a functional scaffold. Based on the strategy, we obtained fluorogenic probes combining three distinct moeities, i.e., a fluorophore, a signal modulator and a unit for labelling of specific cellular targets. Our approach uses the redox properties of Fc and the resulting fluorescence signal modulation for turn-on fluorescence following Fc oxidation. Besides specific cellular localization we found that our probes are involved in a Fenton-like reaction which can activate pyroptosis, a form of programmed cell death.

The modular synthesis approach used here can simplify the production of multifunctional dyes, not only with fluorogenic character, from a common scaffold. We render it feasible to extend the range of functionalities to self-blinking,^[47]^ antifading (self-healing dyes),^[48]^ metal sensing^[49]^ and many others^[50]^. This approach will thus facilitate the exploration of (bio)molecular imaging and therapeutic applications for imaging, biomedicine and beyond.

## ACKNOWLEGDEMENTS

This work was funded by the National Natural Science Foundation of China (Grant Nos. 22175090 to S.J. and 22374075 to L.Z.), the Primary Research & Development Plan of the Jiangsu Province (BE2021712 to S.J.), China Postdoctoral Science Foundation (2022M721408 to H.Y.W.), and the Jiangsu Provincial Excellent Postdoctoral Program (H.Y.W.). Fluorophore development in the lab of T.C. is supported via the German Science Foundation (Sachbeihilfe DFG/CO879-6-1). We thank Menghui Jia from the Materials Characterization Center and East China Normal University for support of TAS measurements and data analysis.

## Supporting information for Li et al

### Part I: Materials, methods and synthesis

#### Materials

All reagents were purchased from commercial suppliers and used without further purification, unless stated otherwise. Tetrahydro-2H-pyran-2-carboxylic acid (97%, Bidepharm), 7-Bromoheptanoic acid (97%, Bidepharm), paraformaldehyde (95%, Sigma Aldrich), ferrocenecarboxaldehyde (97%, Bidepharm), 5-Aminofluorescein (isomer I) (>95%, TCI), 5-Carboxyfluorescein (97%, TCI), cyclohexyl isocyanide (>98%, TCI), *n*-Butyl isocyanide (98%, Bidepharm), triphenylphosphine (≥99%, Macklin), octadecanal (95%, TCI), 1-heptadecanamine (95%, Energy Chemical), 5-Carboxytetramethylrhodamine (95%, Sigma Aldrich), 4,4,4-triphenylbutan-1-amine, morpholine (≥99%, Aladdin), 2-Morpholinoethanamine, (98%, Aladdin), potassium iodide (≥99.5%, Macklin), methanol (≥99.9%, Macklin), acetonitrile (≥99.9%, Macklin), dichloromethane (DCM) (≥99.5%, Energy Chemical), ethyl acetate (≥99.5%, Energy Chemical), and petroleumether (≥99.5%, Energy Chemical), were used as received.

^1^H and ^13^C NMR spectra were recorded on Bruker DRX-400 (400 MHz) and are reported as follows: chemical shift *δ* (ppm) (multiplicity, number of protons). DMSO was used as solvent. TMS was used as internal standard. Chemical shifts are reported in ppm to the nearest 0.01 ppm for both ^1^H and ^13^C measurements.

High resolution mass spectra were recorded on a Waters ACQUITY UPLC/XEVO G2-XS Q-TOF Spectrometer.

#### Synthesis of compound 14

Compound **1** (0.015 g, 0.5 mmol) and compound **4** (0.174 g, 0.5 mmol) were dissolved in methanol (10.0 mL) before compound **8** (0.065 g, 0.5 mmol) and compound **12** (0.055 g, 0.5 mmol) were added. The solution was stirred for 48 hours. This mixture was concentrated in vacuum to give the crude product, crude **14** was purified by column chromatography (eluent: dichloromethane, methanol (50:1, 20:1 then 10:1)) yielding **14** (0.276 g, 92 %).

^1^H NMR (400 MHz, DMSO-*d*_6_) δ 10.07 (s, 2H), 8.40 (s, 1H), 8.0 (d, 1H), 7.18 (d, 1H), 6.66 (m, 2H), 6.58 (m, 5H), 4.21-3.97 (m, 4H), 3.69-3.54 (m, 2H), 1.95-1.23 (m,16H) ppm.

^13^C NMR (101 MHz, DMSO-*d_6_*) δ 172.74, 171.52, 169.09, 159.93, 152.36, 148.16, 140.54, 129.55, 127.81, 126.86, 124.67, 114.97, 113.62, 110.92, 103.34, 83.50, 78.44, 75.57, 69.17, 67.10, 31.14, 29.04, 28.11, 25.55, 23.29, 22.76 ppm.

HRMS [M(**14**+H)]^+^ calculated for C_34_H_35_N_2_O_8_ 599.2388 u, found 599.2405 u.

#### Synthesis of compound 15

Compound **2** (0.214 g, 1.0 mmol) and compound **4** (0.347 g, 1.0 mmol) were dissolved in methanol (10.0 mL) before compound **8** (0.130 g, 1.0 mmol) and compound **12** (0.109 g, 1.0 mmol) were added. The solution was stirred for 48 hours. This mixture was concentrated in vacuum to give the crude product, crude **15** was purified by column chromatography (eluent: dichloromethane, methanol (50:1, 30:1 then 15:1)) yielding **15** (0.588 g, 75 %).

^1^HNMR (400 MHz, DMSO-*d*_6_) δ 10.13 (s, 2H), 8.36-7.88 (m, 2H), 7.31-6.19 (m, 8H), 5.72 (s, 1H), 4.19-3.65 (m, 13H), 1.90-1.33 (m,16H) ppm.

^13^C NMR (101 MHz, DMSO-*d_6_*) δ 168.95, 168.82, 168.29, 160.04, 152.43, 151.13, 140.83, 129.31, 123.39, 113.00, 102.72, 83.57, 82.20, 75.47, 70.08, 69.27, 67.76, 59.93, 48.66, 32.82, 27.50, 25.77, 25.28, 22.27 ppm.

LCMS [M(**15**+H)]^+^ calculated for C_44_H_43_FeN_2_O_8_ 783.2364 u, found 783.19 u.

#### Synthesis of compound 16

Compound **4** (0.347 g, 1.0 mmol) and compound **1** (0.030 g, 1.0 mmol) were dissolved in methanol (5.0 mL) before compound **9** (0.209 g, 1.0 mmol) and compound **13** (0.083 g, 1.0 mmol) were added. The solution was stirred at 50°C for 48 hours. This mixture was concentrated in vacuum to give the crude product, crude **16** was purified by column chromatography (eluent: dichloromethane, methanol (50:1, 25:1 then 12:1)) yielding **16** (0.478 g, 73.5 %).

^1^H NMR (400 MHz, DMSO-*d*_6_): δ 10.17 (s, 2H), 7.99 (s, 1H), 7.78 (s, 1H), 7.61 (s, 1H), 7.32 (s, 1H), 6.70 (m, 2H), 6.57 (m, 4H), 4.22 (s, 2H), 3.52 (m, 2H), 2.19 (m, 2H), 1.74 (m, 2H), 1.49 (m, 2H), 1.23 (m, 13H) ppm.

^13^C NMR (101 MHz, DMSO-*d_6_*) δ 174.92, 168.51, 167.77, 160.06, 152.38, 150.02, 145.25, 136.16, 129.43, 127.91, 125.64, 124.51, 113.05, 109.83, 102.78, 83.75, 50.74, 49.07, 35.51, 33.99, 28.93, 28.08, 27.70, 24.76 ppm.

LCMS [M(**16**+H)]^+^ calculated for C_33_H_36_BrN_2_O_7_ 651.1701 u, found 651.0434 u. HRMS [M(**16**+H)]^+^ calculated for C_33_H_36_BrN_2_O_7_ 651.1701 u, found 651.1695 u.

#### Synthesis of compound 17

Compound **4** (0.695 g, 2.0 mmol) and compound **2** (0.428 g, 2.0 mmol) were dissolved in methanol (10.0 mL) before compound **9** (0.418 g, 2.0 mmol) and compound **13** (0.166 g, 2.0 mmol) were added. The solution was stirred at 50 °C for 48 hours, and then ethyl acetate (40 mL) was added. Subsequently, the organic layer was washed with water (3 x 20 mL). The combined water layers were extracted with ethyl acetate (2 x 20 mL). The organic fractions were then combined, dried over magnesium sulfate, filtered, and concentrated under reduced pressure, yielding compound **17** as a brown-yellow powder. Yield: 1.179 g (72%).

^1^H NMR (400 MHz, DMSO-*d*_6_) δ 10.14 (s, 2H), 8.42 (s, 1H), 6.99-7.38 (m, 2H), 6.55-6.67 (t, 3H), 6.12 (s, 1H), 5.77 (s, 1H), 4.22 (s, 5H), 3.95-4.15 (d, 4H), 3.48 (s, 2H), 2.00 (s, 2H), 1.75 (s, 2H), 1.20-1.44 (m, 15H) ppm.

^13^C NMR (101 MHz, DMSO-*d_6_*) δ 170.96, 168.97, 167.87, 159.83, 152.33, 152.05, 150.57, 141.79, 138.49, 129.26, 128.83, 126.84, 123.75, 112.63, 102.42, 82.50, 69.82, 68.99, 67.64, 67.21, 60.05, 50.59, 45.43, 35.21, 34.36, 32.86, 32.20, 28.67, 27.30, 26.06, 24.76 ppm.

LCMS [M(**17**+H)]^+^ calculated for C_43_H_44_BrFeN_2_O_7_ 835.1676 u, found 835.1088 u. HRMS [M(**17**+H)]^+^ calculated for C_43_H_43_BrFeN_2_O_7_ 835.1676 u, found 835.1666 u.

#### Synthesis of compound 18

Compound **1** (0.03 g, 1.0 mmol) and compound **5** (0.130 g, 1.0 mmol) were dissolved in methanol (10.0 mL) before compound **10** (0.376 g, 1.0 mmol) and compound **13** (0.083 g, 1.0 mmol) were added. The solution was stirred for 48 hours. This mixture was concentrated in vacuum to give the crude product, crude **18** was purified by preparative HPLC (eluent, a 30 min linear gradient, from 10% to 60% solvent B; flow rate, 19.0 mL/min, eluent A (ddH_2_O with 0.1% formic acid) and eluent B (CH_3_CN with 0.1% formic acid)) and lyophilized to obtain **18**: *t*_R_ = 20.6 min, (0.416 g, 69.2%).

LCMS [M(**18**+H)]^+^ calculated for C_33_H_36_N_3_O_8_ 602.2497 u, found 602.09 u.

#### Synthesis of compound 19

Compound **2** (0.214 g, 1.0 mmol) and compound **5** (0.130 g, 1.0 mmol) were dissolved in methanol (10.0 mL) before compound **10** (0.376 g, 1.0 mmol) and compound **13** (0.083 g, 1.0 mmol) were added. The solution was stirred for 48 hours. This mixture was concentrated in vacuum to give the crude product, crude **19** was purified by purified by preparative HPLC (eluent, a 30 min linear gradient, from 10% to 60% solvent B; flow rate, 19.0 mL/min, eluent A (ddH_2_O with 0.1% formic acid) and eluent B (CH_3_CN with 0.1% formic acid)) and lyophilized to obtain **19**: *t*_R_ = 24.4 min, (0.642 g, 81.8%).

LCMS [M(**19**+H)]^+^ calculated for C_43_H_44_FeN_3_O_8_ 786.2473 u, found 786.07 u.

#### Synthesis of compound 20

Compound **1** (0.03 g, 1.0 mmol) and compound **5** (0.130 g, 1.0 mmol) were dissolved in methanol (10.0 mL) before compound **11** (0.431 g, 1.0 mmol) and compound **13** (0.083 g, 1.0 mmol) were added. The solution was stirred for 48 hours. This mixture was concentrated in vacuum to give the crude product, crude **20** was purified by preparative HPLC (eluent, a 30 min linear gradient, from 10% to 60% solvent B; flow rate, 19.0 mL/min, eluent A (ddH_2_O with 0.1% formic acid) and eluent B (CH_3_CN with 0.1% formic acid)) and lyophilized to obtain **20**: *t*_R_ = 15.4 min, (0.402 g, 61.1%).

LCMS [M(**20**)] calculated for C_37_H_47_N_5_O_6_ 657.3526 u, found 656.17 u.

#### Synthesis of compound 21

Compound **2** (0.214 g, 1.0 mmol) and compound **5** (0.130 g, 1.0 mmol) were dissolved in methanol (10.0 mL) before compound **11** (0.431 g, 1.0 mmol) and compound **13** (0.083 g, 1.0 mmol) were added. The solution was stirred for 48 hours. This mixture was concentrated in vacuum to give the crude product, crude **21** was purified by preparative HPLC (eluent, a 30 min linear gradient, from 10% to 60% solvent B; flow rate, 19.0 mL/min, eluent A (ddH_2_O with 0.1% formic acid) and eluent B (CH_3_CN with 0.1% formic acid)) and lyophilized to obtain **21**: *t*_R_ = 19.7 min, (0.517 g, 61.4%).

LCMS [M(**21**)] calculated for C_47_H_55_N_5_O_6_ 841.3502 u, found 840.29 u.

#### Synthesis of compound 22

Compound **1** (0.03 g, 1.0 mmol) and compound **6** (0.320 g, 1.0 mmol) were dissolved in methanol (10.0 mL) before compound **10** (0.376 g, 1.0 mmol) and compound **13** (0.083 g, 1.0 mmol) were added. The solution was stirred for 48 hours. This mixture was concentrated in vacuum to give the crude product, crude **22** was purified by preparative HPLC (eluent, a 30 min linear gradient, from 10% to 60% solvent B; flow rate, 19.0 mL/min, eluent A (ddH_2_O with 0.1% formic acid) and eluent B (CH_3_CN with 0.1% formic acid)) and lyophilized to obtain **22**: *t*_R_ = 15.6 min, (0.348 g, 43.9%).

LCMS [M(**22**)] calculated for C_48_H_44_N_2_O_7_P^+^ 791.2881 u, found 791.15 u.

#### Synthesis of compound 23

Compound **2** (0.214 g, 1.0 mmol) and compound **6** (0.320 g, 1.0 mmol) were dissolved in methanol (10.0 mL) before compound **10** (0.376 g, 1.0 mmol) and compound **13** (0.083 g, 1.0 mmol) were added. The solution was stirred for 48 hours. This mixture was concentrated in vacuum to give the crude product, crude **23** was purified by preparative HPLC (eluent, a 30 min linear gradient, from 10% to 60% solvent B; flow rate, 19.0 mL/min, eluent A (ddH_2_O with 0.1% formic acid) and eluent B (CH_3_CN with 0.1% formic acid)) and lyophilized to obtain **23**: *t*_R_ = 21.4 min, (0.653 g, 66.9%).

LCMS [M(**23**)] calculated for C_58_H_52_FeN_2_O_7_P^+^ 975.2856 u, found 975.29 u.

#### Synthesis of compound 24

Compound **1** (0.03 g, 1.0 mmol) and compound **6** (0.320 g, 1.0 mmol) were dissolved in methanol (10.0 mL) before compound **11** (0.431 g, 1.0 mmol) and compound **13** (0.083 g, 1.0 mmol) were added. The solution was stirred for 48 hours. This mixture was concentrated in vacuum to give the crude product, crude **24** was purified by preparative HPLC (eluent, a 30 min linear gradient, from 10% to 60% solvent B; flow rate, 19.0 mL/min, eluent A (ddH_2_O with 0.1% formic acid) and eluent B (CH_3_CN with 0.1% formic acid)) and lyophilized to obtain **24**: *t*_R_ = 21.7 min, (0.549 g, 64.9%).

LCMS [M(**24**)] calculated for C_52_H_54_N_4_O_5_P^+^ 845.3826 u, found 845.36 u.

#### Synthesis of compound 25

Compound **2** (0.214 g, 1.0 mmol) and compound **6** (0.320 g, 1.0 mmol) were dissolved in methanol (10.0 mL) before compound **11** (0.431 g, 1.0 mmol) and compound **13** (0.083 g, 1.0 mmol) were added. The solution was stirred for 48 hours. This mixture was concentrated in vacuum to give the crude product, crude **25** was purified by preparative HPLC (eluent, a 30 min linear gradient, from 10% to 60% solvent B; flow rate, 19.0 mL/min, eluent A (ddH_2_O with 0.1% formic acid) and eluent B (CH_3_CN with 0.1% formic acid)) and lyophilized to obtain **25**: *t*_R_ = 22.7 min, (0.760 g, 73.8%).

LCMS [M(**25**)] calculated for C_62_H_62_FeN_4_O_5_P^+^ 1029.3802 u, found 1029.3767 u.

#### Synthesis of compound 26

Compound **3** (0.268 g, 1.0 mmol) and compound **7** (0.255 g, 1.0 mmol) were dissolved in methanol (10.0 mL) before compound **11** (0.431 g, 1.0 mmol) and compound **13** (0.083 g, 1.0 mmol) were added. The solution was stirred for 48 hours. The crude product **26** precipitated in the form of a solid. It was then sequentially washed three times with water and methanol. Following these washings, the product was dried, yielding the final compound **26**, 0.692 g, 67.9%.

HRMS [M(**26**)] calculated for C_65_H_104_N_4_O_5_ 1020.8007 u, found 1020.7988 u.

^1^H NMR (400 MHz, CDCl_3_): δ 8.12 (s, 1H), 7.75-7.37 (m, 7H), 7.14-7.09 (m, 2H), 5.39-5.37 (m, 2H), 4.33-3.94 (m, 8H), 2.07-2.00 (m, 2H), 1.76-1.71 (m, 2H), 1.40 (s, 9H), 1.36-1.26 (m, 58H), 0.99-0.90 (m, 12H) ppm.

#### Synthesis of compound 27

Compound **16** (0.110 g, 0.169 mmol) and morpholine (0.33 mL, 3.79 mmol) were refluxed in ethanol (7.0 mL) for 24 hours. The mixture was cooled to room temperature and concentrated in vacuum to obtain a crude product. crude **27** was purified by column chromatography (eluent: dichloromethane, methanol (20:1, 10:1 then 6:1)) yielding **27** (0.082 g, 73.7 %).

^1^H NMR (400 MHz, DMSO-*d*_6_): δ 7.96-7.30 (m, 4H), 6.71-6.56 (m, 5H), 4.23 (s, 2H), 3.59 (m, 4H), 3.40 (m, 4H), 2.79-1.88 (m, 6H), 1.47-1.34 (m, 2H), 1.23 (s, 9H) ppm.

^13^C NMR (101 MHz, DMSO-*d_6_*) δ 170.05, 168.95, 167.83, 161.73, 153.00, 150.82, 137.97, 129.60, 125.20, 121.31, 117.25, 113.84, 111.25, 107.55, 102.98, 81.39, 66.96, 66.63, 58.70, 53.81, 52.05, 50.68, 49.02, 45.31, 28.91, 27.06, 26.23 ppm.

LCMS [M(**27**+H)]^+^ calculated for C_37_H_44_N_3_O_8_ 658.3123 u, found 658.09 u. HRMS [M(**27**+H)]^+^ calculated for C_37_H_44_N_3_O_8_ 658.3123 u, found 658.3121 u.

#### Synthesis of compound 28

Compound **17** (0.050 g, 0.06 mmol) and morpholine (0.25 mL, 2.87 mmol) were refluxed in ethanol (5.0 mL) for 24 hours. The mixture was cooled to room temperature and concentrated in vacuum to obtain a crude product. crude **28** was purified by column chromatography (eluent: dichloromethane, methanol (25:1, 15:1 then 8:1)) yielding **28** (0.035 g, 69.3 %).

^1^H NMR (400 MHz, CDCl_3_): δ 7.91-7.32 (m, 2H), 7.08-6.86 (m, 2H), 6.61-6.40 (m, 6H), 5.83 (s, 1H), 4.11-3.88 (m, 9H), 3.61 (m, 4H), 3.16-3.03 (m, 2H), 2.52-1.98 (m, 8H), 1.67-1.46 (m, 15H).

^13^C NMR (101 MHz, DMSO-*d_6_*): δ 170.94, 168.77, 161.01, 156.69, 153.30, 151.63, 141.24, 139.04, 129.63, 129.25, 129.19, 128.28, 114.79, 112.59, 110.52, 109.58, 102.39, 102.10, 82.17, 69.05, 68.80, 67.46, 67.11, 66.21, 65.79, 58.62, 58.25, 53.39, 48.60, 45.21, 30.17, 28.48, 26.60, 25.82, 24.89 ppm.

LCMS [M(**28**+H)]^+^ calculated for C_47_H_52_FeN_3_O_8_ 842.3099 u, found 842.20 u. HRMS [M(**28**+H)]^+^ calculated for C_47_H_52_FeN_3_O_8_ 842.3099 u, found 842.3132 u.

#### Synthesis of compound 29

A solution of compound **16** (0.200 g, 0.30 mmol), triphenylphosphine (0.779 g, 3.0 mmol), and sodium iodide (0.329 g, 1.98 mmol) in dry acetonitrile (18 mL) was refluxed under argon for 24 hours. This mixture was cooled to room temperature and concentrated in vacuum to give the crude product, crude **29** was purified by column chromatography (eluent: dichloromethane, methanol (100:1, 50:1 then 25:1)) yielding **29** (0.125 g, 50 %).

^1^H NMR (400 MHz, DMSO-*d*_6_): δ 7.95-7.77 (m, 15H), 7.62 (m, 2H), 7.39-7.25 (m, 2H), 6.64-6.46 (m, 6H), 4.21 (s, 2H), 3.70-3.17 (m, 2H), 2.13 (m, 2H), 1.87 (m, 2H), 1.30-1.20 (m, 15H).

^13^C NMR (101 MHz, DMSO-*d_6_*) δ 171.68, 168.19, 167.41, 160.04, 152.09, 144.73, 131.60, 130.23, 130.22, 128.94, 128.82, 120.32, 119.44, 118.95, 118.09, 112.95, 110.42, 109.53, 102.40, 86.23, 55.14, 48.68, 29.65, 29.48, 28.50, 27.83, 24.37, 21.71, 20.62, 20.12 ppm.

LCMS [M(**29**)] calculated for C_51_H_50_N_2_O_7_P^+^ 833.3350 u, found 832.89 u.

HRMS [M(**29**)] calculated for C_51_H_50_N_2_O_7_P^+^ 833.3350 u, found 833.3374 u.

#### Synthesis of compound 30

A solution of compound **17** (0.206 g, 0.25 mmol), triphenylphosphine (0.649 g, 2.5 mmol), and sodium iodide (0.274 g, 1.65 mmol) in dry acetonitrile (16 mL) was refluxed under argon for 24 hours. This mixture was cooled to room temperature and concentrated in vacuum to give the crude product, crude **30** was purified by column chromatography (eluent: dichloromethane, methanol (50:1, 20:1 then 8:1)) yielding **30** (0.129 g, 50.7 %).

^1^H NMR (400 MHz, DMSO-*d*_6_): δ 10.27 (s, 2H), 8.17-8.02 (m, 2H), 7.90-7.79 (m, 15H), 7.40-6.97 (m, 2H), 6.66-6.52 (m, 5H), 6.11 (s, 1H), 5.73 (s, 1H), 4.21 (s, 5H), 4.09-3.80 (m, 4H), 3.53 (m, 2H), 1.99 (m, 2H), 1.47-1.34 (m, 15H), 1.19 (m, 2H) ppm.

^13^C NMR (101 MHz, DMSO-*d_6_*): δ 170.76, 168.85, 168.10, 152.12, 141.54, 138.12, 134.93, 133.61, 133.51, 130.33, 130.21, 129.16, 128.82, 127.14, 123.86, 118.93, 118.08, 112.90, 109.83, 109.09, 102.31, 82.33, 69.63, 68.84, 67.47, 67.05, 60.03, 50.39, 34.22, 29.72, 29.55, 28.47, 27.47, 24.58, 21.71, 20.42, 19.92 ppm.

LCMS [M(**30**)] calculated for C_61_H_58_FeN_2_O_7_P^+^ 1017.3326 u, found 1017.2709 u.

HRMS [M(**30**)] calculated for C_61_H_58_FeN_2_O_7_P^+^ 1017.3326 u, found 1017.3329 u.

### Part II: Chemical characterization of synthesized compounds

**Figure S1.**
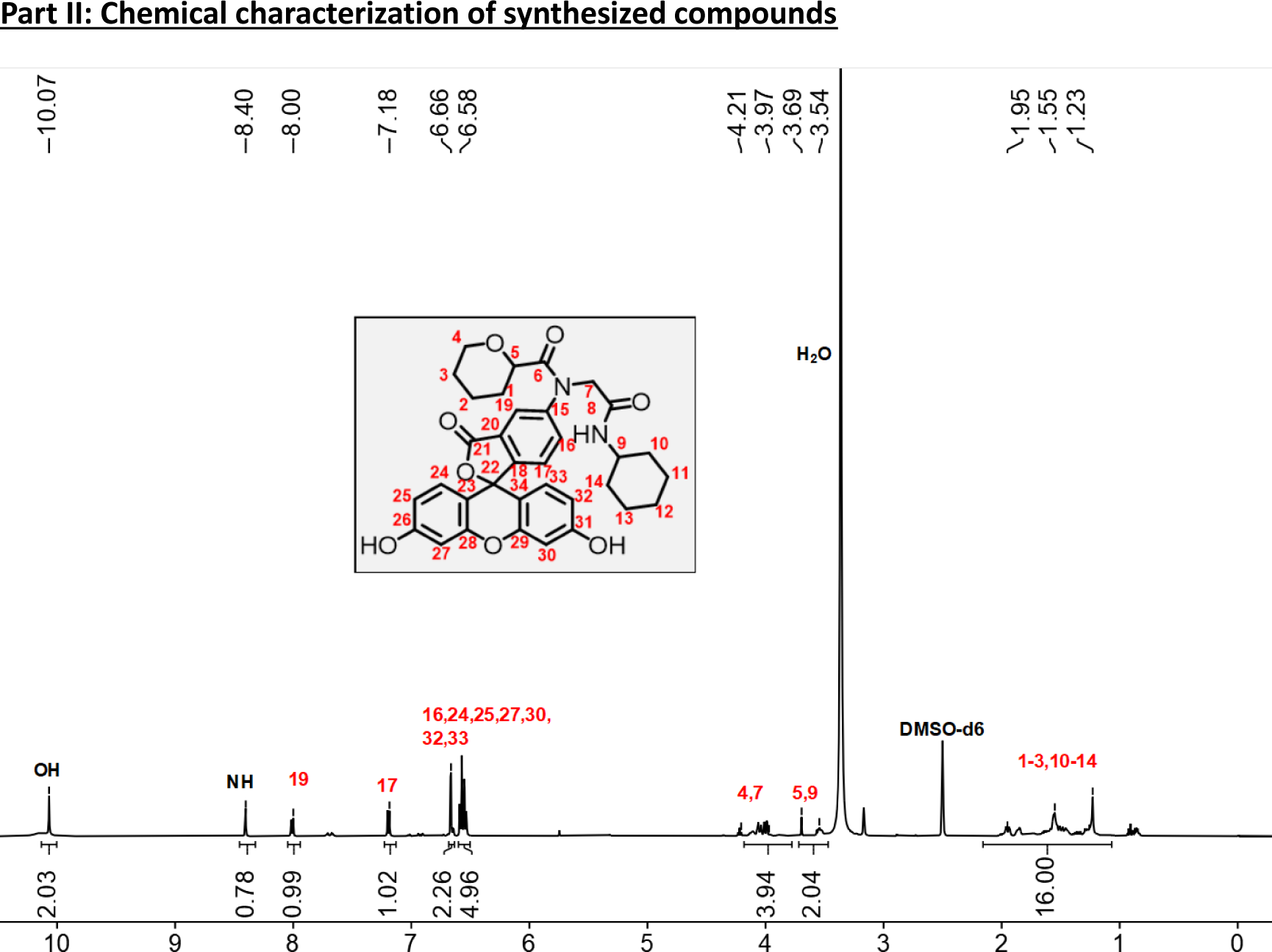
^1^H NMR spectrum of **14**.

**Figure S2.**
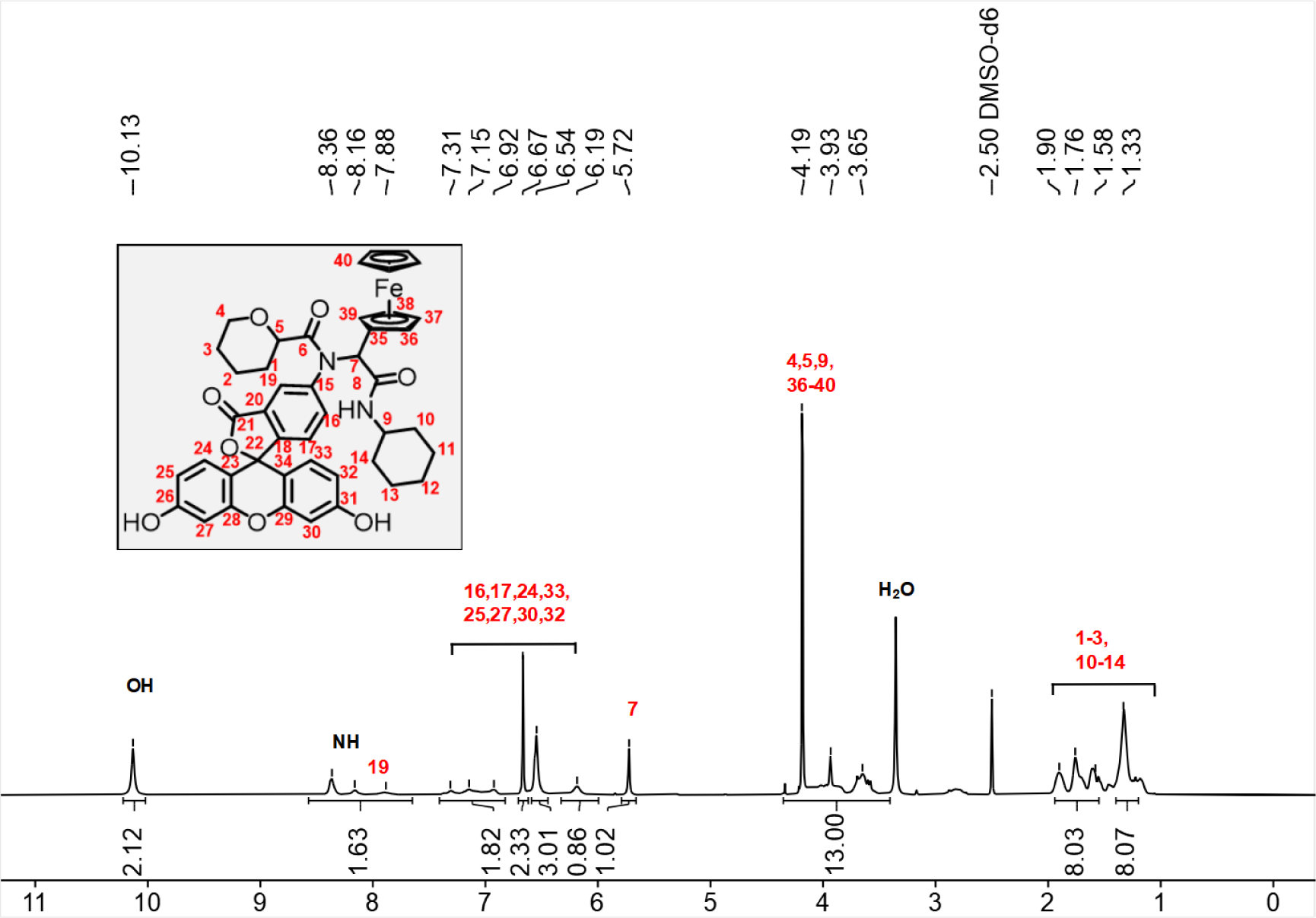
^1^H NMR spectrum of **15**.

**Figure S3.**
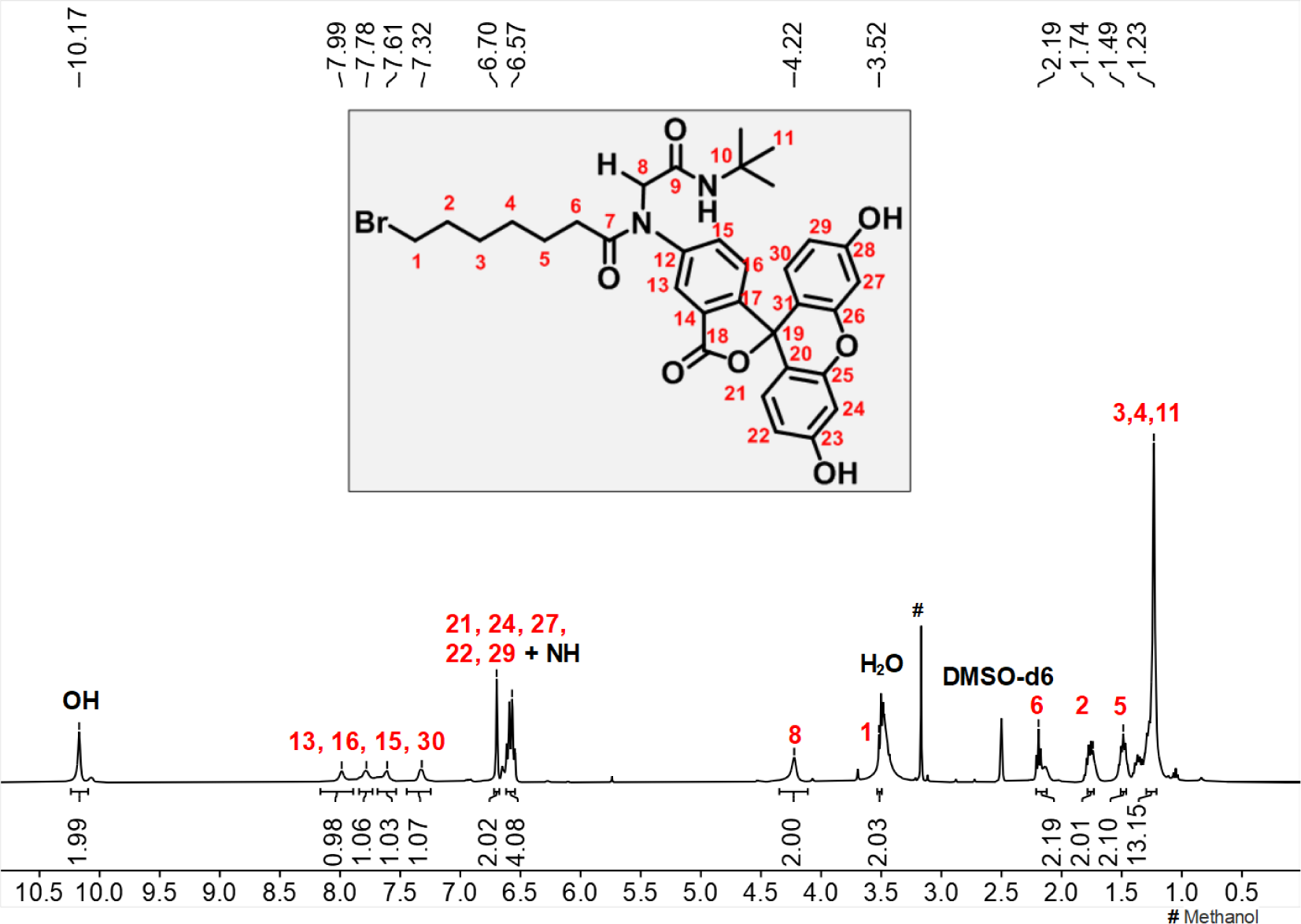
^1^H NMR spectrum of **16**.

**Figure S4.**
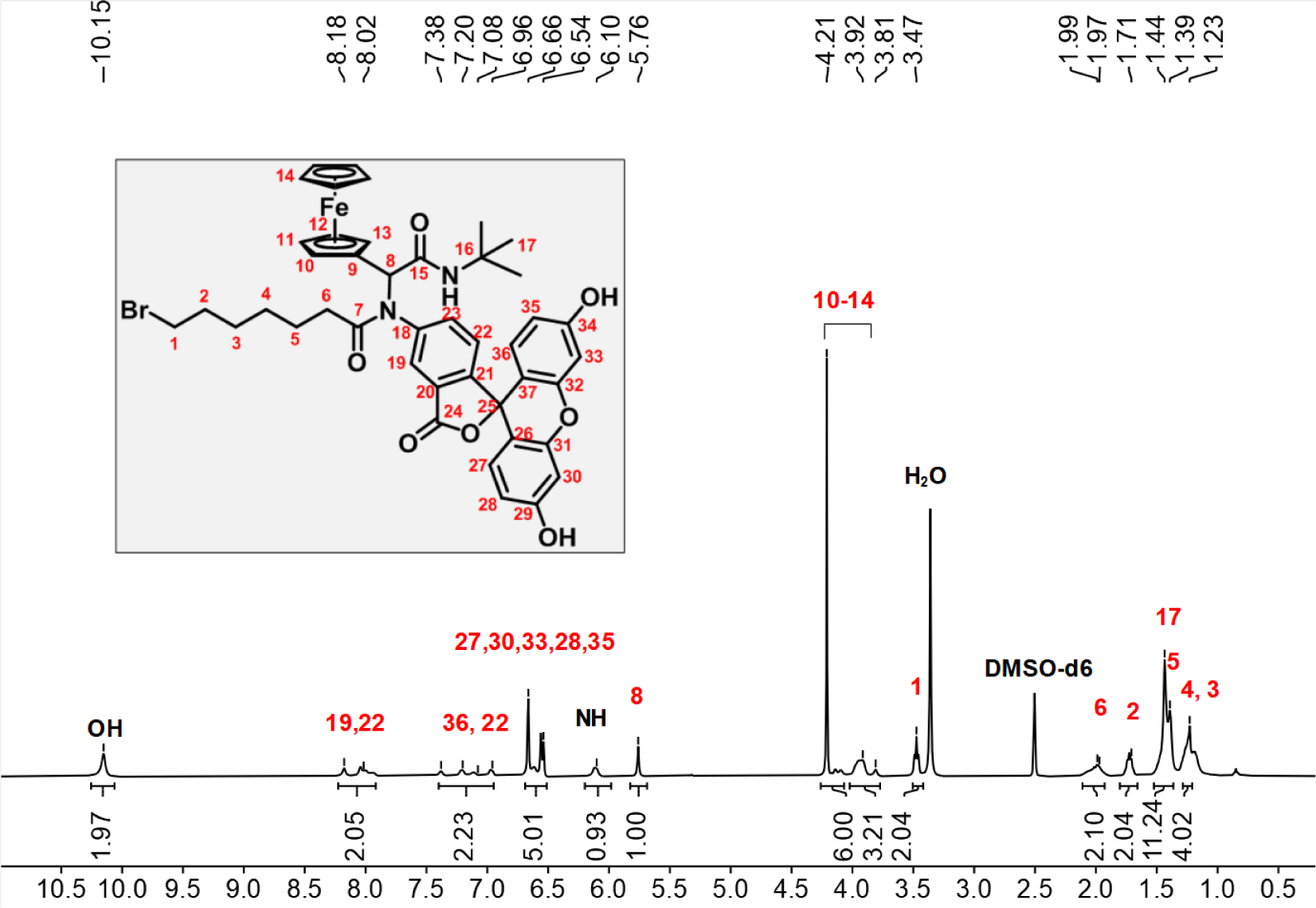
^1^H NMR spectrum of **17**.

**Figure S5.**
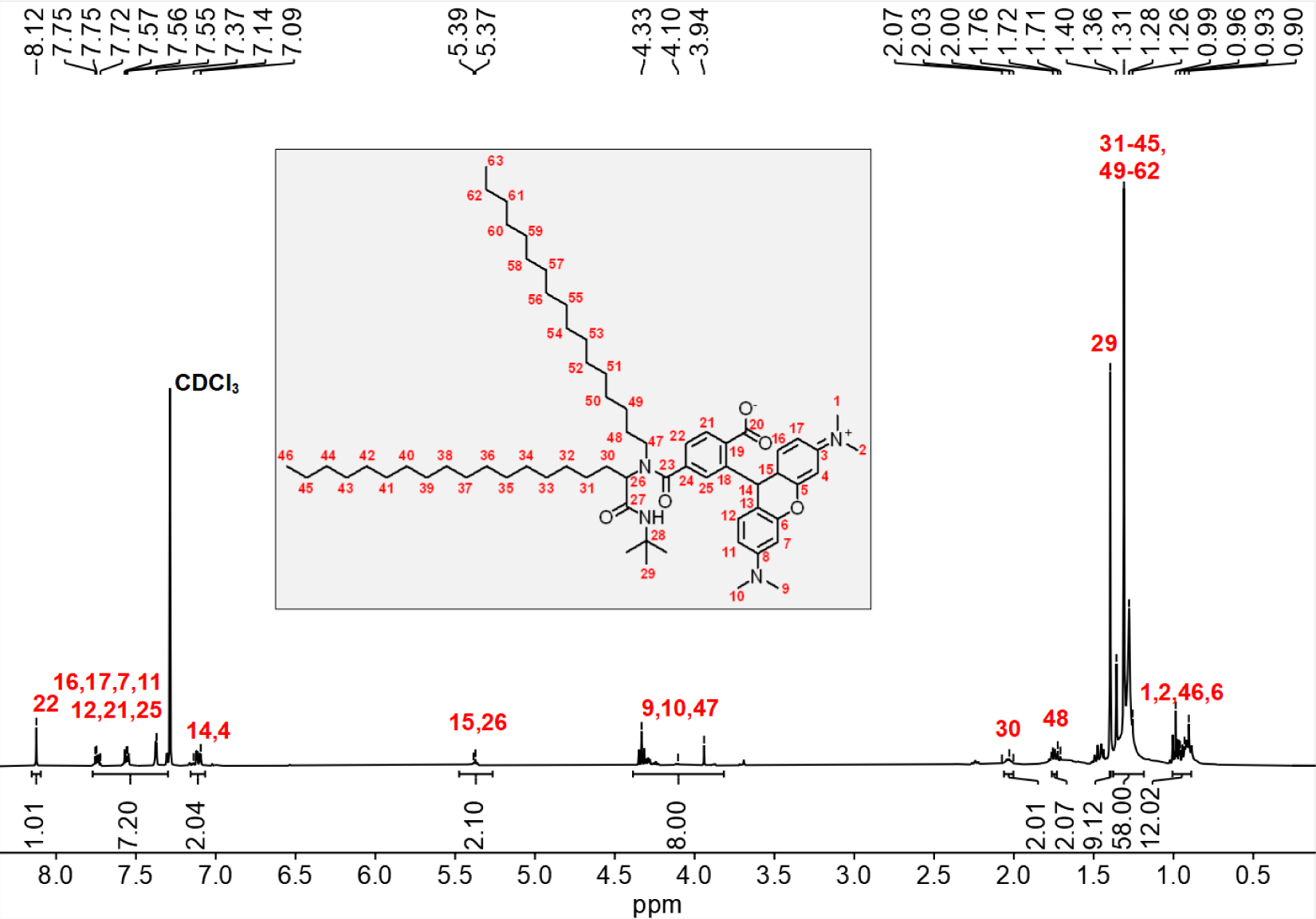
^1^H NMR spectrum of **26**.

**Figure S6.**
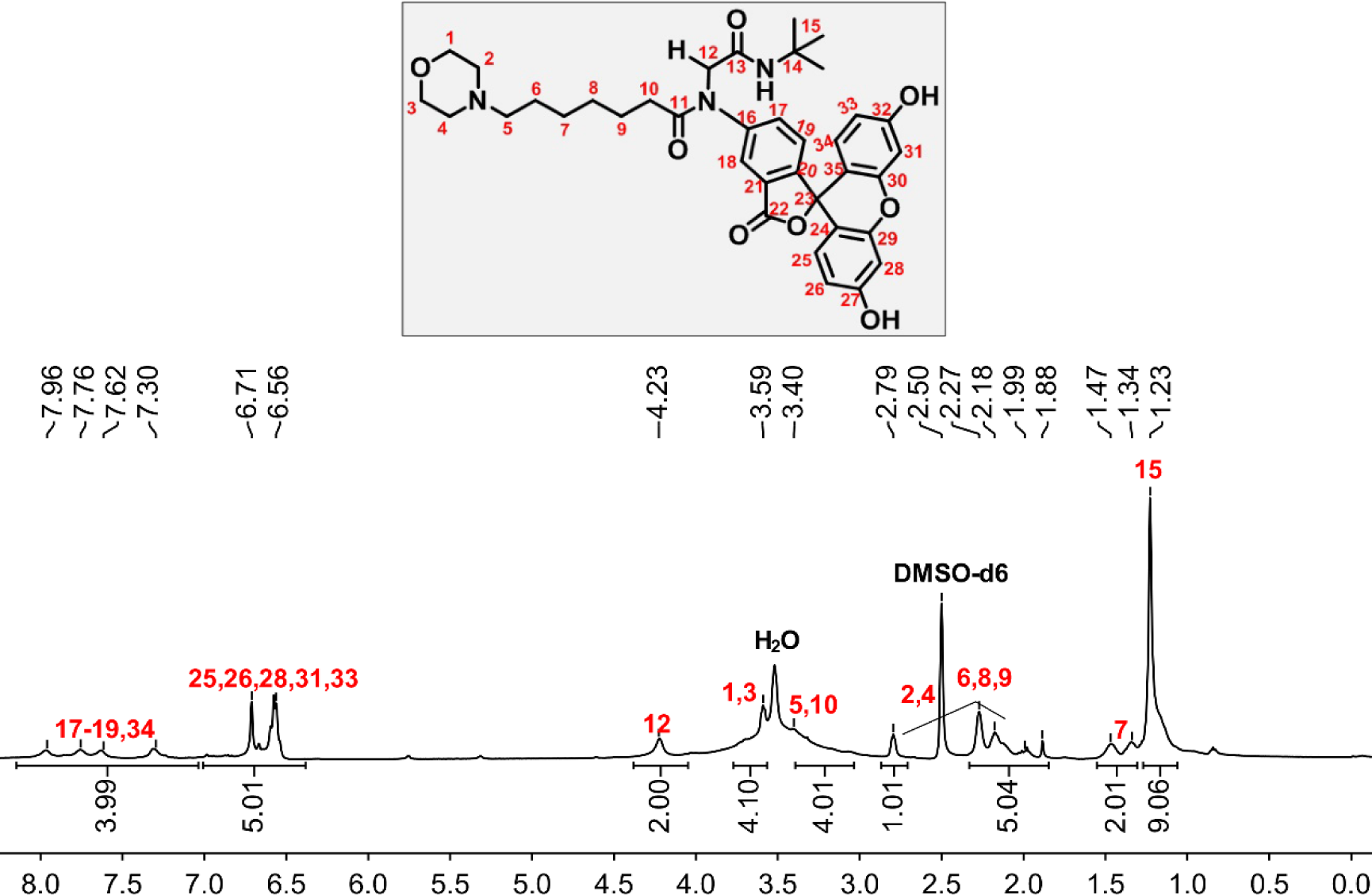
^1^H NMR spectrum of **27**.

**Figure S7.**
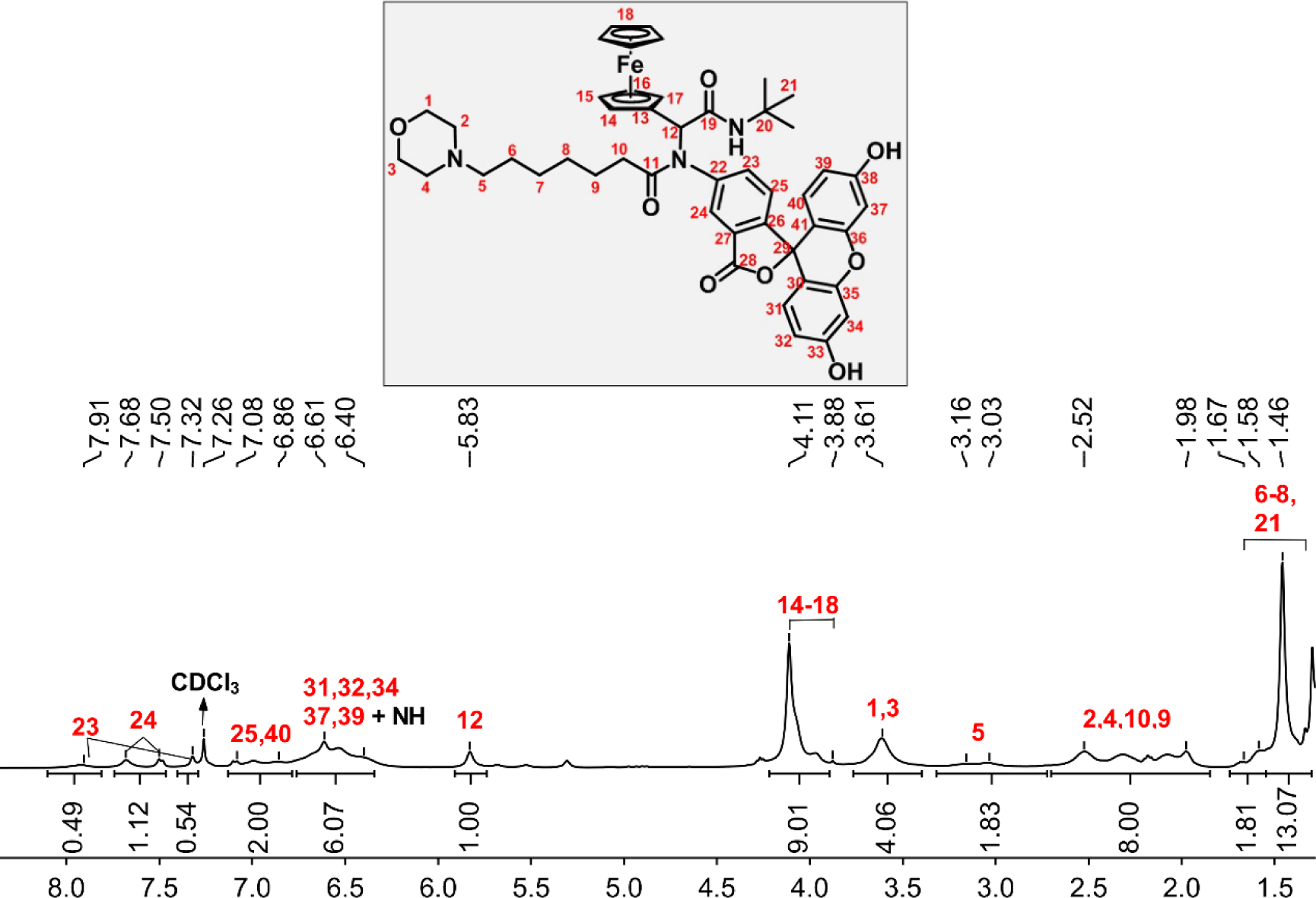
^1^H NMR spectrum of **28**.

**Figure S8.**
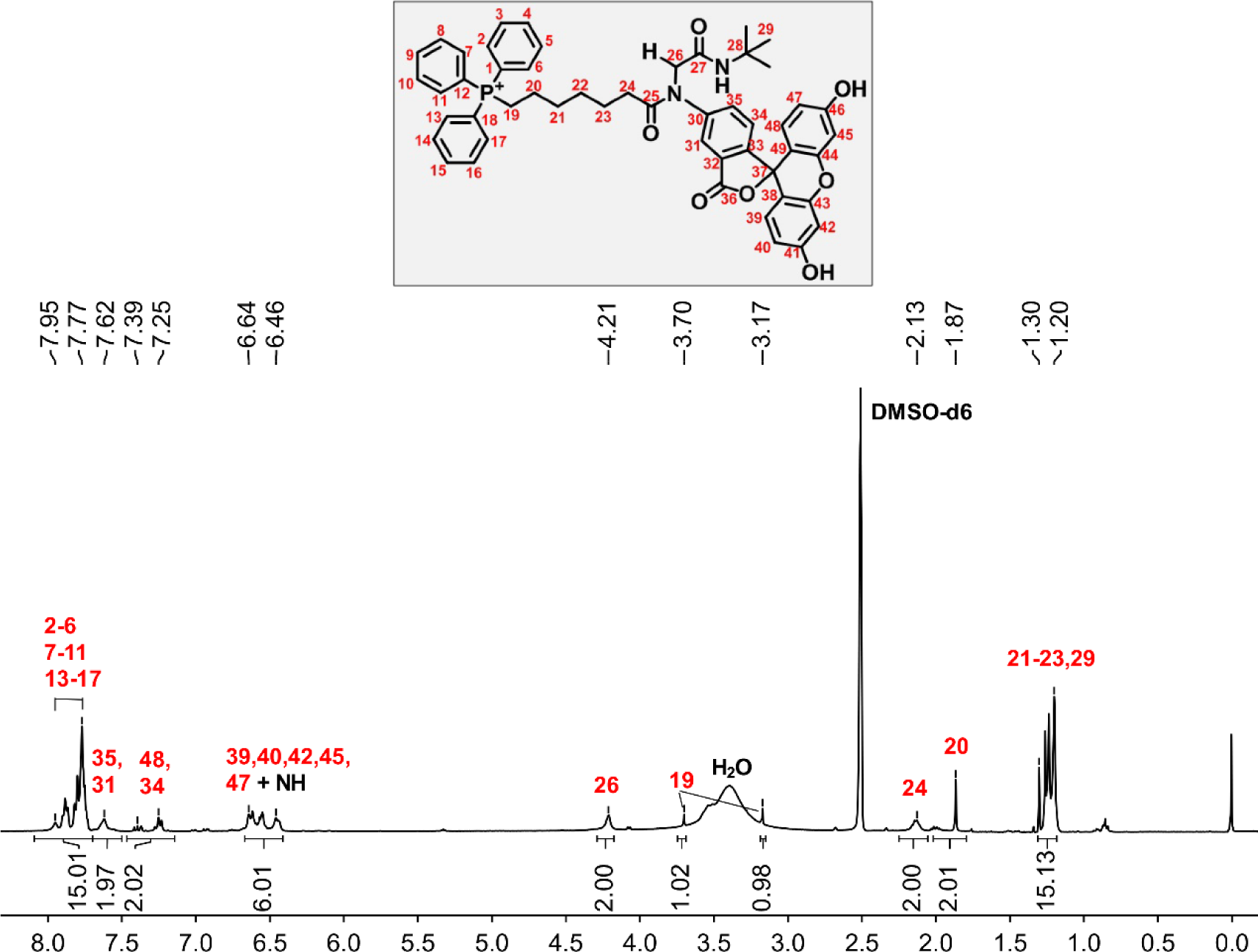
^1^H NMR spectrum of **29**.

**Figure S9.**
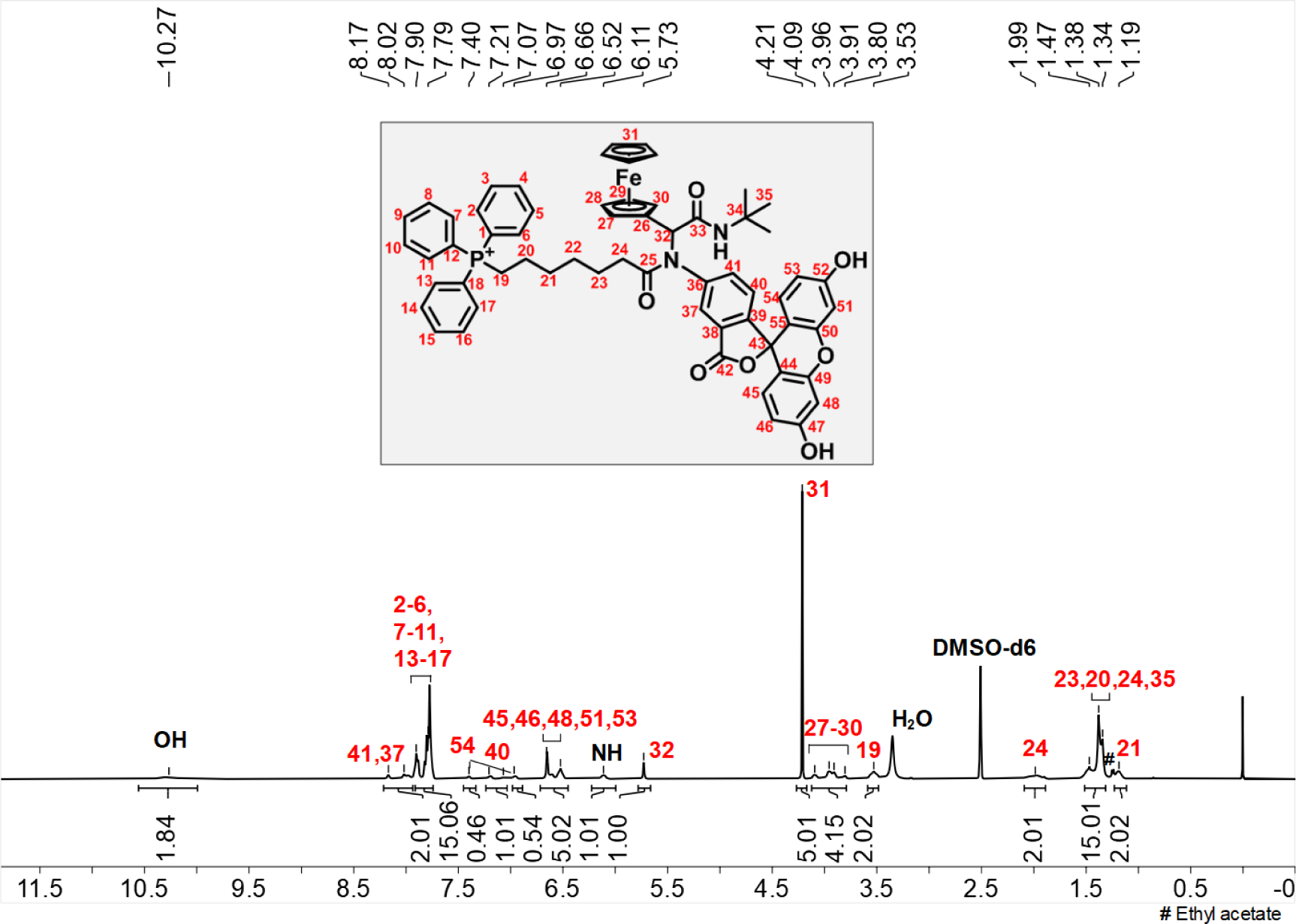
^1^H NMR spectrum of **30**.

**Figure S10.**
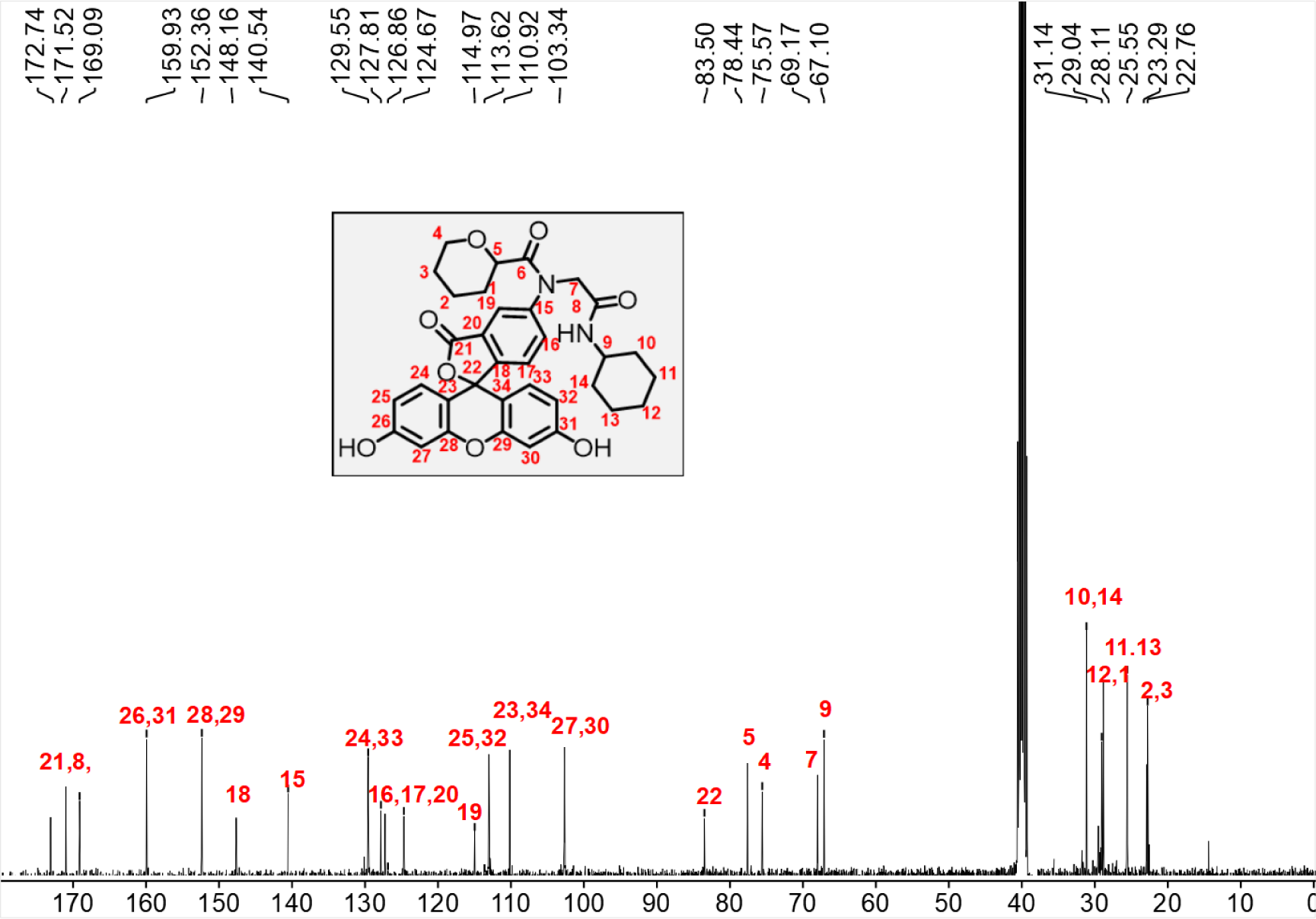
^1^C NMR spectrum of **14**.

**Figure S11.**
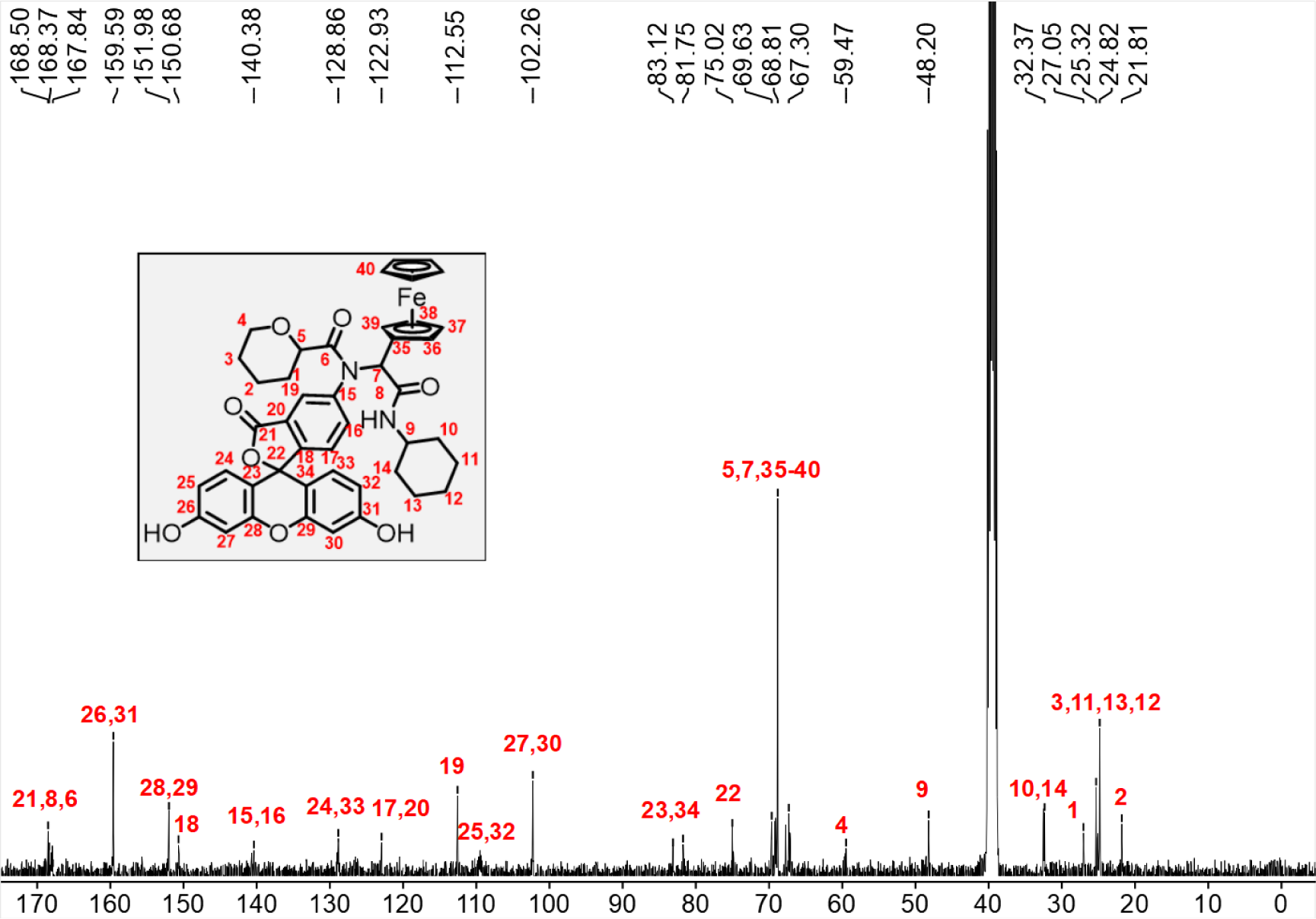
^1^C NMR spectrum of **15**.

**Figure S12.**
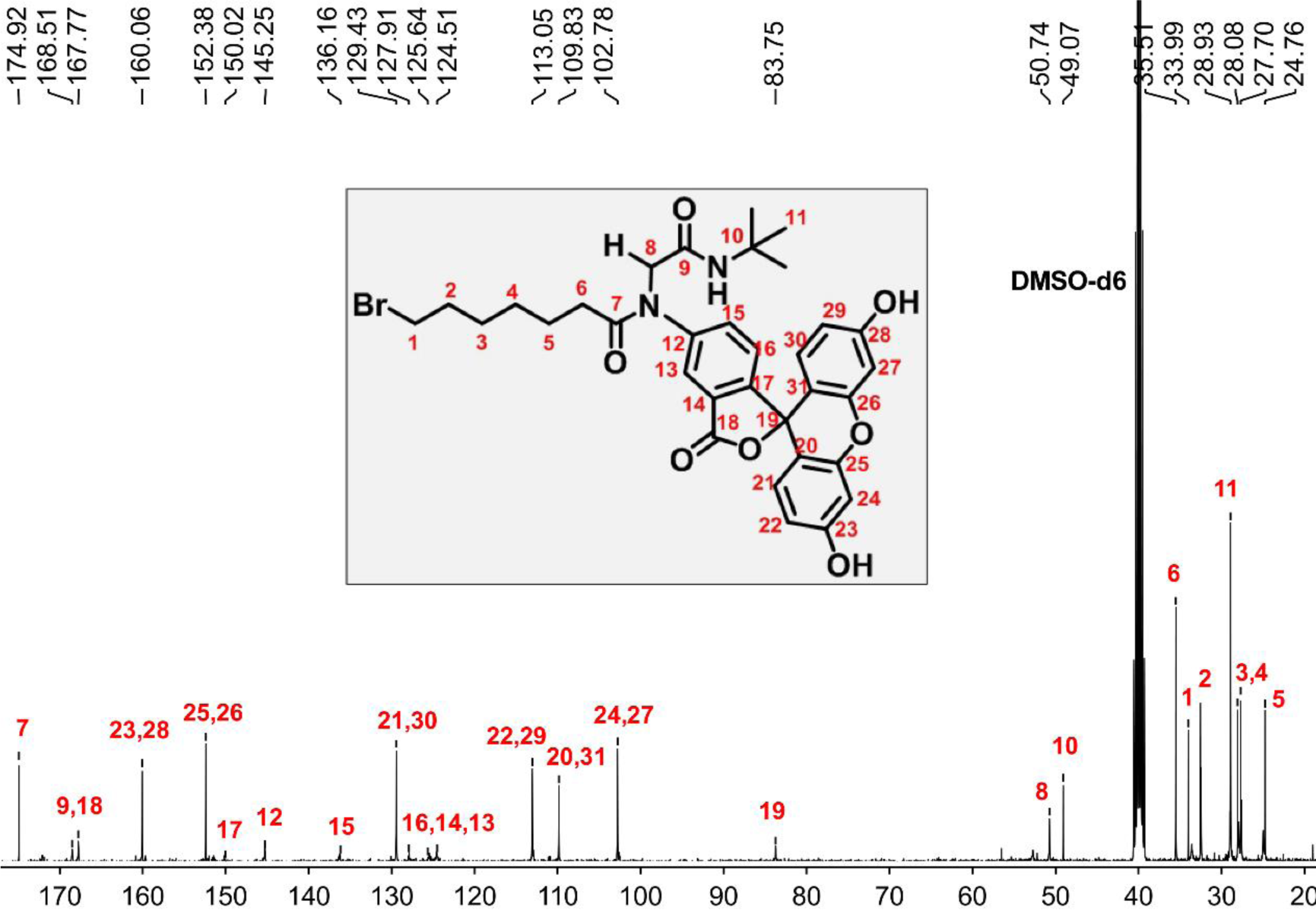
^13^C NMR spectrum of **16**.

**Figure S13.**
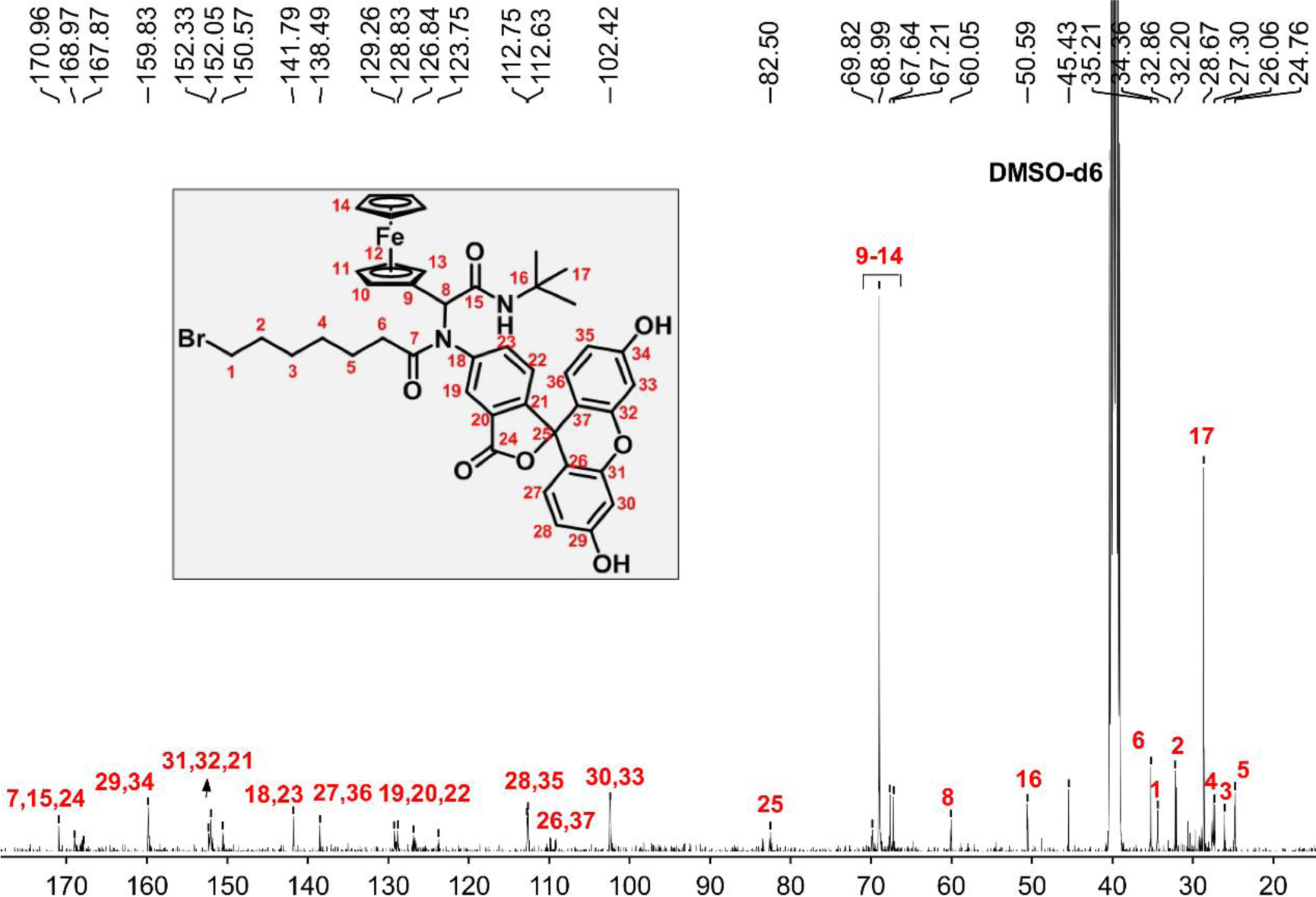
^13^C NMR spectrum of **17**.

**Figure S14.**
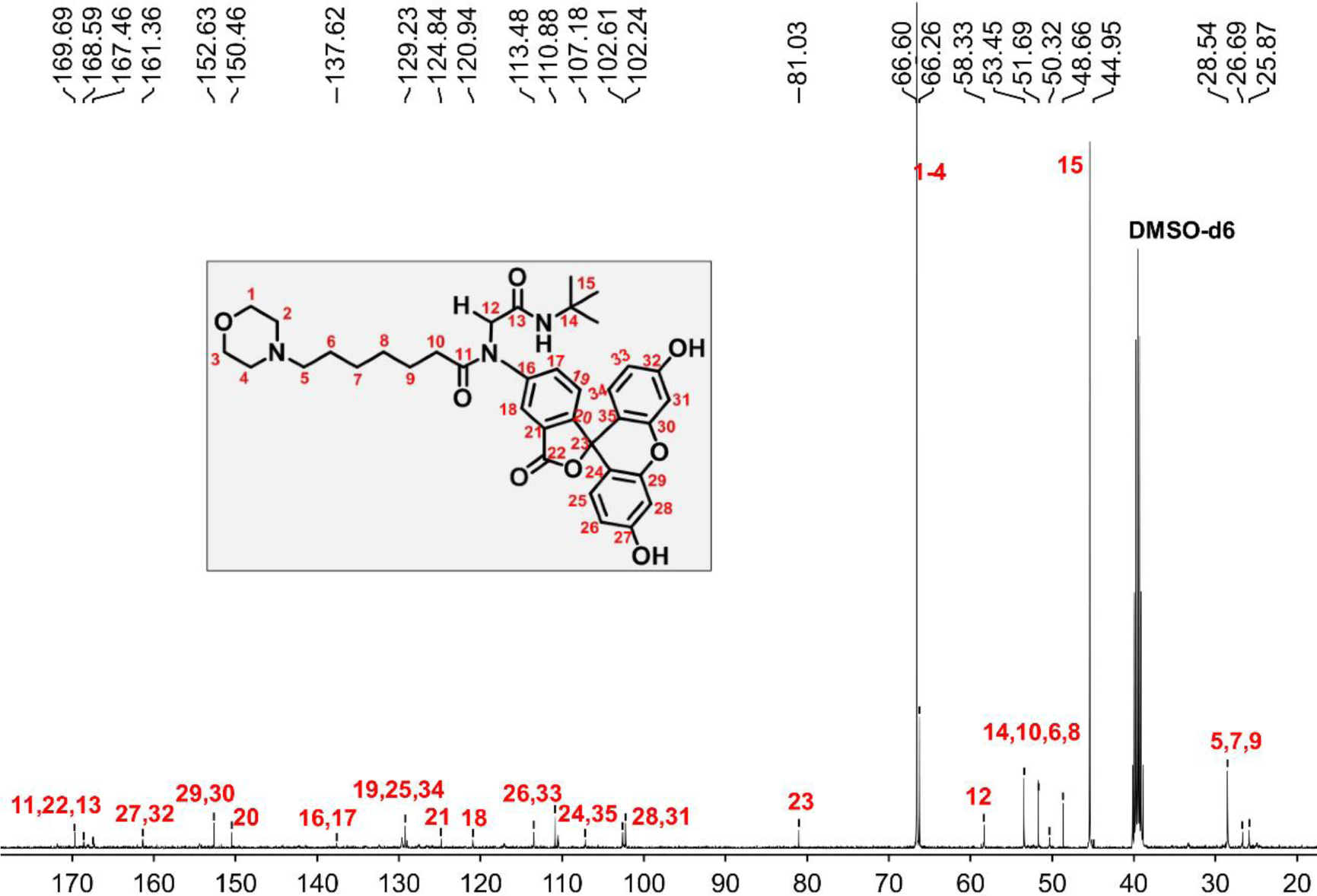
^13^C NMR spectrum of **27**.

**Figure S15.**
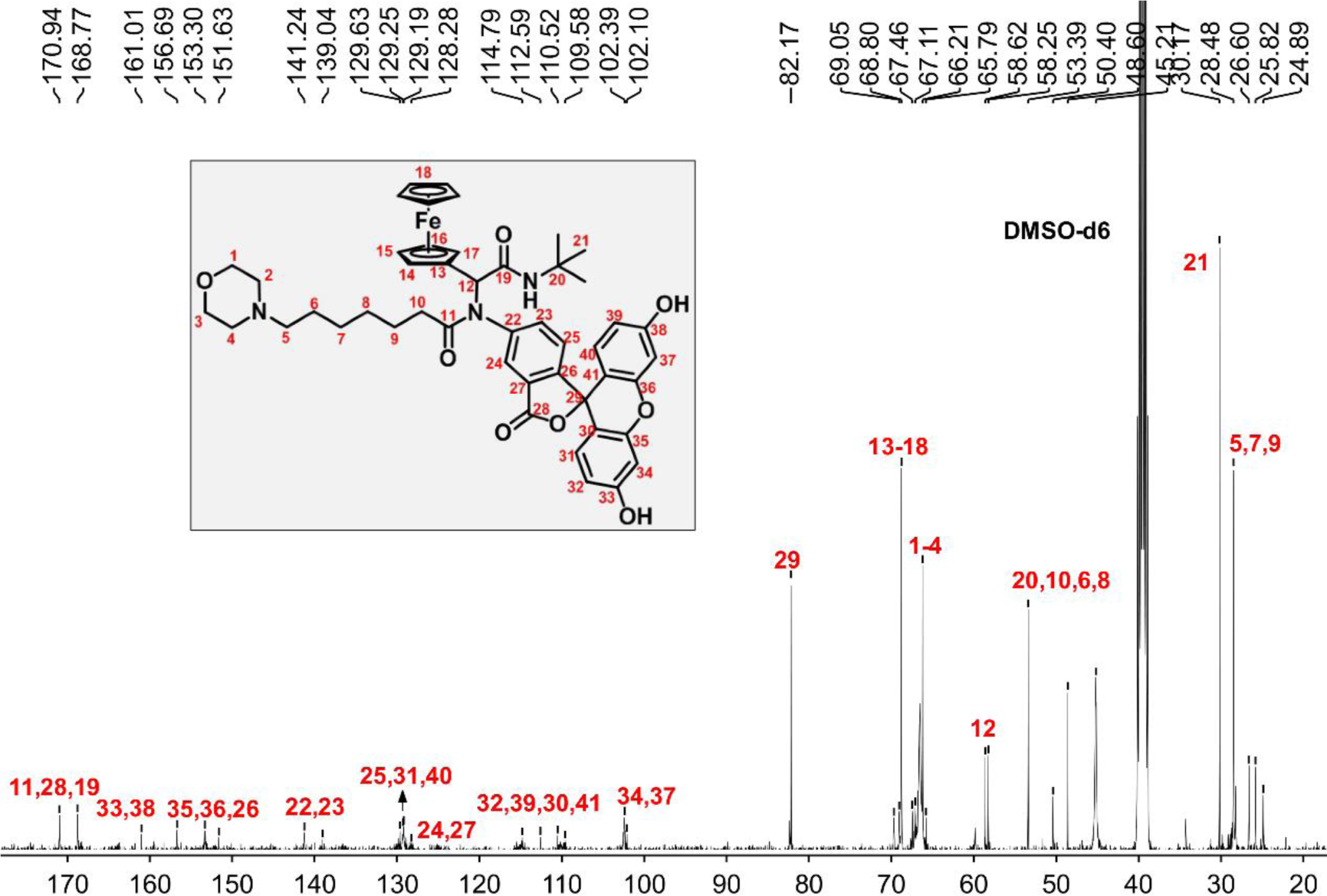
^13^C NMR spectrum of **28**.

**Figure S16.**
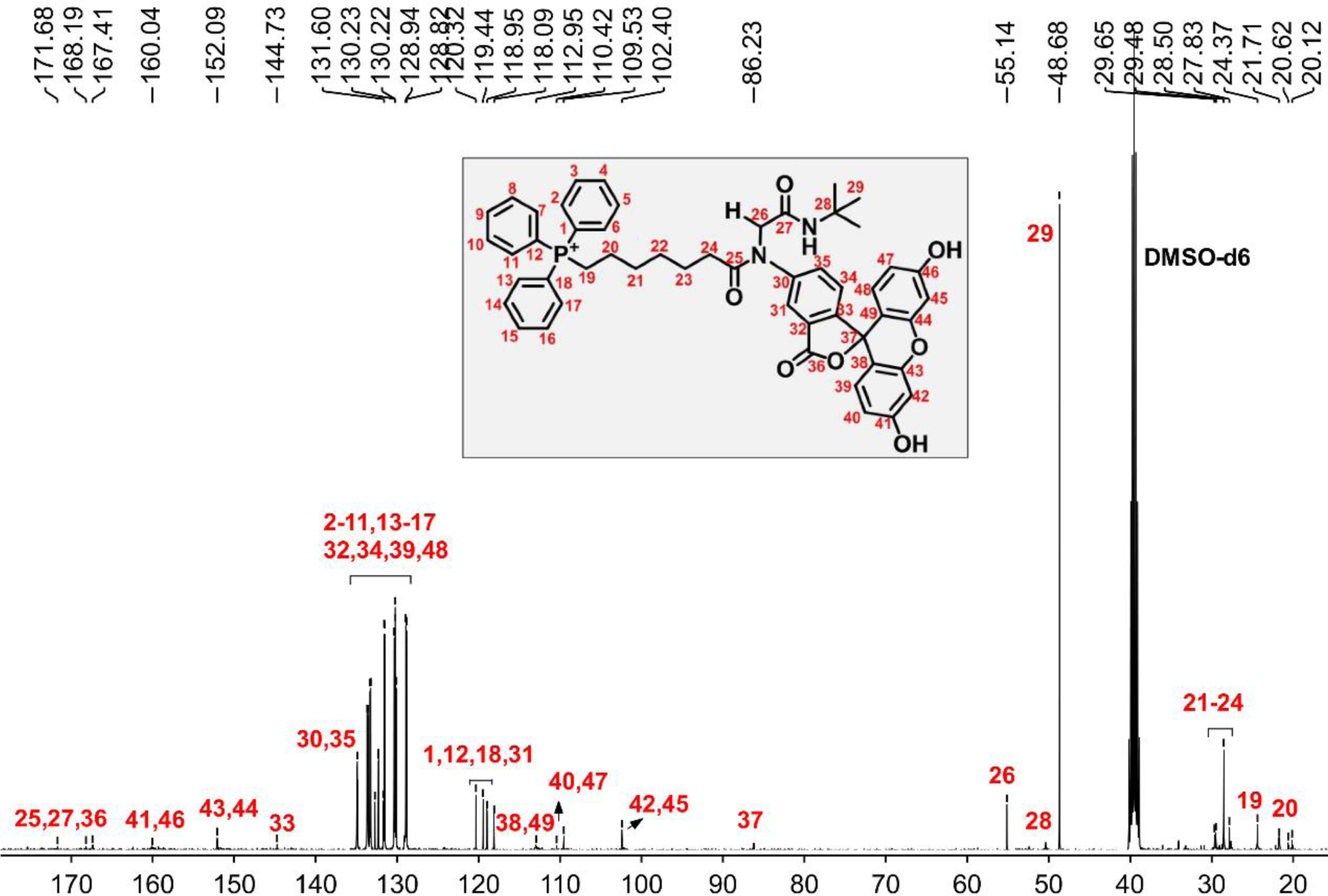
^13^C NMR spectrum of **29**.

**Figure S17.**
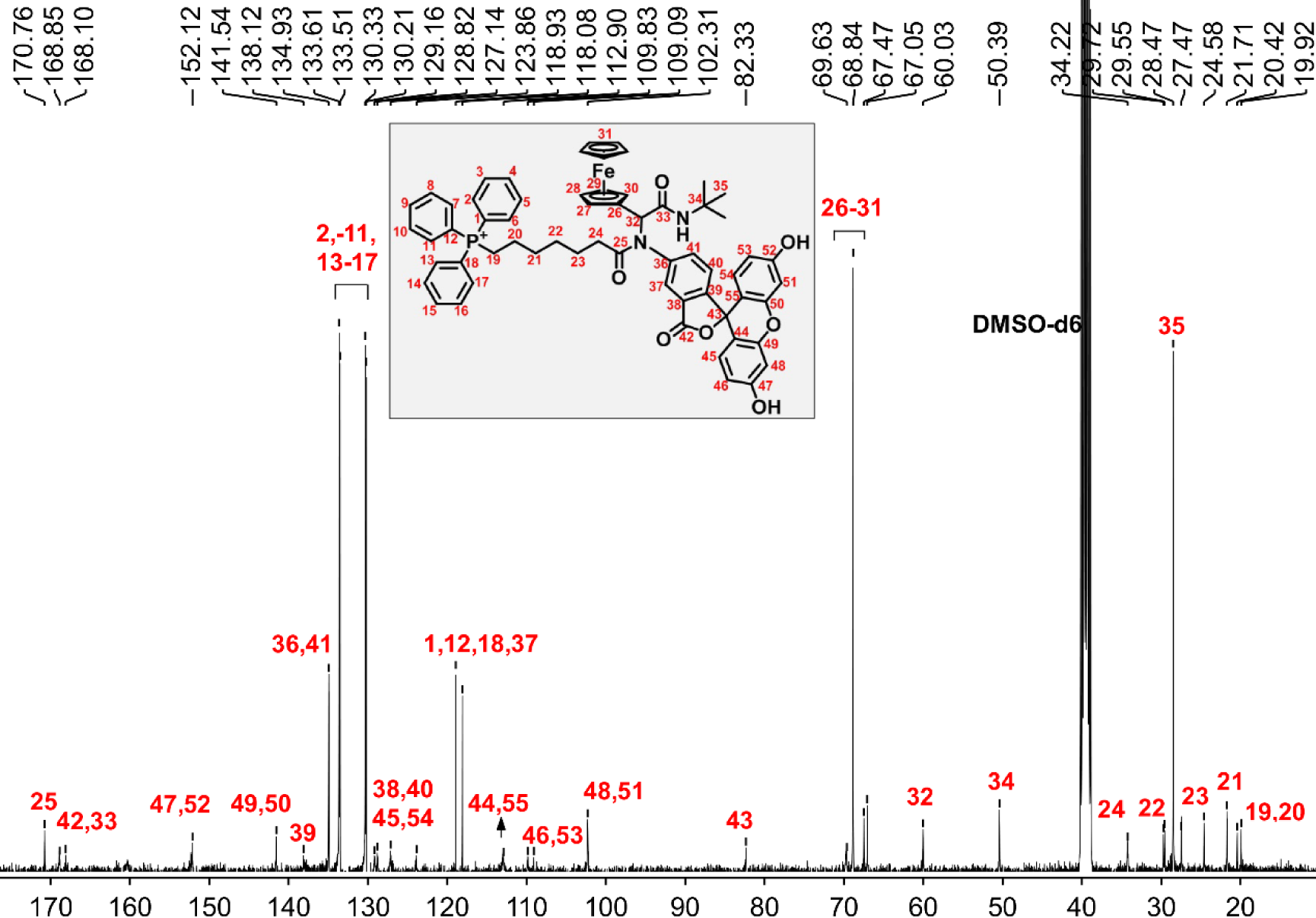
^13^C NMR spectrum of **30**.

**Figure S18.**
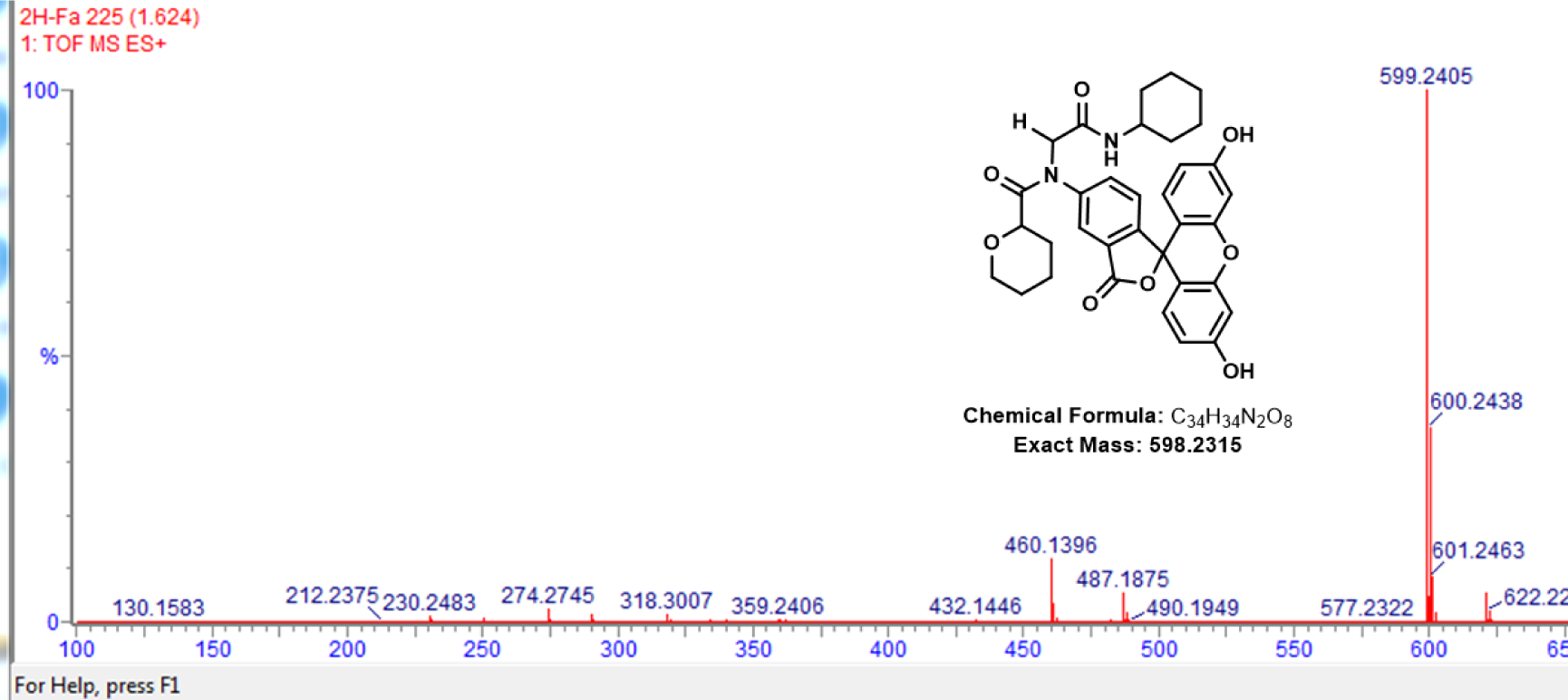
HRMS of **14**. m/z = 599.2405 Da [M(**14**+H)], m/z = 600.2438 Da [M(**14**+2H)].

**Figure S19.**
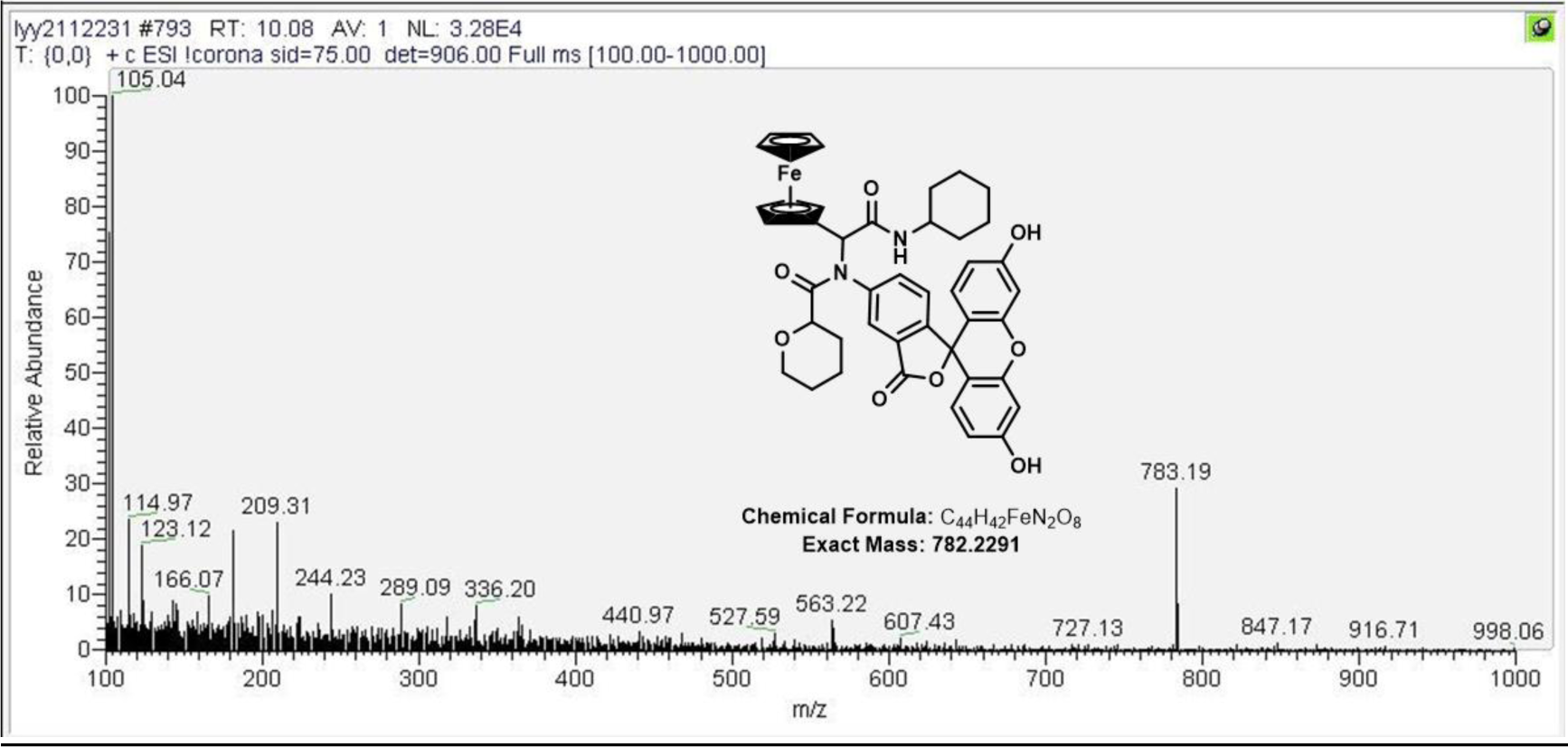
LCMS of **15**. m/z = 783.19 Da [M(**15**+H)].

**Figure S20.**
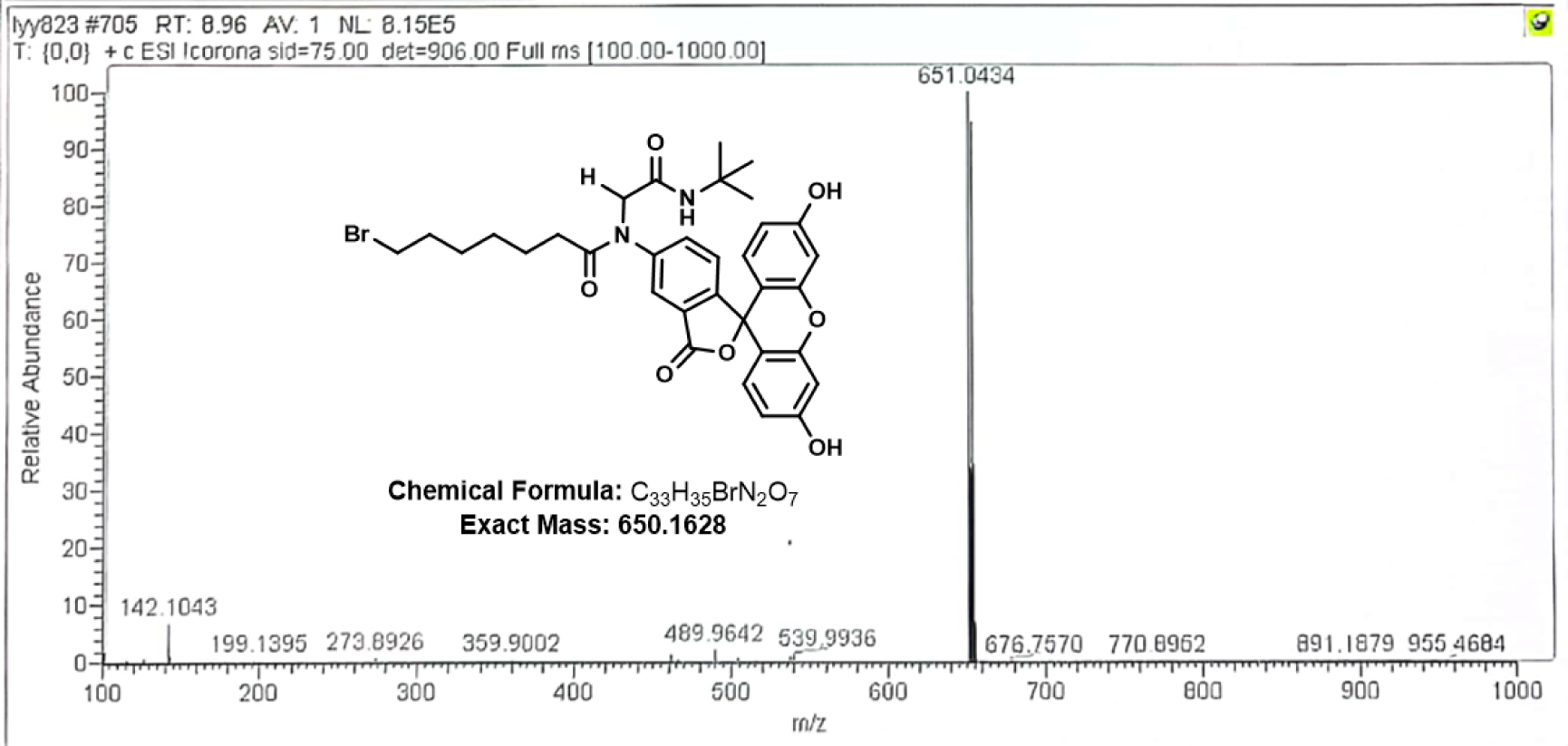
LCMS of **16**. m/z = 651.0434 Da [M(**16**+H)].

**Figure S21.**
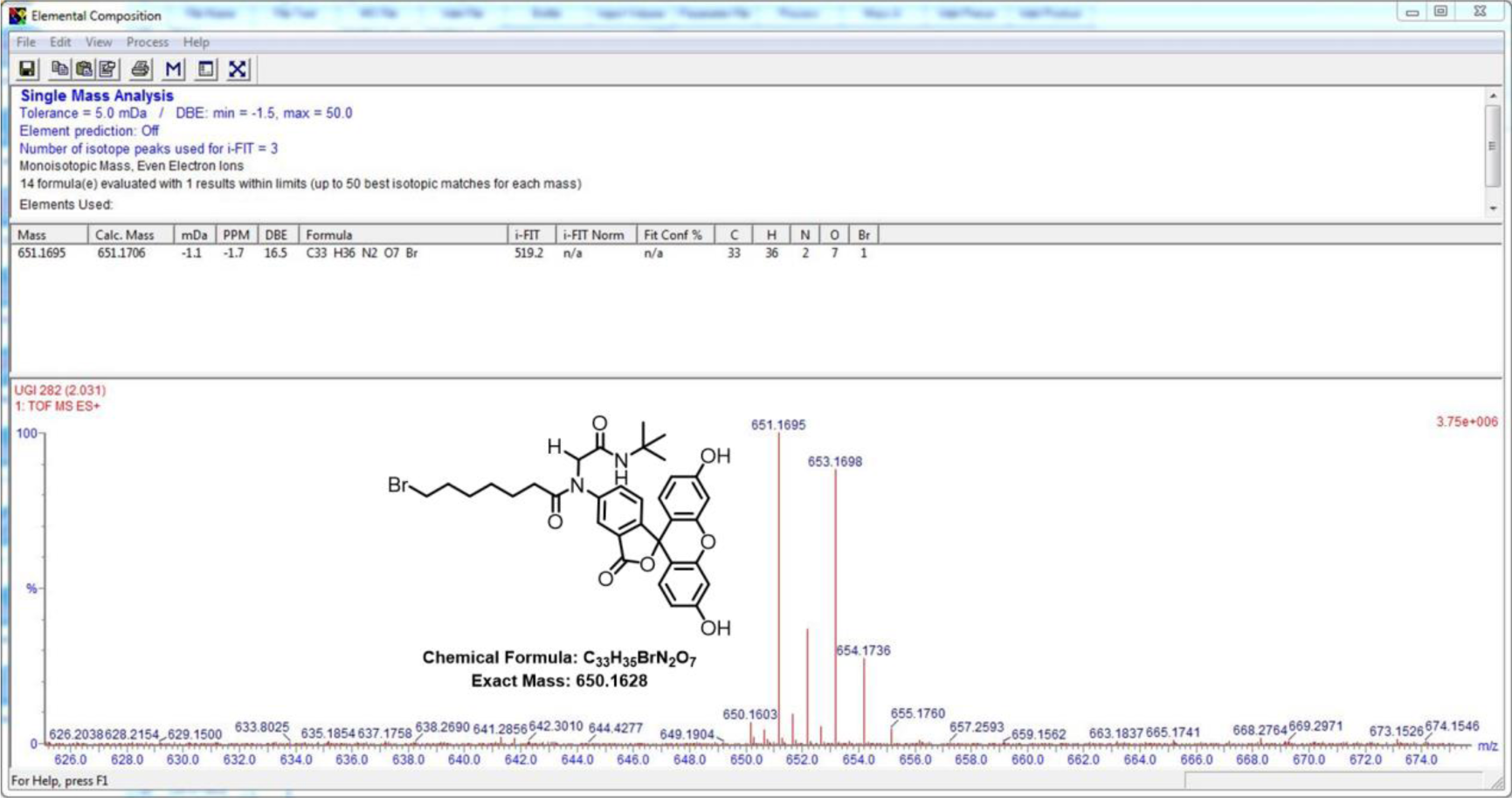
HRMS of **16**. m/z = 650.1603 Da [M(**16**)], m/z = 651.1695 Da [M(**16**+H)], m/z = 653.1698 Da [M(**16**+3H)].

**Figure S22.**
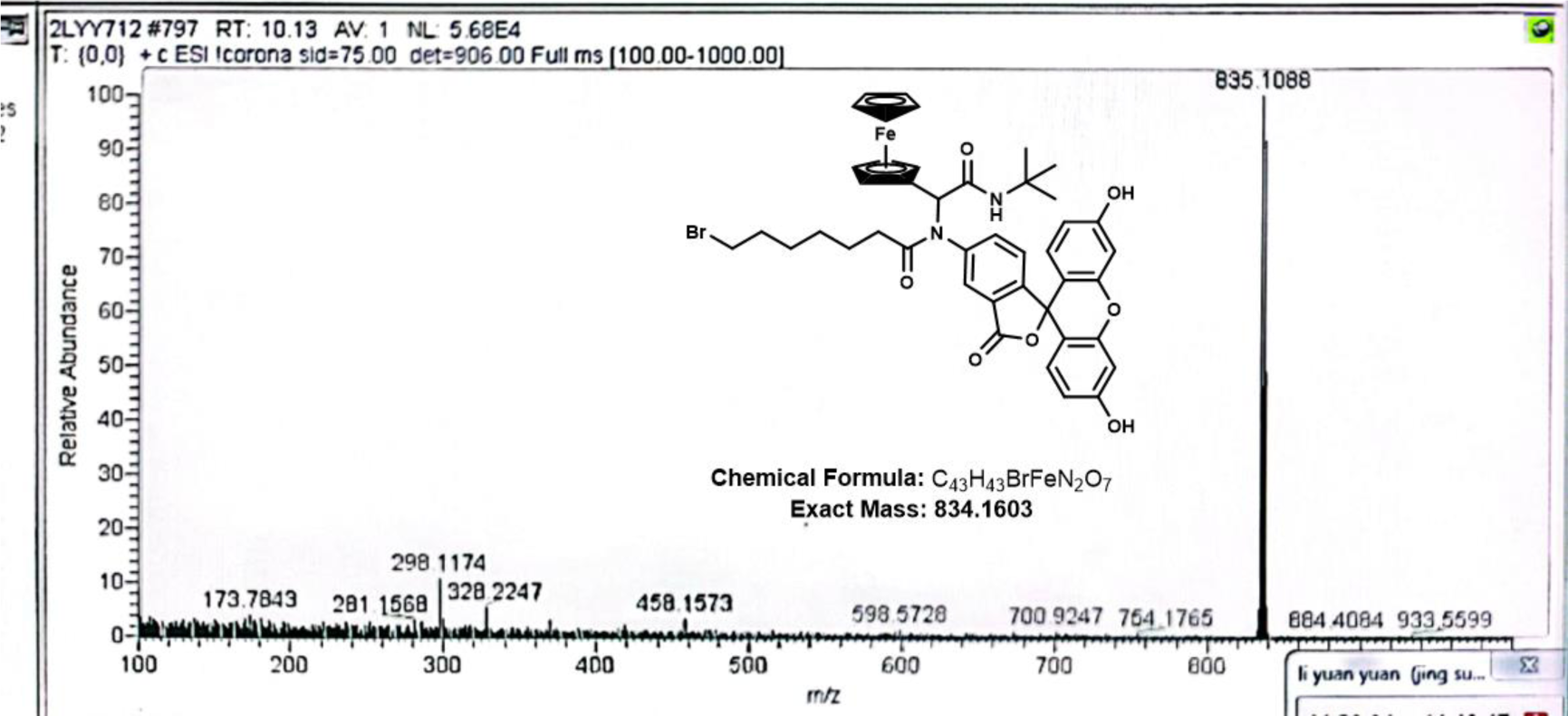
LCMS of **17**. m/z = 835.1088 Da [M(**17**+H)].

**Figure S23.**
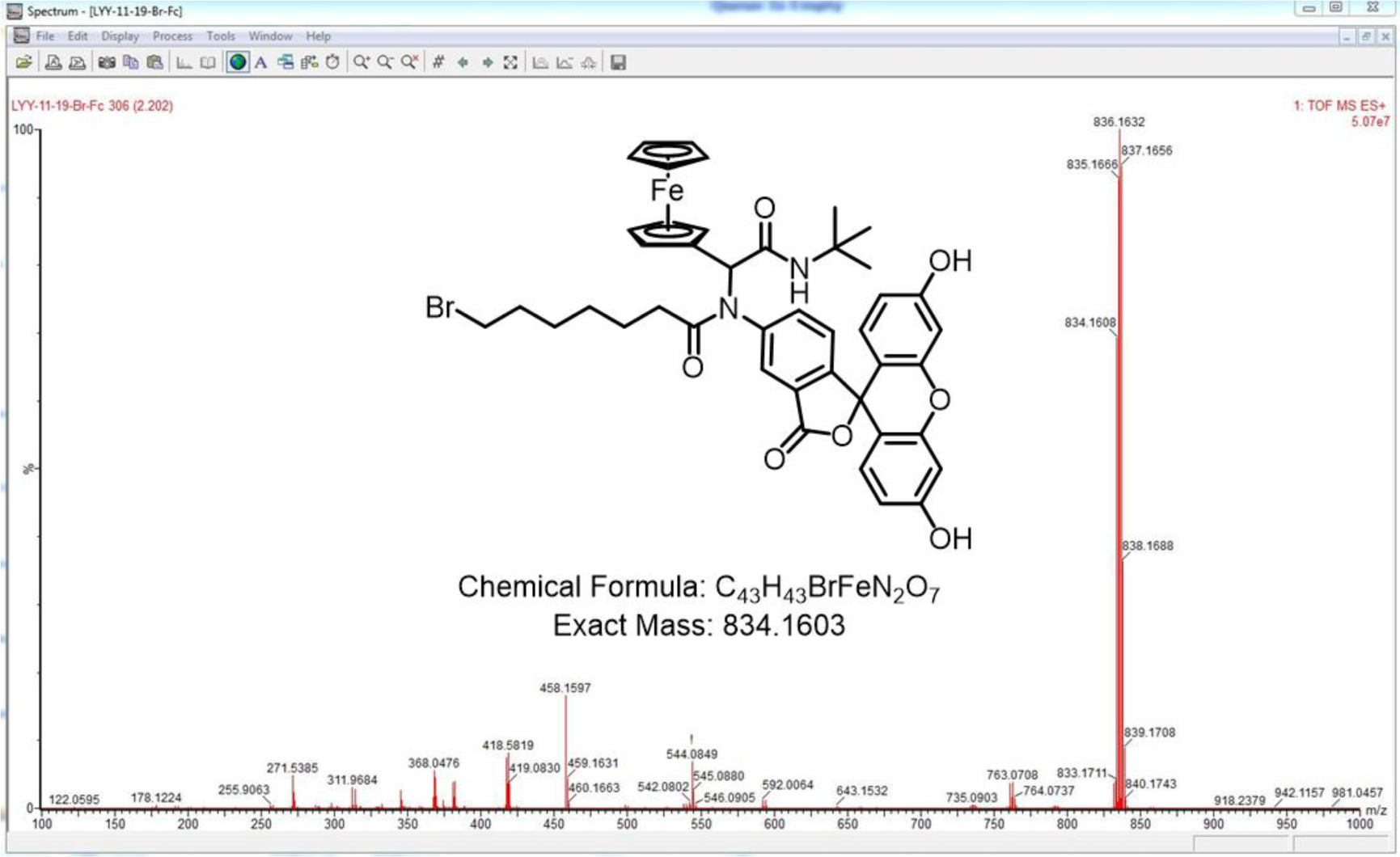
HRMS of **17**. m/z = 834.1608 Da [M(**17**)], m/z = 835.1666 Da [M(**17**+H)], m/z = 836.1632 Da [M(**17**+2H)], m/z = 837.1656 Da [M(**17**+3H)], m/z = 838.1688 Da [M(**17**+4H)].

**Figure S24.**
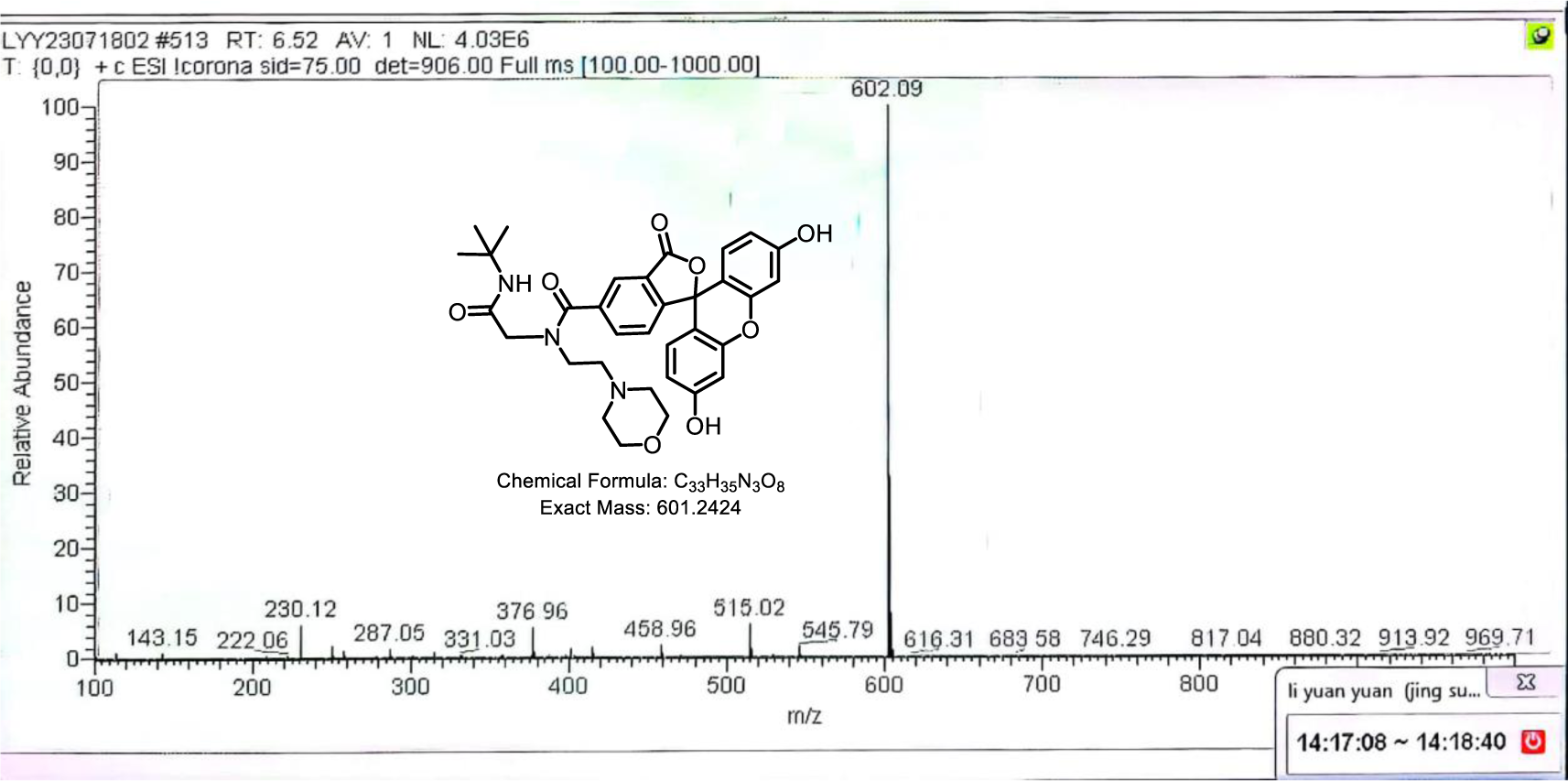
HPLC-MS of **18**. m/z = 601.24 Da [M(**18**)], m/z = 602.09 Da [M(**18**+H)].

**Figure S25.**
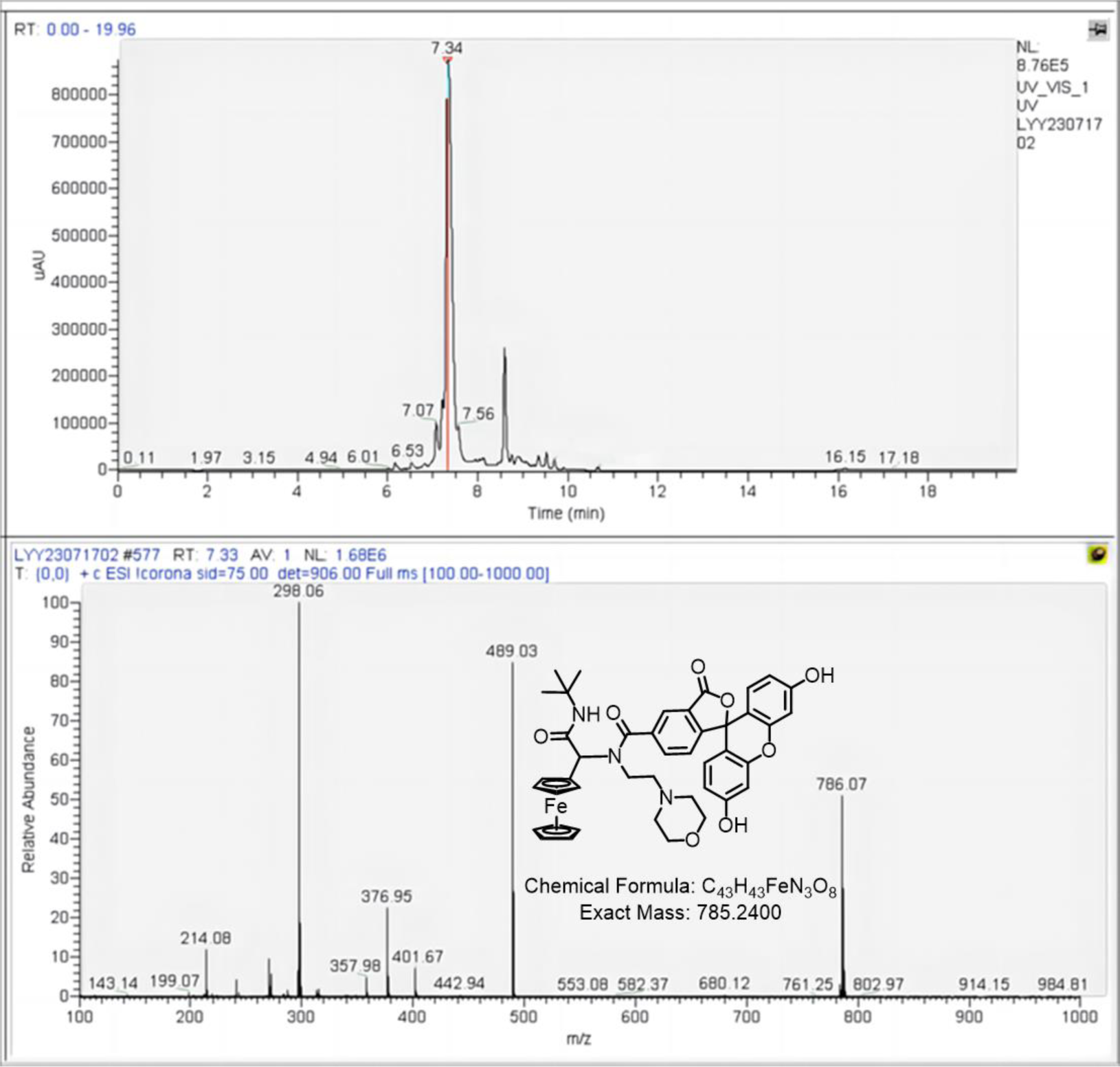
HPLC-MS of **19**. m/z = 785.2400 Da [M(**19**)], m/z = 786.07 Da [M(**19**+H)].

**Figure S26.**
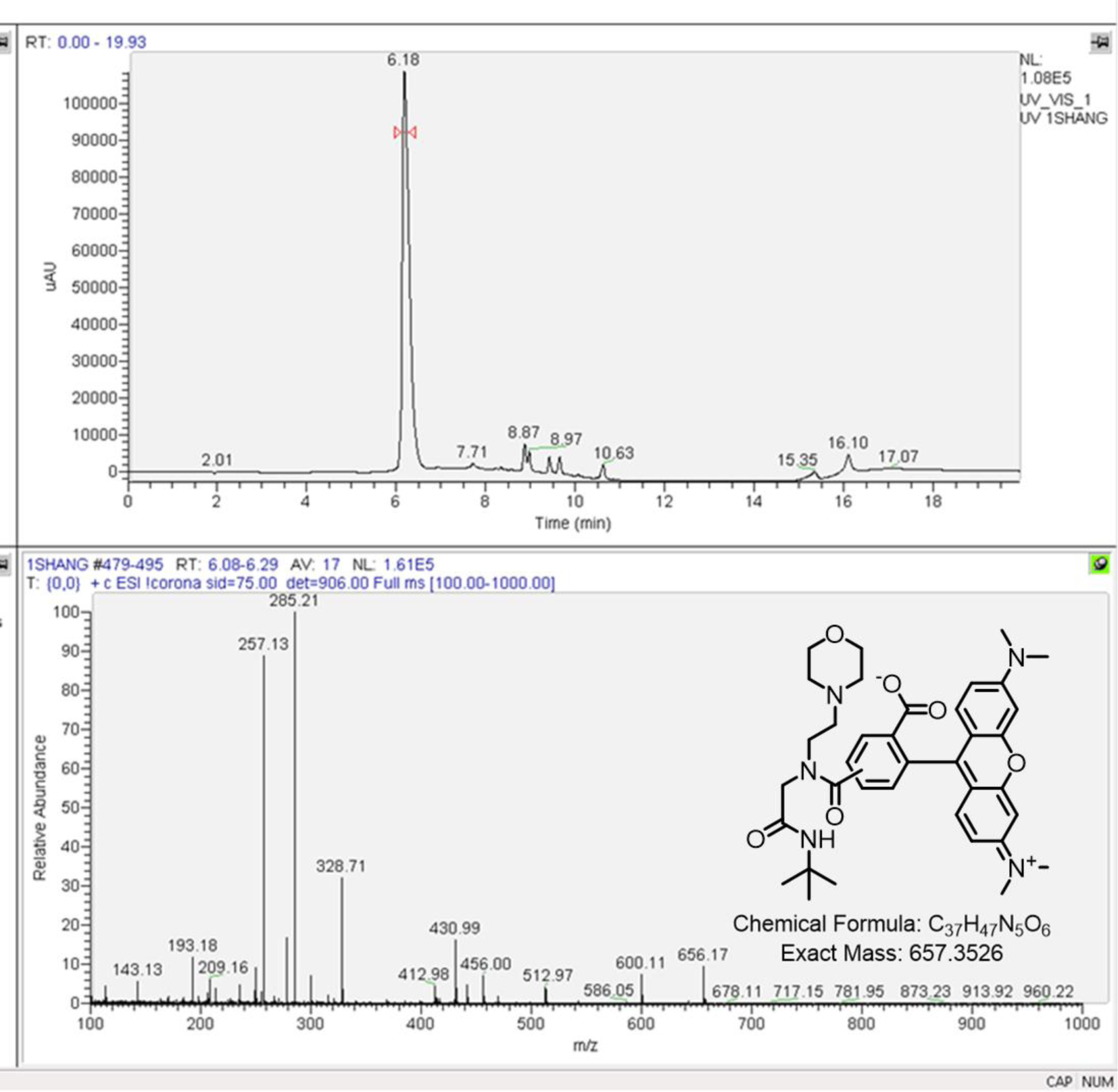
HPLC-MS of **20**. m/z = 656.17 Da [M(**20**)].

**Figure S27.**
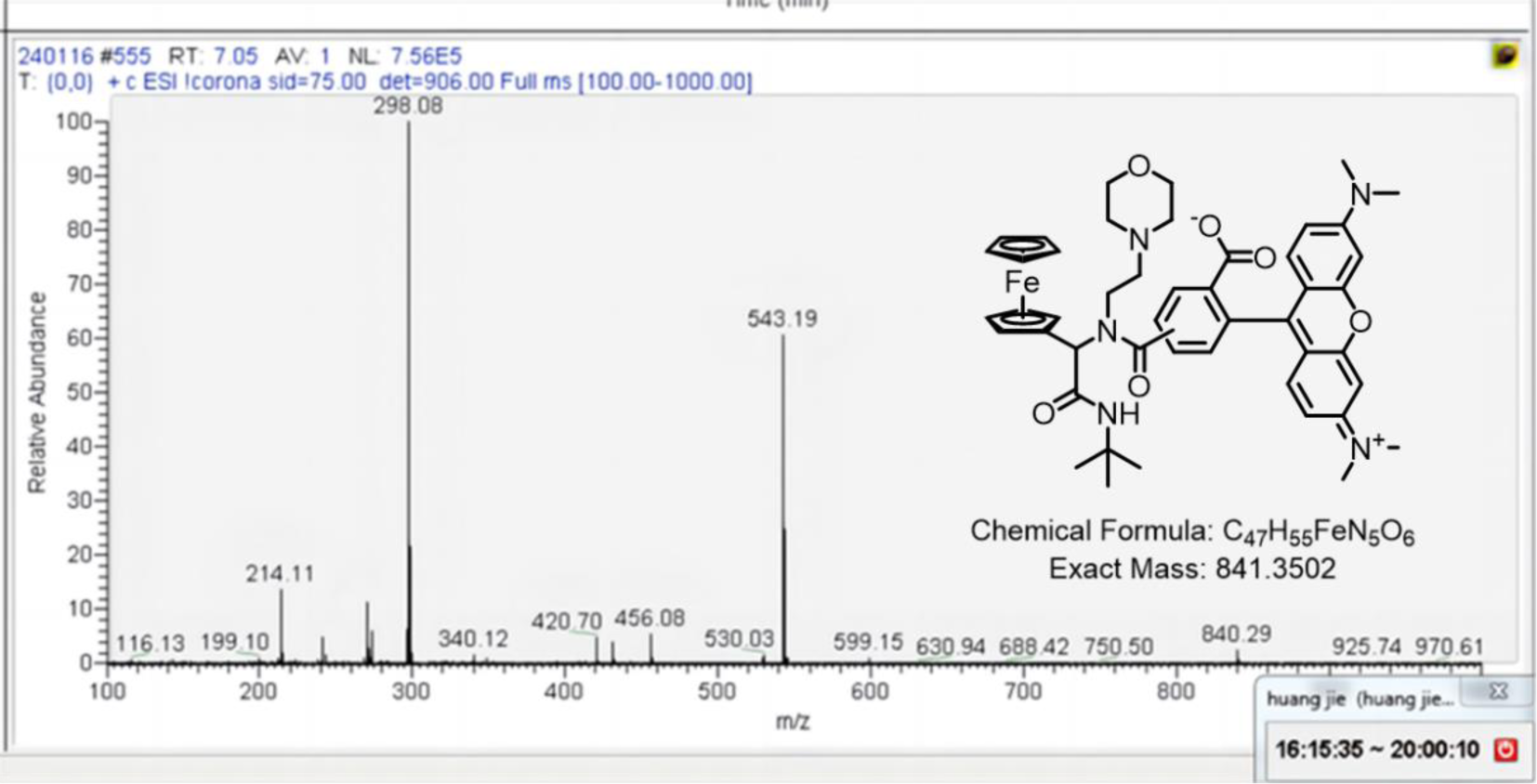
HPLC-MS of **21**. m/z = 840.29 [M(**21**)].

**Figure S28.**
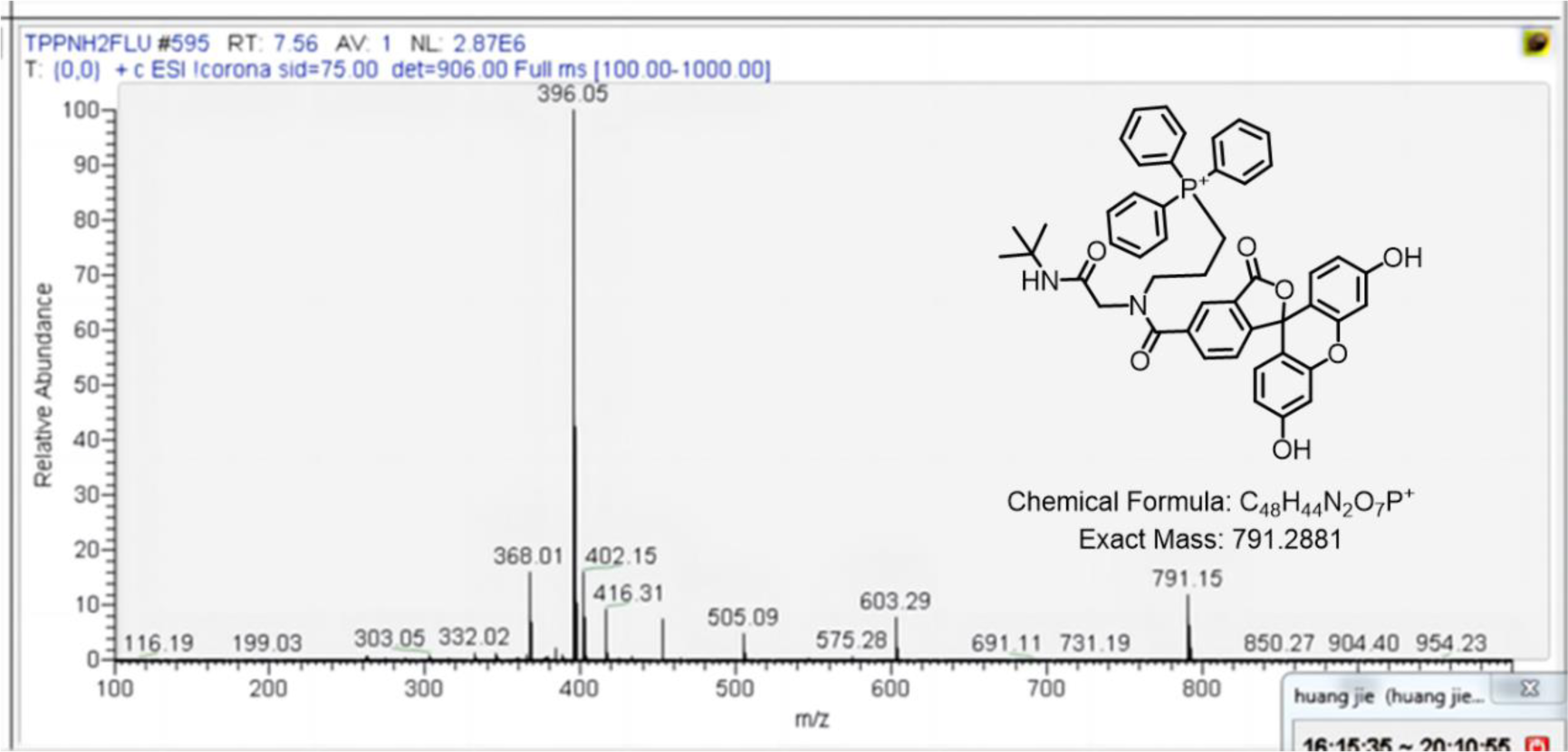
HPLC-MS of **22**. m/z = 791.15 Da [M(**22**)].

**Figure S29.**
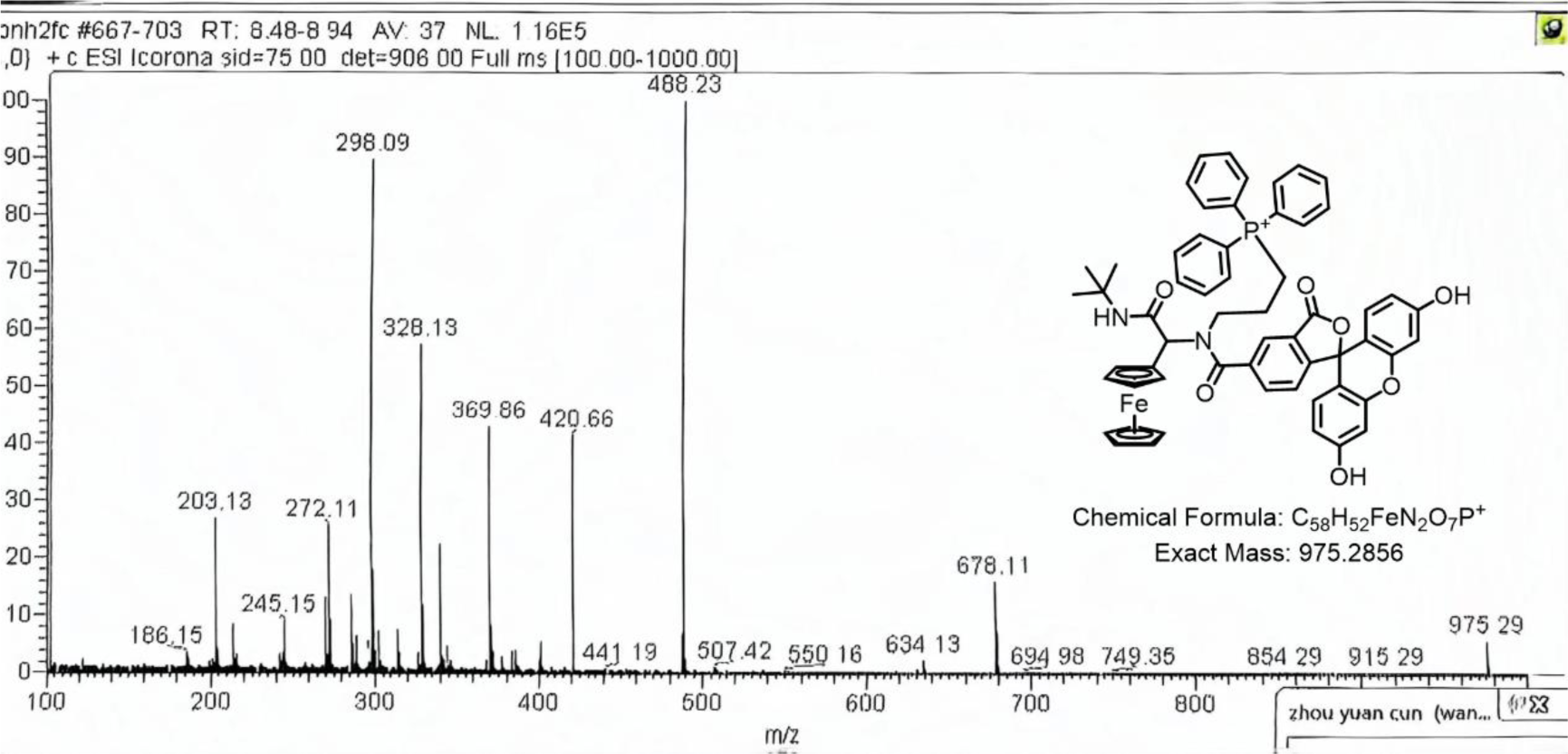
HPLC-MS of **23**. m/z = 975.29 Da [M(**23**)].

**Figure S30.**
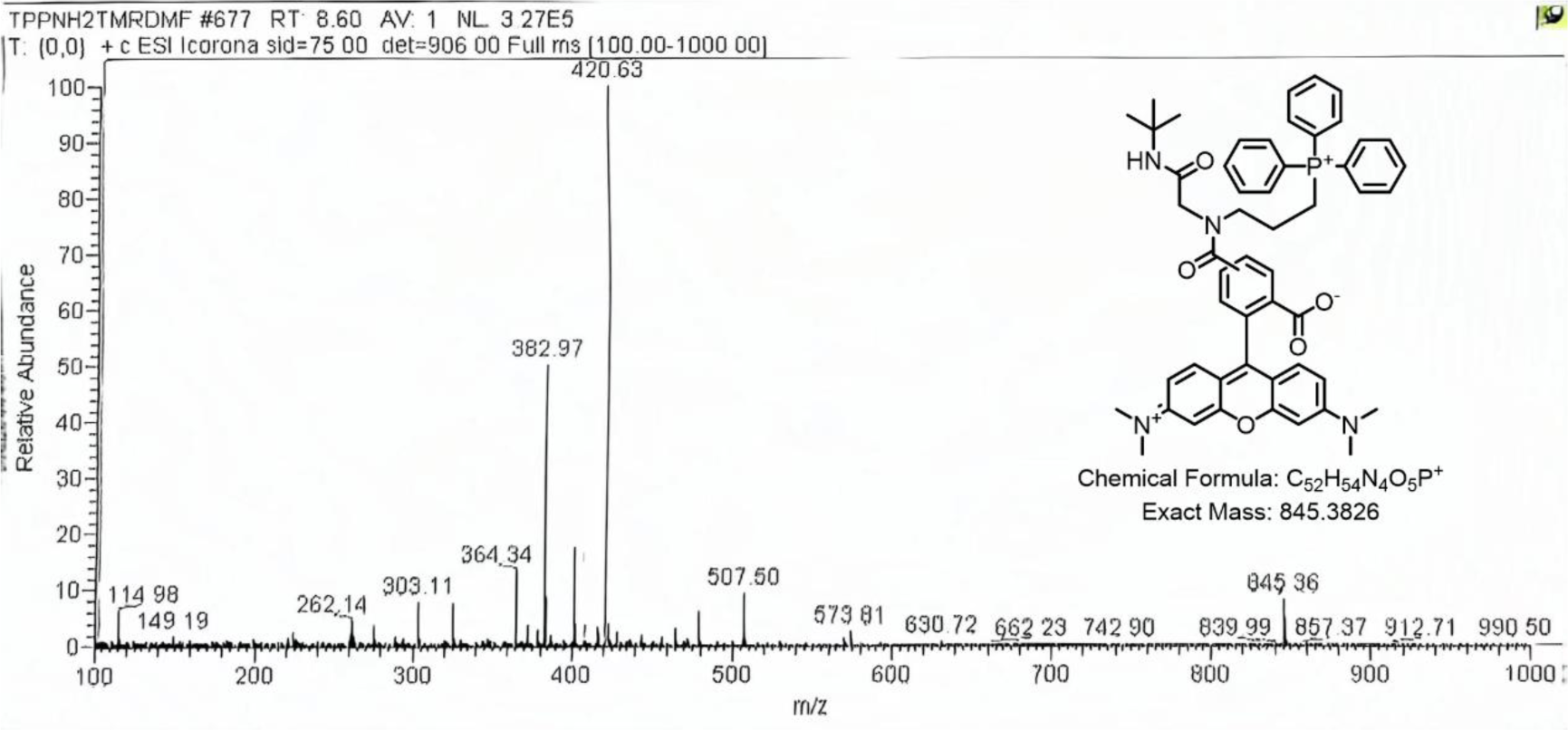
HPLC-MS of **24**. m/z = 845.36 Da [M(**24**)].

**Figure S31.**
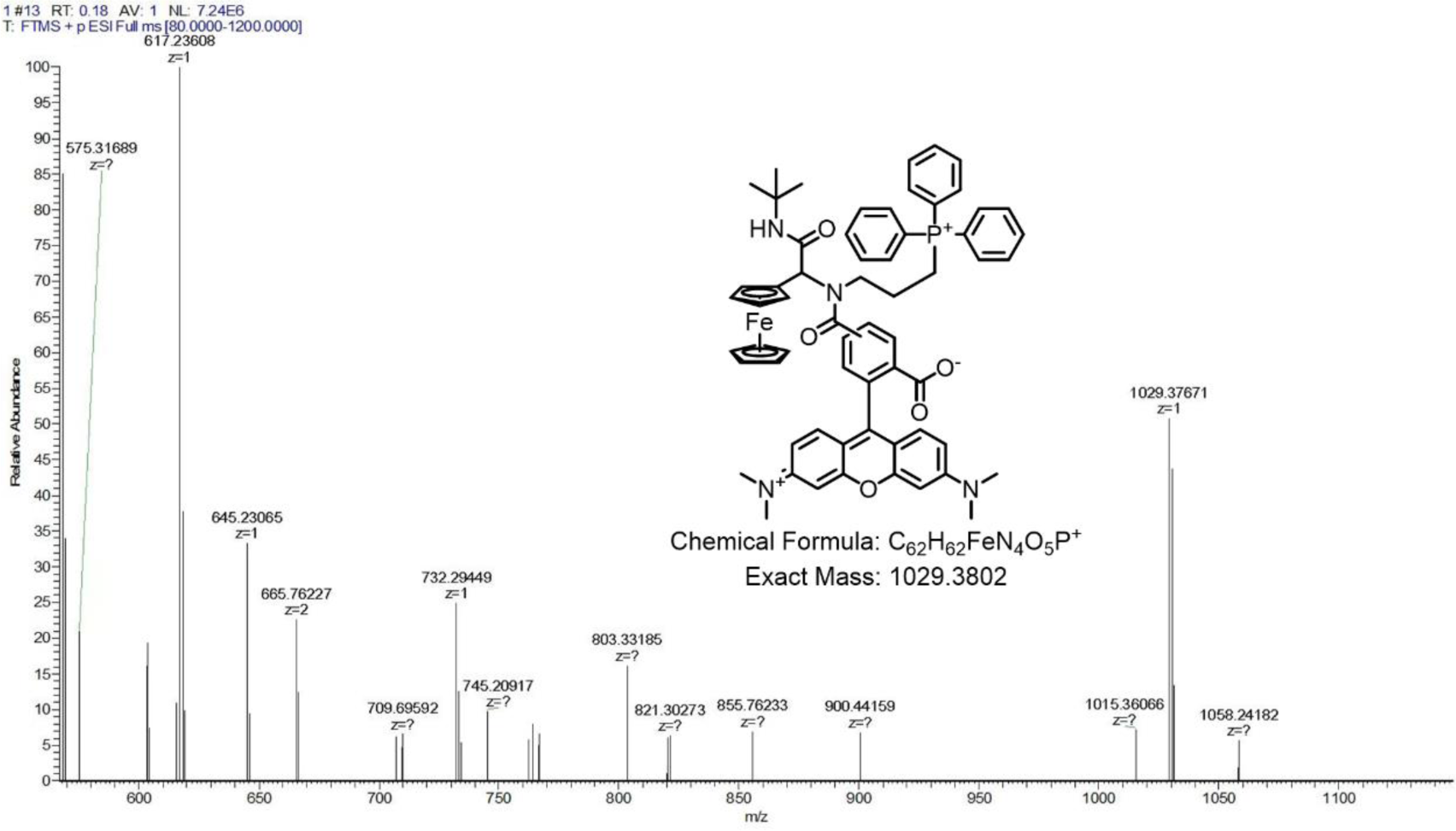
HPLC-MS of **25**. m/z = 1029.3767 Da [M(**25**)].

**Figure S32.**
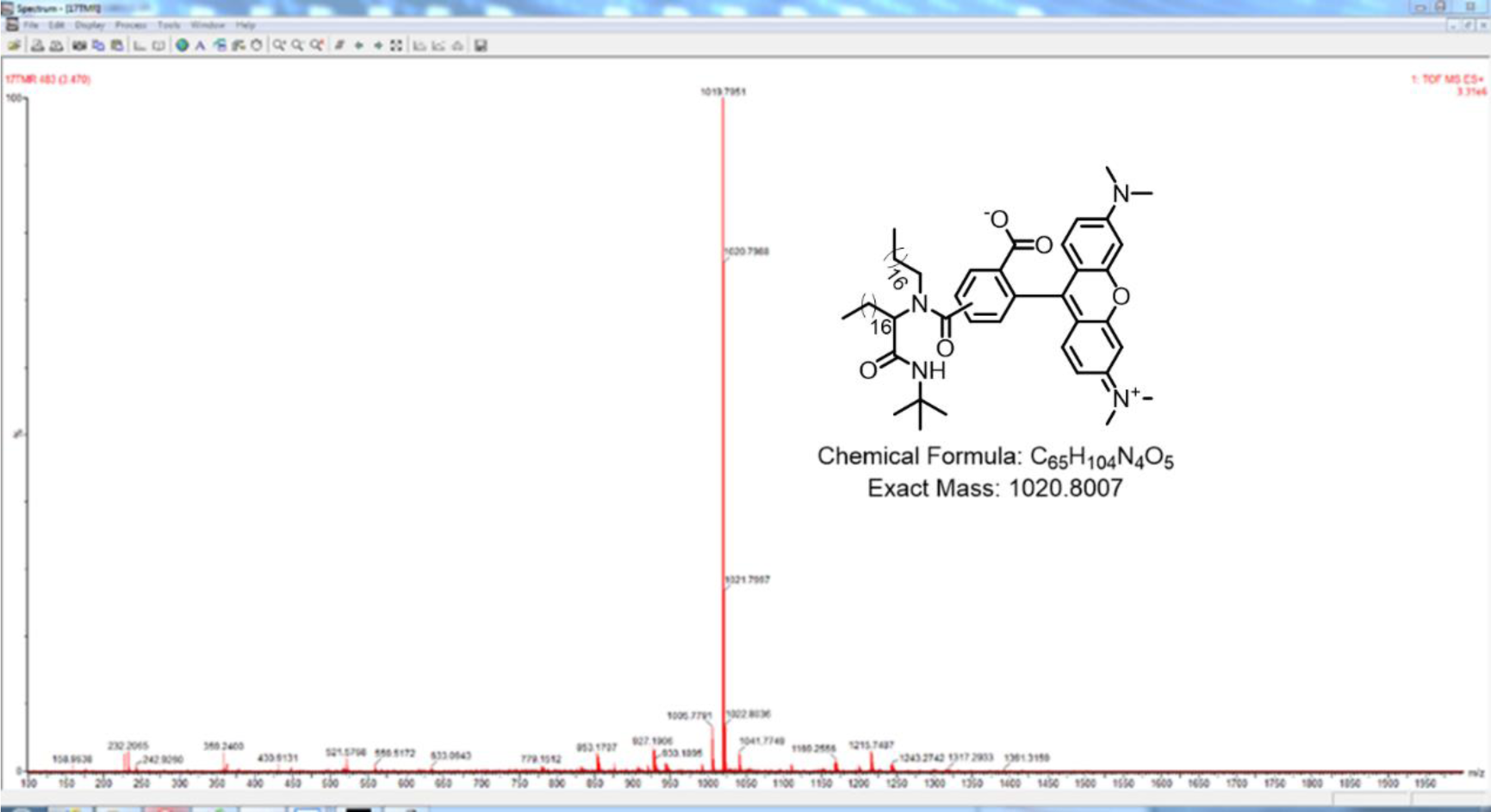
HR-MS of **26**. m/z = 1020.7988 Da [M(**26**)].

**Figure S33.**
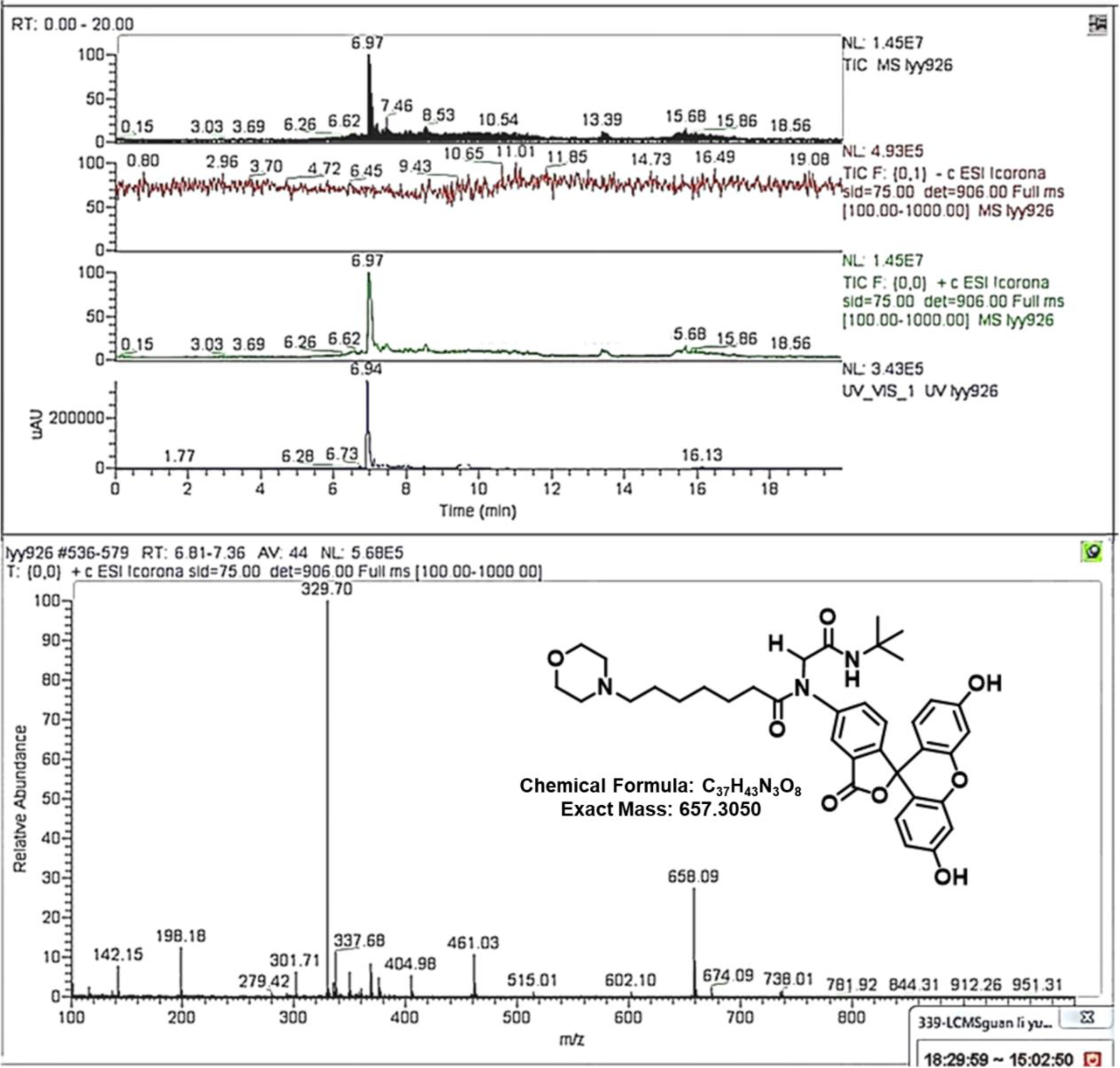
HPLC-MS of **27**. m/z = 658.09 Da [M(**27**+H)].

**Figure S34.**
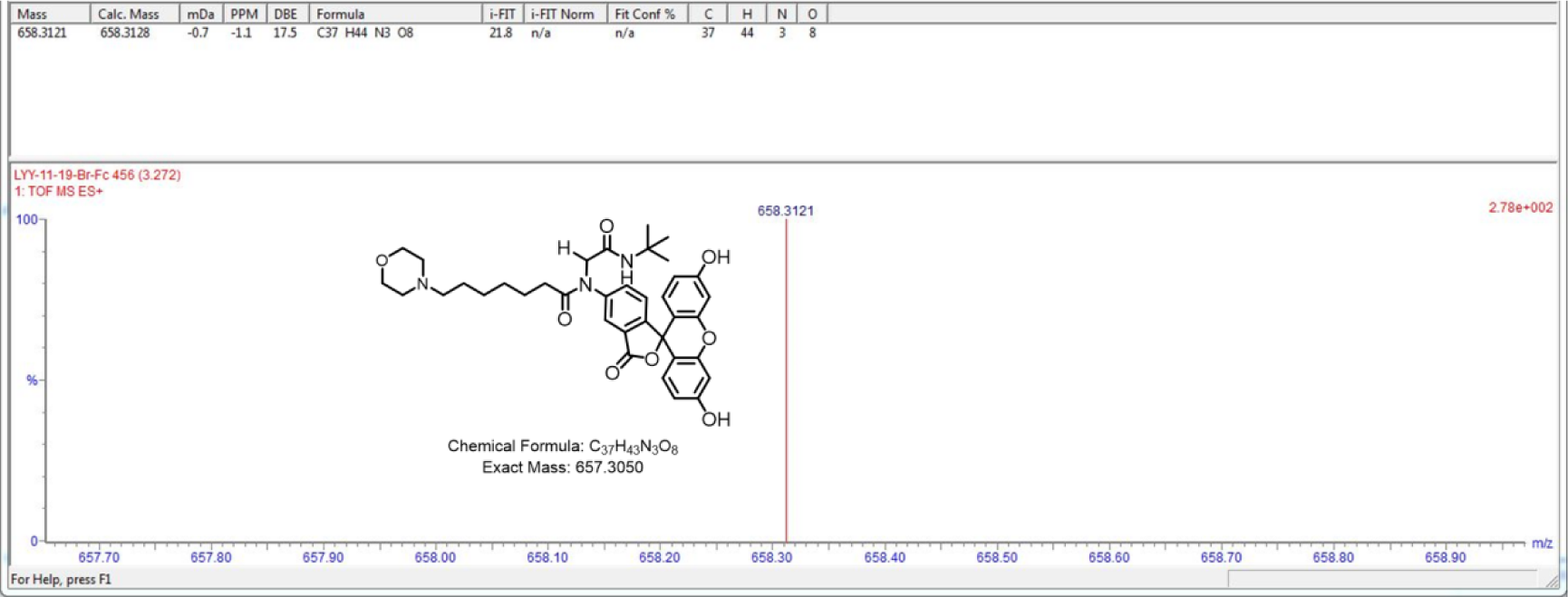
HRMS of **27**. m/z = 658.3121 Da [M(**27**+H)].

**Figure S35.**
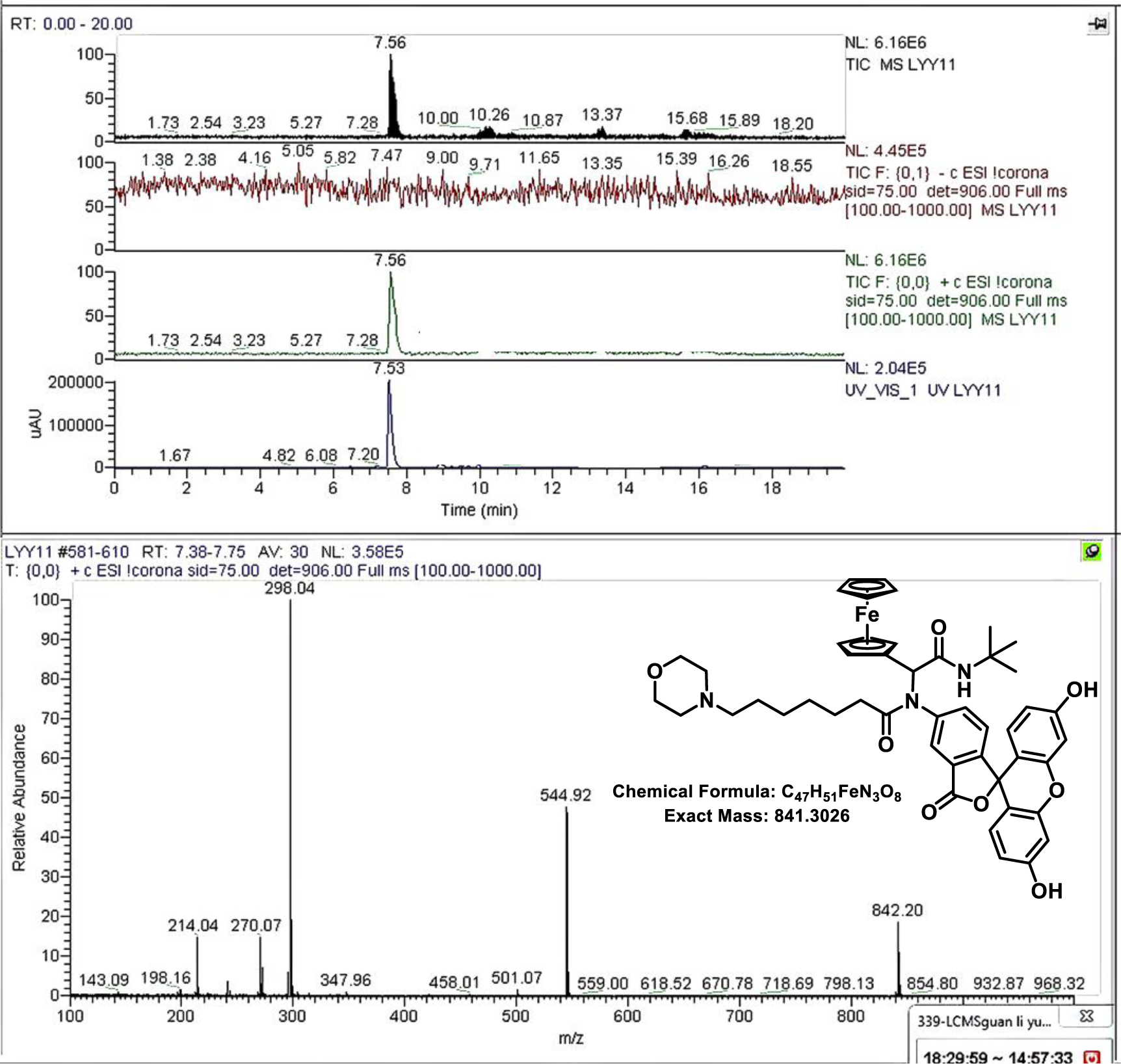
HPLC-MS of **28**. m/z = 842.20 Da [M(**28**+H)].

**Figure S36.**
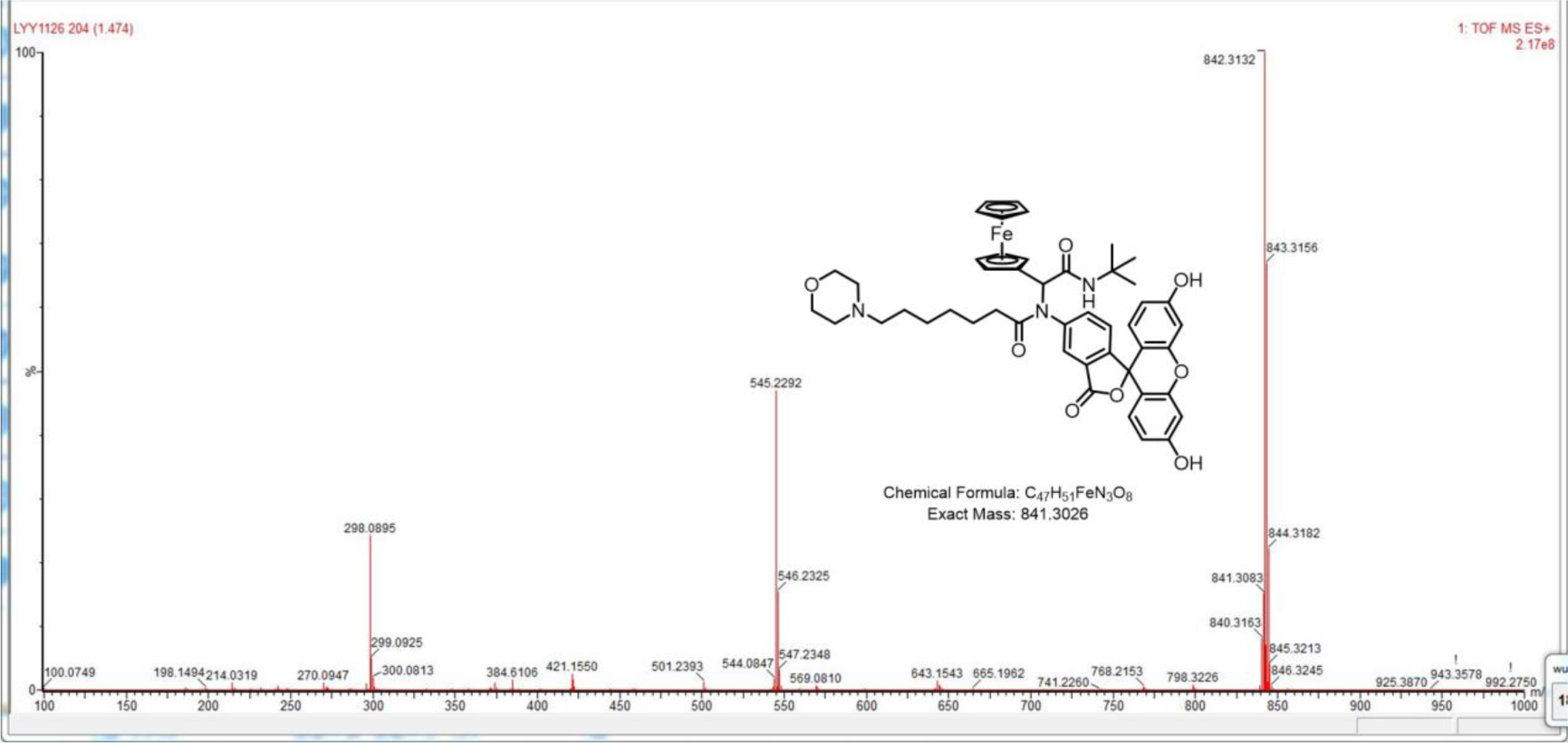
HRMS of **28**. m/z = 842.3132 Da [M(**28**+H)], m/z = 843.3156 Da [M(**28**+2H)], m/z = 844.3182 Da [M(**28**+3H)].

**Figure S37.**
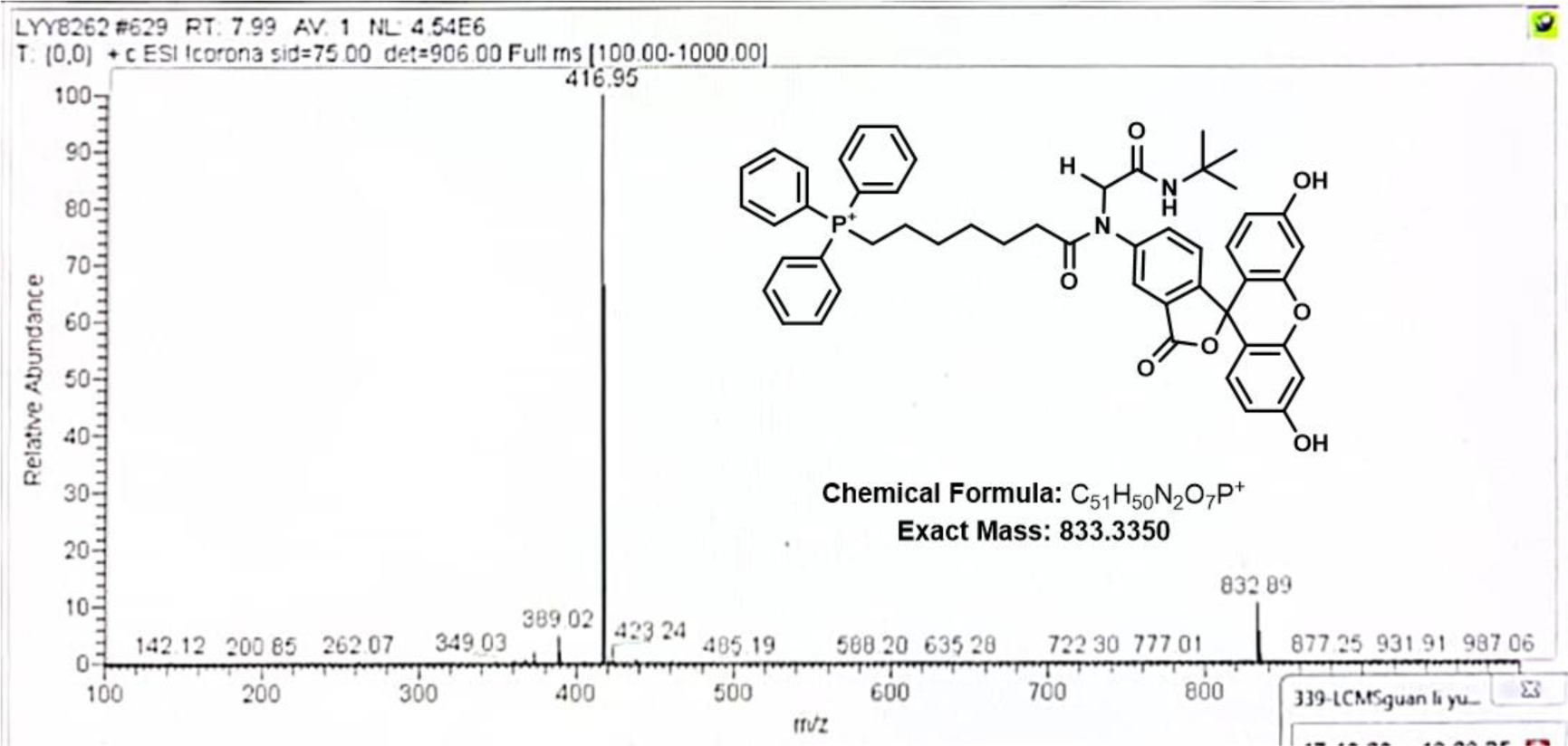
HPLC-MS of **29**. m/z = 832.89 Da [M(**29**)].

**Figure S38.**
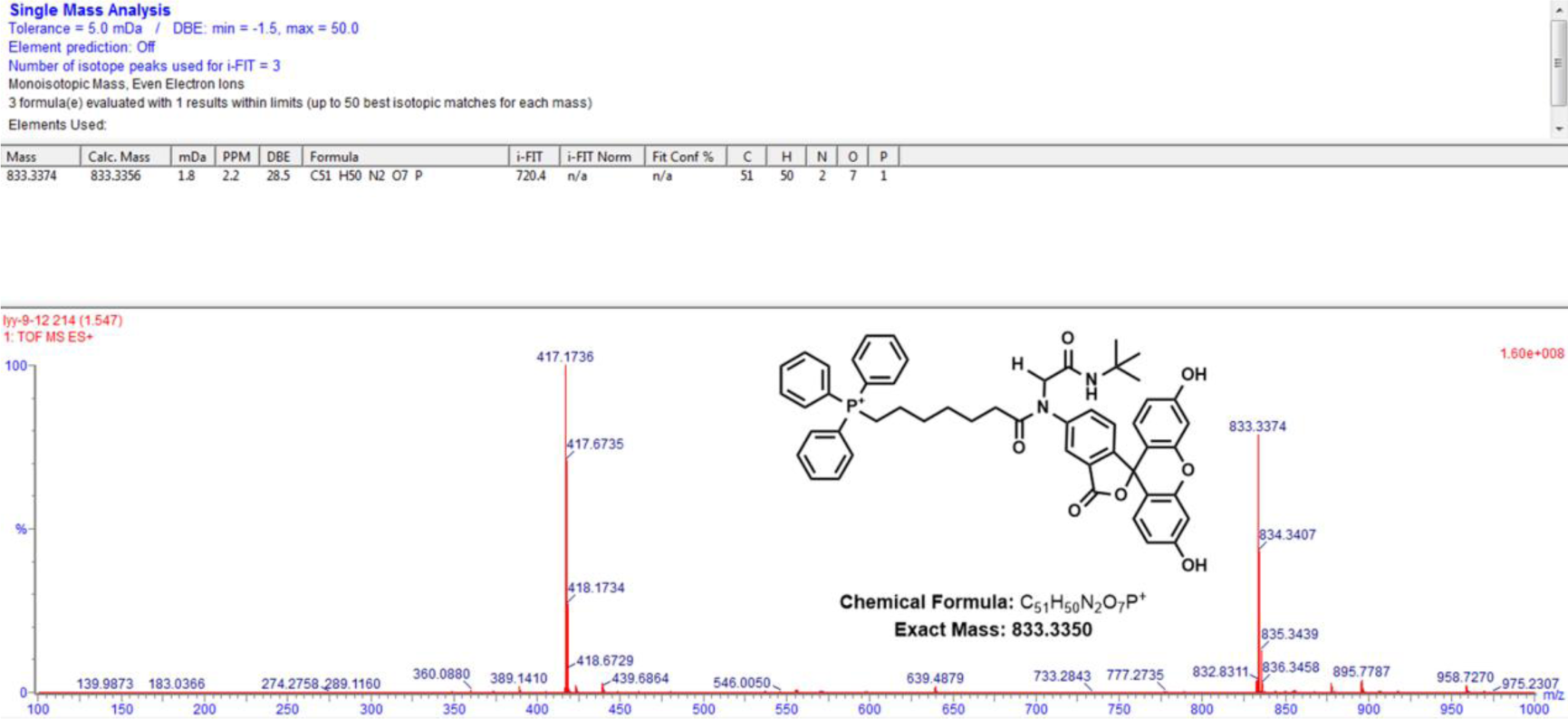
HRMS of **29**. m/z = 833.3374 Da [M(**29**)], m/z = 834.3407 Da [M(**29**+H)], m/z = 835.3439 Da [M(**29**+2H)].

**Figure S39.**
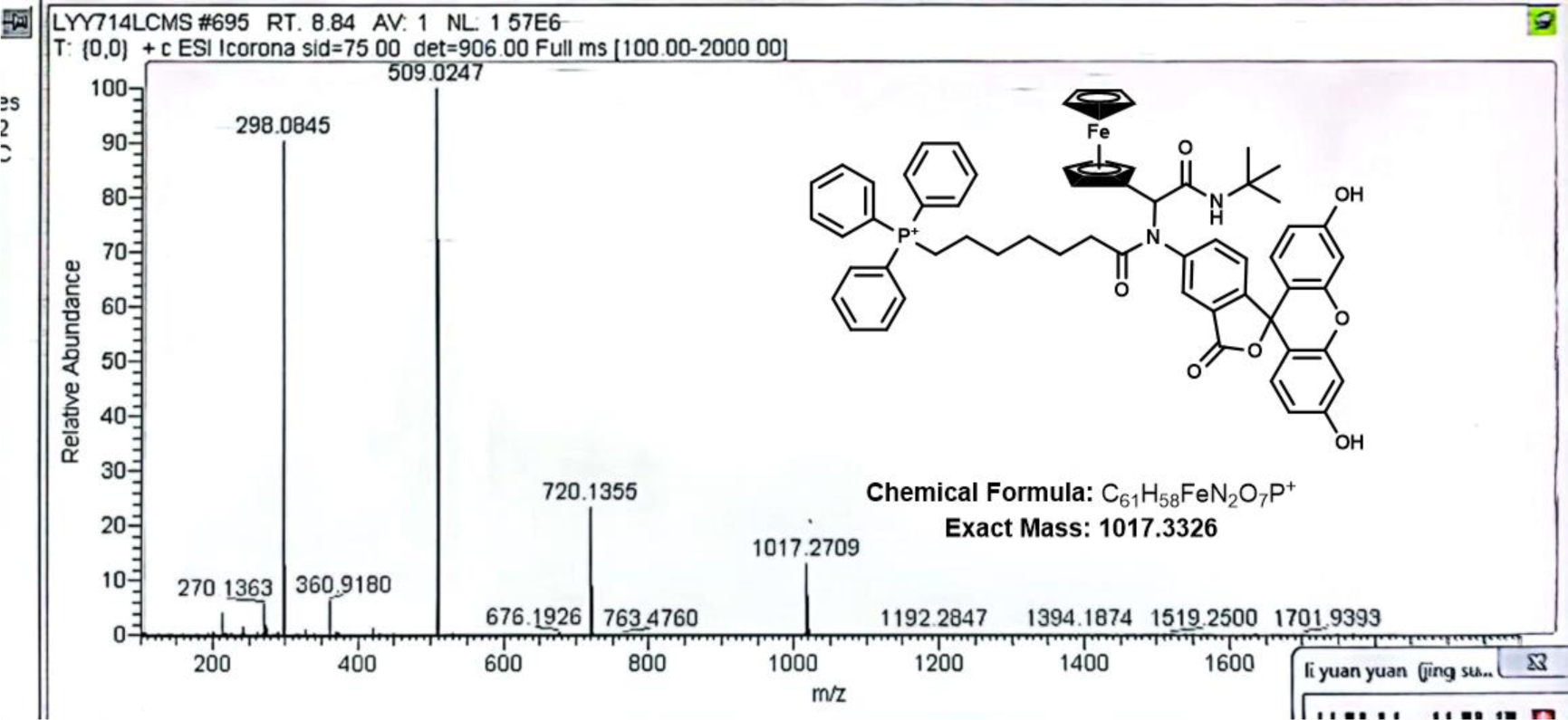
HPLC-MS of **30**. m/z = 1017.2709 Da [M(**30**)].

**Figure S40.**
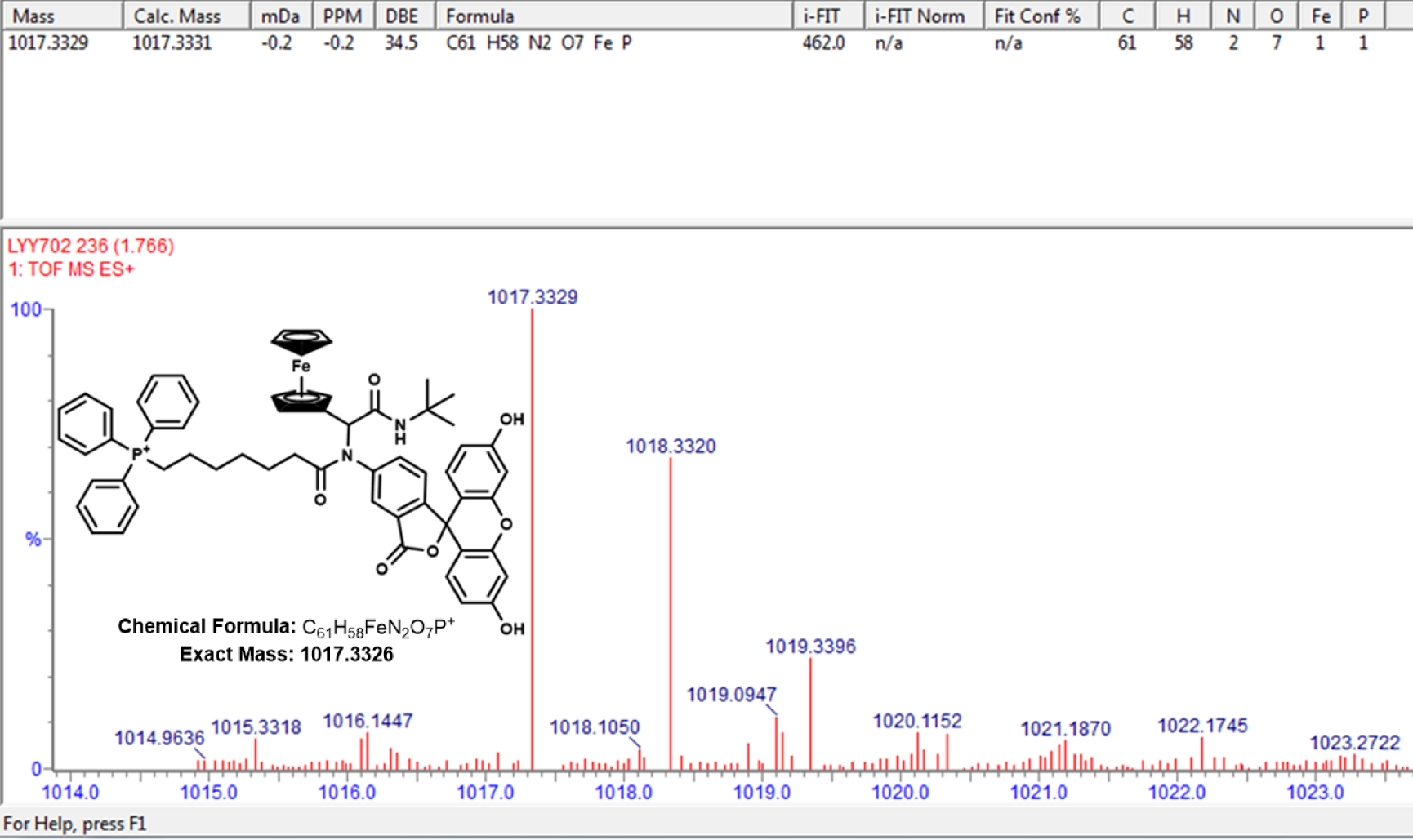
HRMS of **30**. m/z = 1017.3329 Da [M(**30**)], m/z = 1018.3320 Da [M(**30**+H)], m/z = 1019.3396 Da [M(**30**+2H)].

#### LC/MS Characterizations of Compound 18-25 Purified by Preparative HPLC

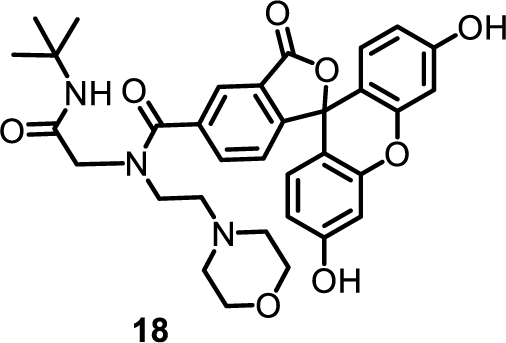

N-(2-(tert-butylamino)-2-oxoethyl)-3’,6’-dihydroxy-N-(2-morpholinoethyl)-3-oxo-3H-spiro[isobenzofuran-1,9’-xanthene]-5-carboxamide

HPLC-MS of **18**. m/z = 601.24 Da [M(**18**)], m/z = 602.09 Da [M(**18**+H)]

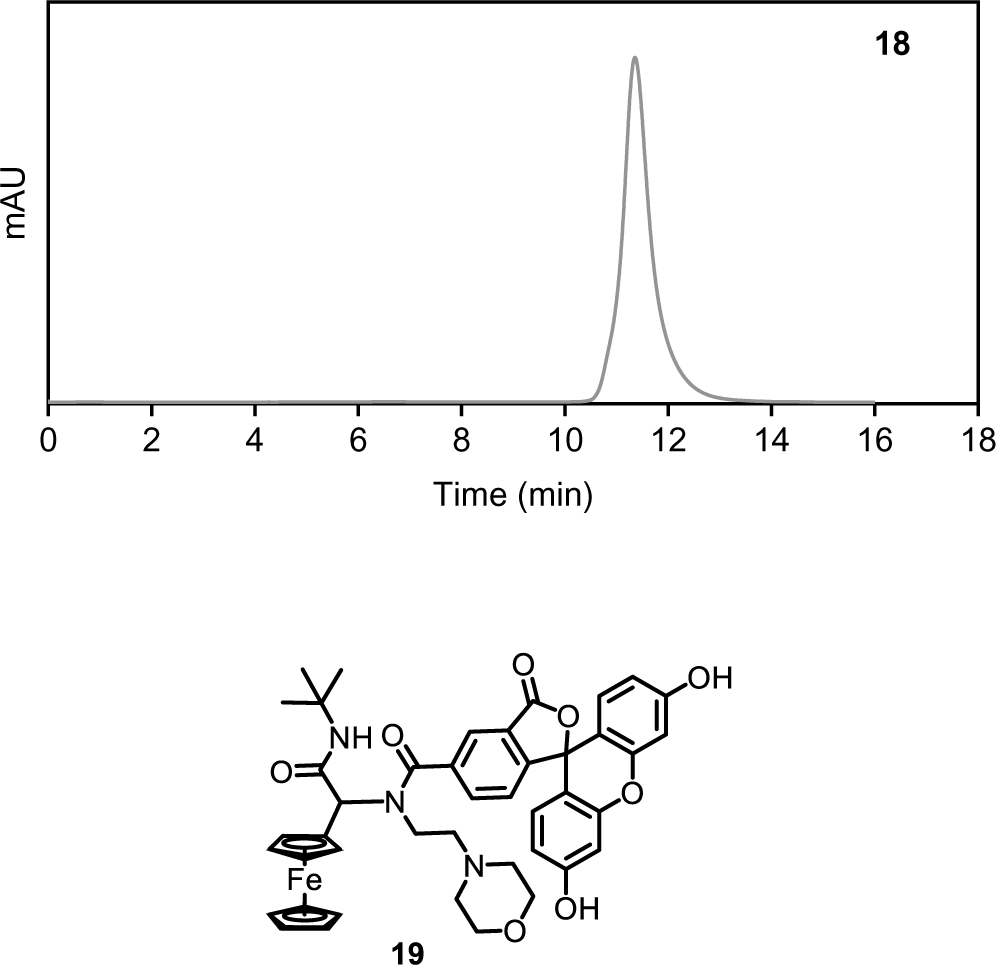

N-(2-(tert-butylamino)-2-oxoethyl)-3’,6’-dihydroxy-N-(2-morpholino-1-ethylferrocene)-3-oxo-3H-spiro[isobenzofuran-1,9’-xanthene]-5-carboxamide

HPLC-MS of **19**. m/z = 785.2400 Da [M(**19**)], m/z = 786.08 Da [M(**19**+H)]

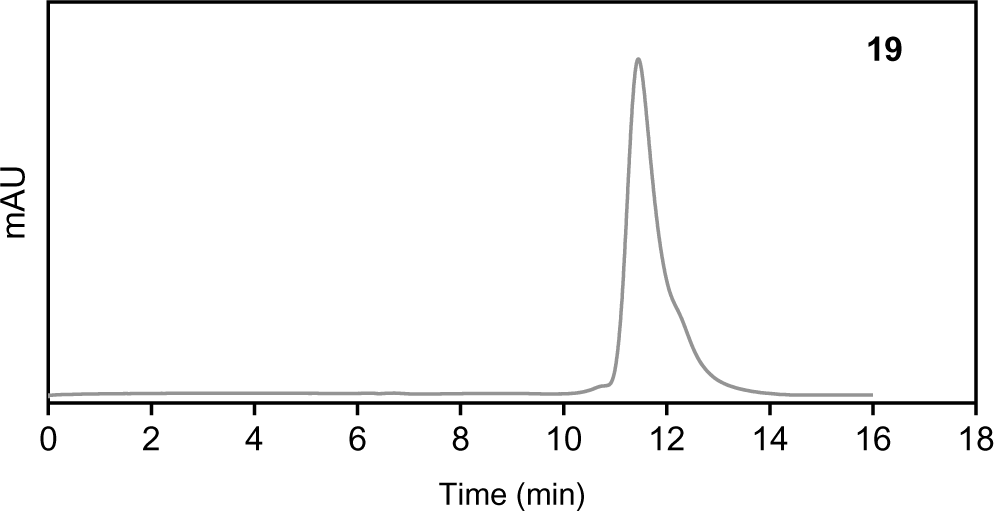

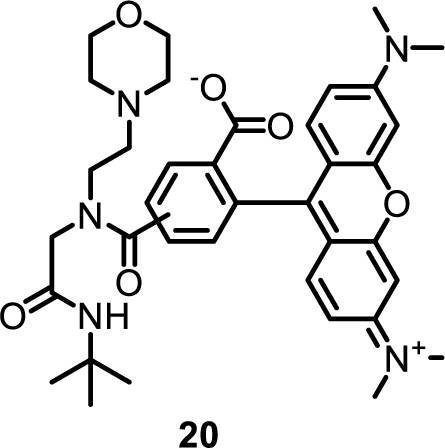

4,5-((2-(tert-butylamino)-2-oxoethyl)(2-morpholinoethyl)carbamoyl)-2-(6-(dimethylamino)-3-(dimethyliminio)-3H-xanthen-9-yl)benzoate

HPLC-MS of **20**. m/z = 656.17 Da [M(**20**+H)].

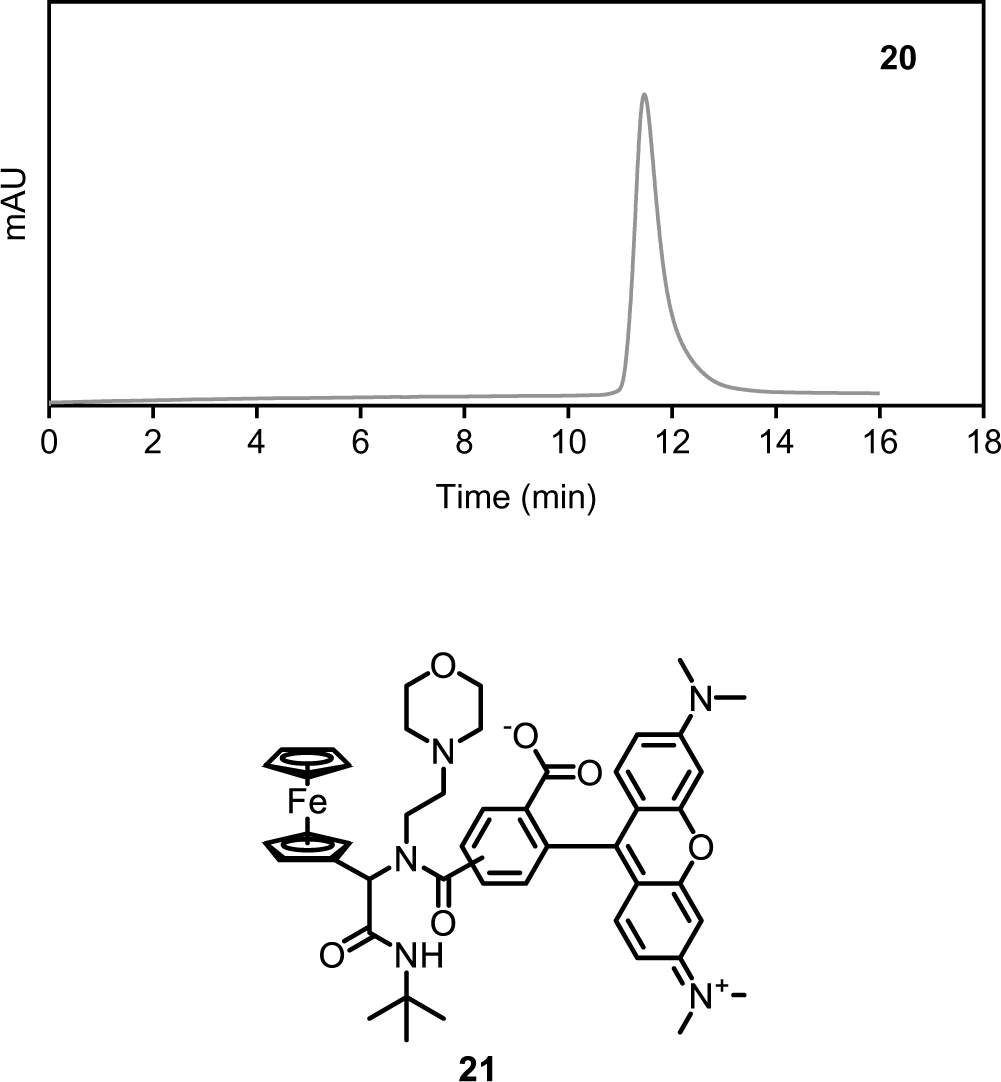

4,5-((2-(tert-butylamino)-2-oxoethyl)(2-morpholino-1-ethylferrocene)carbamoyl)-2-(6-(dimethylamino)-3-(dimethyliminio)-3H-xanthen-9-yl)benzoate

HPLC-MS of **21**. m/z = 840.29 [M(**21**)].

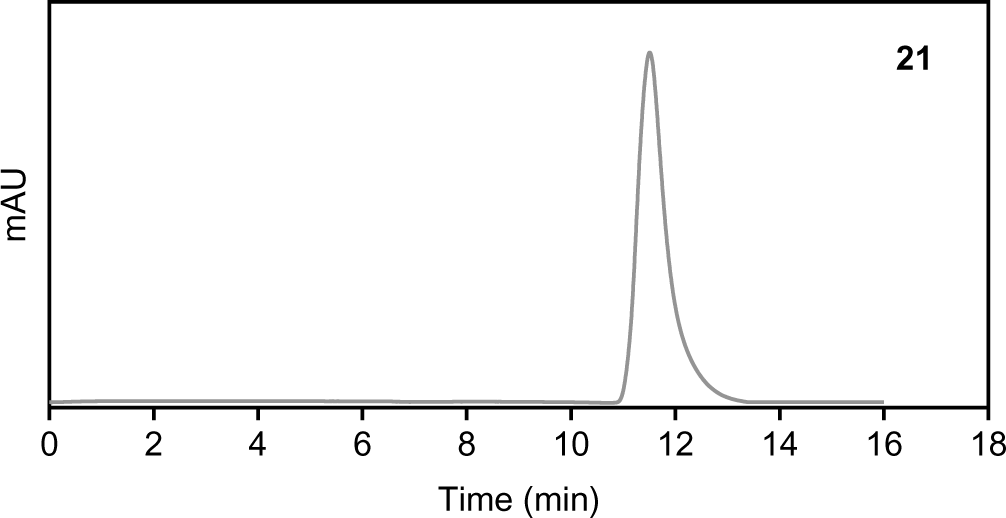

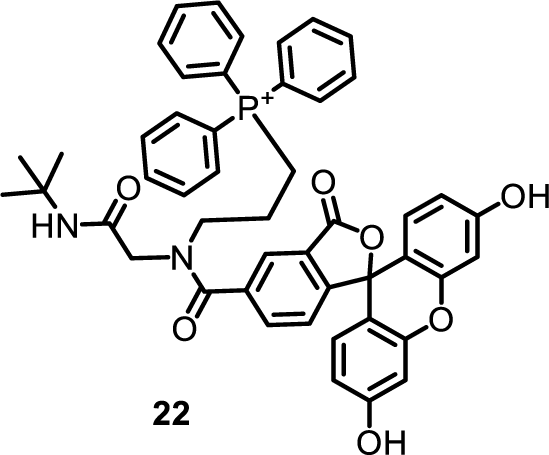

(3-(N-(2-(tert-butylamino)-2-oxoethyl)-3’,6’-dihydroxy-3-oxo-3H-spiro[isobenzofuran-1,9’-xanthene]-5-carboxamido)propyl)triphenylphosphonium

HPLC-MS of **22**. m/z = 791.15 Da [M(**22**)].

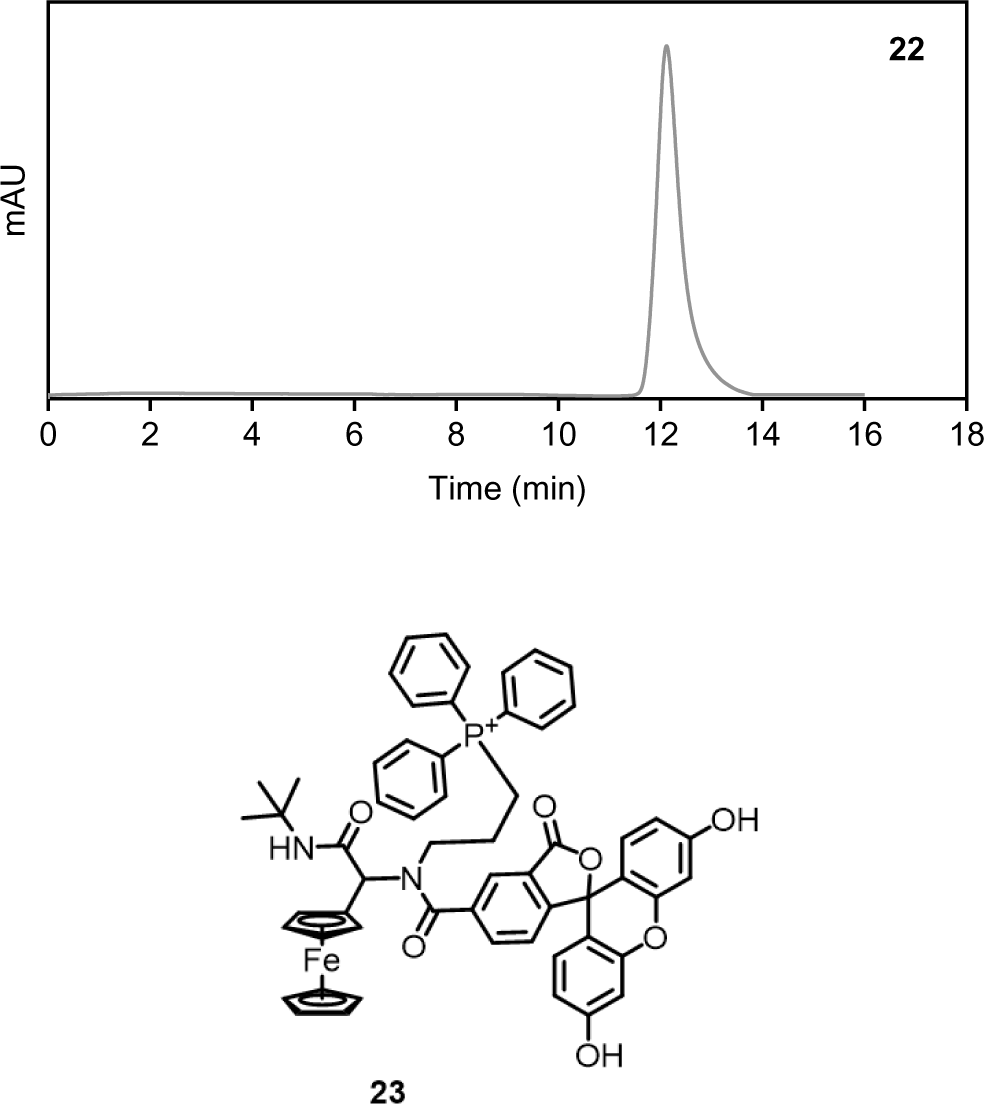

(3-(N-(2-(tert-butylamino)-2-oxo-1-ethylferrocene)-3’,6’-dihydroxy-3-oxo-3H-spiro[isobenzofuran-1,9’-xanthene]-5-carboxamido)propyl)triphenylphosphonium

HPLC-MS of **23**. m/z = 795.29 Da [M(**23**)].

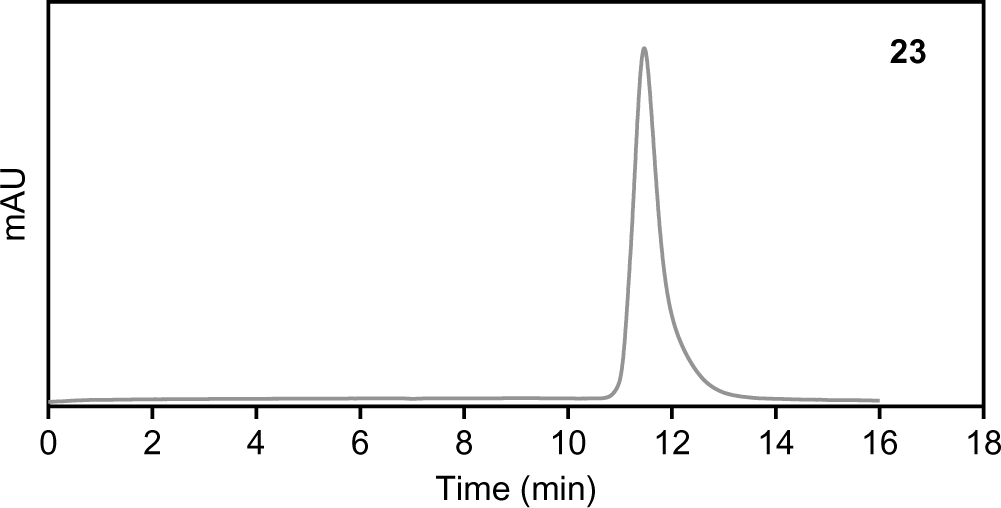

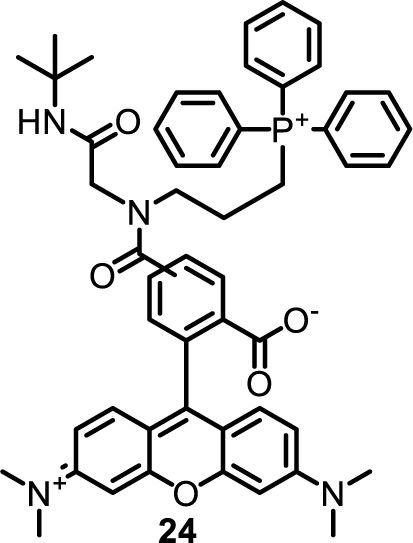

4,5-((2-(tert-butylamino)-2-oxoethyl)(3-(triphenylphosphonio)propyl)carbamoyl)-2-(6-(dimethylamino)-3-(dimethyliminio)-3H-xanthen-9-yl)benzoate

HPLC-MS of **24**. m/z = 845.36 Da [M(**24**)].

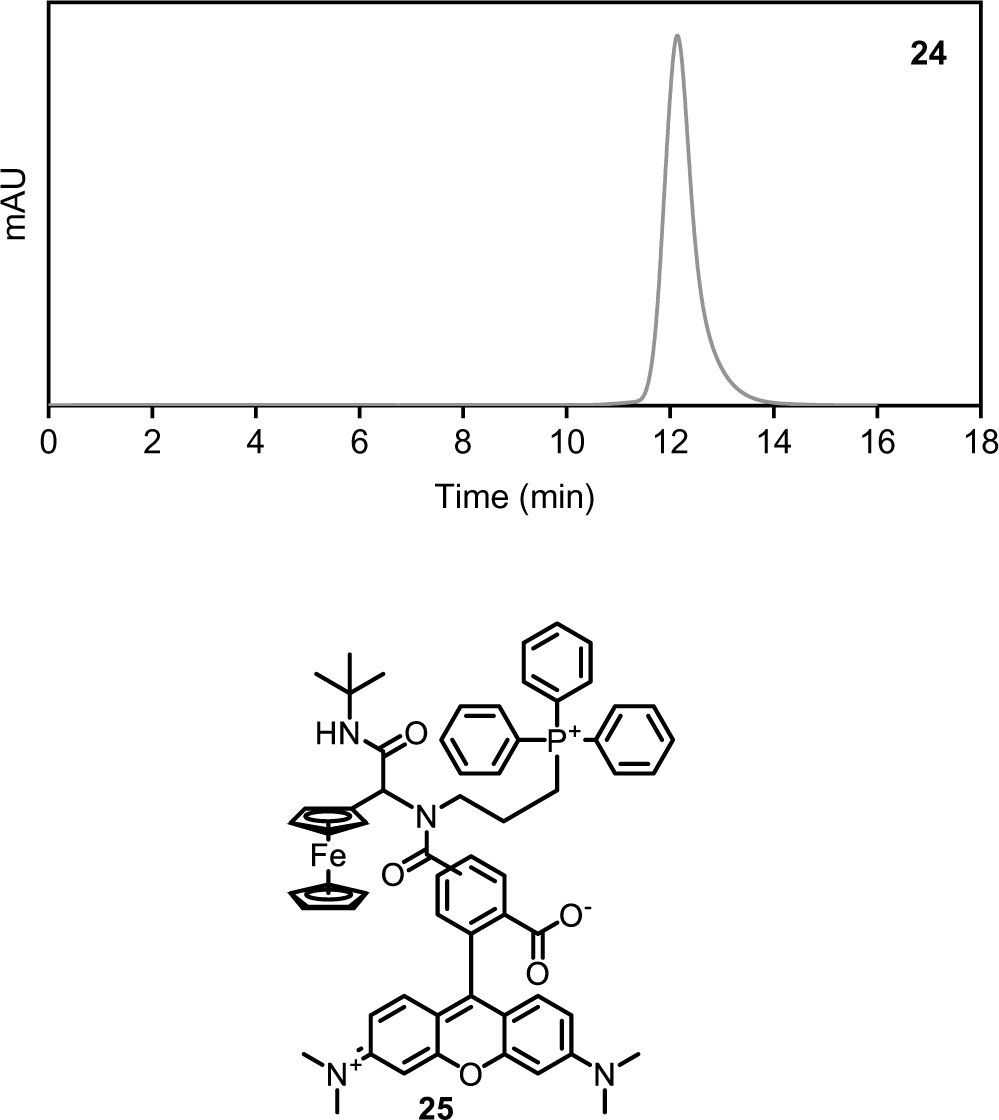

4,5-((2-(tert-butylamino)-2-oxo-1-ethylferrocene)(3-(triphenylphosphonio)propyl)carbamoyl)-2-(6-(dimethylamino)-3-(dimethyliminio)-3H-xanthen-9-yl)benzoate

HPLC-MS of **25**. m/z = 1029.3767 Da [M(**25**)].

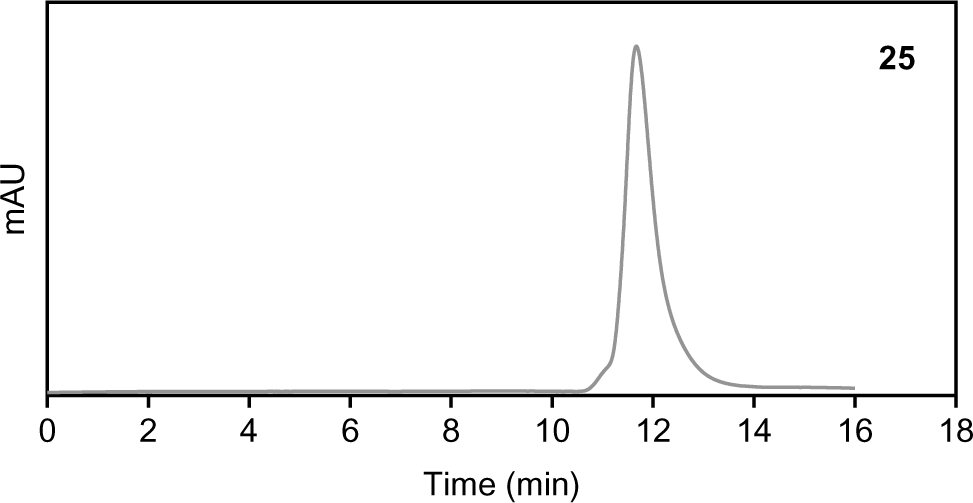

### Part III: Spectroscopy, microscopy and biochemical assays

#### Steady-state absorption and fluorescence spectroscopy

Steady-state absorption spectra were recorded on a UV-2000 UV-VIS spectrometer (UNICO, USA). Steady-state fluorescence spectra were recorded on a FluoroMax-4 luminescence spectrometer (HORIBA, USA).

#### Transient absorption spectroscopy

The femtosecond pump-probe transient absorption measurements were conducted using a regenerative amplified Ti:Sapphire laser system (Coherent). The laser source emitted light at 800 nm with a pulse width of 100 fs, and a repetition rate of 1 kHz. This 800 nm pulse is bifurcated by a beam splitter into two pathways: one channel directs the light into an optical parametric amplifier (OPerA Solo, Coherent Inc.) to produce pump light at wavelengths (670 nm and 540 nm) requisite for the experiment, while the other channel, following traversal through a delay line, guides the light into a transient absorption spectrometer (Helios-Eos Fire, Ultrafast Systems). Here, it is focused onto a sapphire crystal to generate supercontinuum probe light. The pump light, now at a frequency of 500 Hz, and the probe light are both focused onto the sample after modulation by a chopper, which reduces the pump light frequency to 500 Hz. The probe light, having been absorbed by the sample, conveys information regarding the excited and ground states of the particles and subsequently impinges on a fiber optic probe. Employing a time delay line to adjust the probe light’s delay relative to the pump light’s arrival at the sample enables acquisition of the spectral information of the sample particles at various time delays post-excitation. The samples were placed in cuvettes and analyzed under ambient conditions. Finally, the pump wavelength was set to its absorption maximum around 490 nm. A time window of 7.6 ns was then sampled with a broadband white-light pulse covering the spectral range from 325–650 nm.

#### Synthesis of double-stranded DNA containing adjacent ferrocene on different fluorophores

Synthetic oligomers (P1: Dye-5’-TAA TAT TCG ATT CCT CTG GAC G-3’, P2: 5’-CGT CCA GAG GAA TCG AAT ATT A-3’-Ferrocene) were received in HPLC-purified quality from Sangon Inc. (Shanghai, China). The dye modified on P2 included aminofluorescein (FAM), tetramethylrhodamine (TMR), Alexa Fluor 488 (AF488), Cy3, Cy5, sulfo-Cy5 (sCy5), Alexa Fluor 647 (AF647), ATTO647N, CY5.5, and Cy7. Double-stranded DNA was achieved form the hybridization of P1 and P2 by a typical DNA annealing protocol. In brief, mix 10 µL of 100 µM P1 with 10 µL complementary 100 µM P2 in 80 µL annealing buffer (20 mM Tris-HCl, 50 mM NaCl, 1 mM EDTA, pH = 7.6), heat the mixed solution to 95°C for 5 min and cool down to 4 °C in 1°C/min steps.

#### Cell culture

The human cervical adenocarcinoma (HeLa) and normal human colon mucosal epithelial cell line (NCM460) cell lines were cultured separately in flasks using Dulbecco’s modified Eagle’s medium (DMEM, Gibco) with 10% fetal calf serum (FCS, Gibco), penicillin (100 μg mL^-1^), and streptomycin (100 μg mL^-1^) at 37°C with 5% CO_2_ in a humidified environment. The cell count was determined using a Petroff-Hausser cell counter (USA).

#### Confocal fluorescence imaging

HeLa cells were seeded at a density of 5.0 × 10^4^ (per dish) onto Glass Bottom Dishes with a diameter of 35-mm and incubated for 6 hours at 37°C. The cells were then treated with culture medium containing the probe (100 nM) and subjected to different experimental conditions. Prior to imaging, the cells were rinsed three times with PBS and kept in PBS. The fluorescence of the cells was visualized with a Zeiss LSM880 NLO Confocal Microscope System equipped with a LD C-Apochromat 63×/1.15 W Korr objective (Zeiss). Parameter settings for laser intensity, exposure time, and objective lens were maintained for all samples during image acquisition. The cells were observed under a Finally, ImageJ software was utilized to analyze the micrographs.

To investigate the colocalization effect of our probes, HeLa cells were initially incubated with the synthesized probe at a concentration of 100 nM for 6 hours. followed by the staining of lysosomes and mitochondria using LysoTracker Deep Red (50 nM) and MitoTracker Deep Red (50 nM) for 30 minutes, respectively. The colocalization of the probes were determined using confocal laser fluorescence microscopy (Zeiss).

#### Intracellular •OH evaluation

HeLa cells were seeded into 35-mm confocal dishes at a density of 5 × 10^4^ cells per dish and incubated at 37°C for 24 hours. Following sufficient growth, the cells were treated with a specified concentration of **29**/**30** and incubated for an additional 6 hours at 37°C. Subsequently, they were exposed to a blue LED light at an intensity of 0.1 W/cm^2^for varying durations. After exposure, the cells were washed thrice with PBS and placed in fresh medium for another 6-hour period. For further analysis, the treated cells underwent a 60-minute incubation with MitoROS™ OH580 at 37°C. Post incubation, following another PBS wash, the cells were analyzed using confocal scanning fluorescence microscopy. The laser source was set at 543 nm for excitation, with emission recorded between 570 and 620 nm.

#### *In vitro* cytotoxicity evaluation

To assess cytotoxicity, HeLa cells (5.0 × 10^5^) were seeded into the wells of a 96-well plate and allowed to settle for 12 hours. Subsequently, the medium was removed, and the cells were washed twice with PBS. They were then incubated with a culture medium containing varying concentrations of the synthesized probe for 12 hours. For the groups subjected to laser irradiation, the cells were exposed to LED light at an intensity of 0.1 W/cm^2^ for 30 minutes, utilizing an automatic cooling device to prevent overheating. The total irradiation period did not exceed 15 minutes. In contrast, control cells were incubated with 100 μL of culture medium devoid of the probe. Following incubation, cells were washed with PBS, and CCK-8 solution was added. The cells were then incubated for an additional 2 hours at 37°C. The absorbance of each well was measured at a wavelength of 450 nm using a microplate reader. Relative cell viability (%) was determined using the formula (*A*_test_/*A*_control_) × 100%.

#### Flow cytometry analyses

HeLa cells were collected, washed with PBS twice and treated with probes as indicated. Stained cells were analysed with a BD FACSAria III flow cytometer and data were processed using FlowJo software.

#### Cytokines release assays

The levels of IL-18 and IL-1β in cell culture supernatants were quantified using the ELISA method ^[51^], following treatment with probes **27–30** as indicated.

#### Western blotting test

Briefly, the cells that had been treated were harvested and lysed in a cell lysis buffer. After that, the lysate was subjected to centrifugation at 4 ◦C. The concentrations of the collected proteins were determined using a BCA Protein Assay Kit. The protein samples were then separated using sodium dodecyl sulfate–polyacrylamide gel electrophoresis (SDS-PAGE, 12%) and transferred onto polyvinylidene difluoride (PVDF) membranes. The membranes were blocked with 3% bovine serum albumin for 2 h and then incubated with specific primary antibodies at 4 ^◦^C overnight. Following this, the membranes were washed with TBST buffer (1 X Tris-buffered saline and 0.1% Tween-20) and then incubated with fluorescent secondary antibodies for 2 h. The fluorescent bands were visualized using an ECL Kit through the process of chemiluminescence. The primary antibodies used were anti-full-length GSDMD (Abcam, #ab210070), anti-cleaved GSDMD-N (Abcam, #ab215203), anti-GSDME (Abcam, #ab215203), anti-cleaved GSDME-N (Abcam, #ab222408), anti-cleaved caspase-1 (CST, #4199) and anti-cleaved caspase-3 (CST, #9664) and anti-*α*-tublin (proteintech, #11224-1-AP).

### Part IV: Supplementary figures

**Figure S41.**
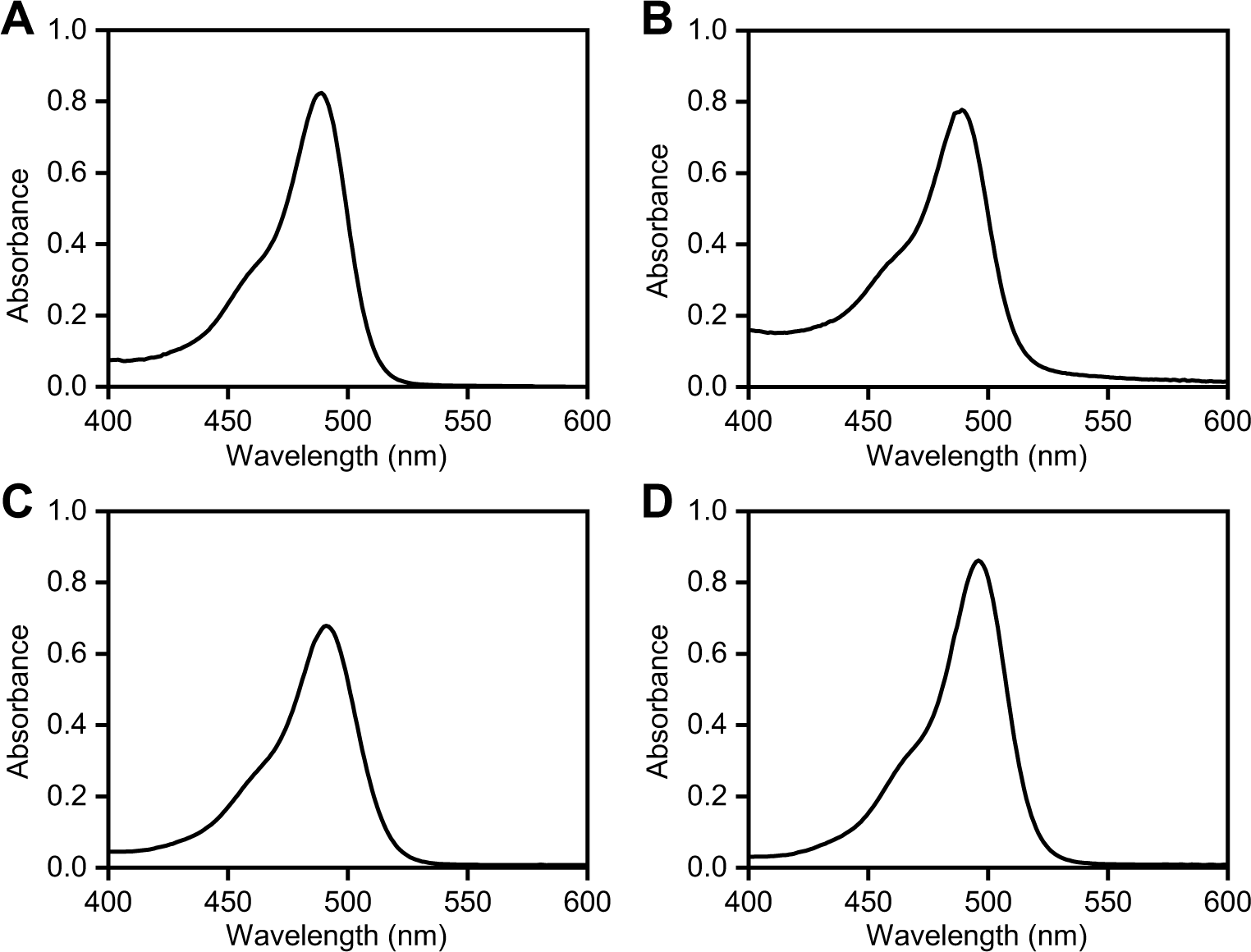
UV-visible absorption spectra of **27** (A), **28** (B) and **29** (C) and **30** (D) in pH 7.4 PBS solutions with 1% DMSO.

**Figure S42.**
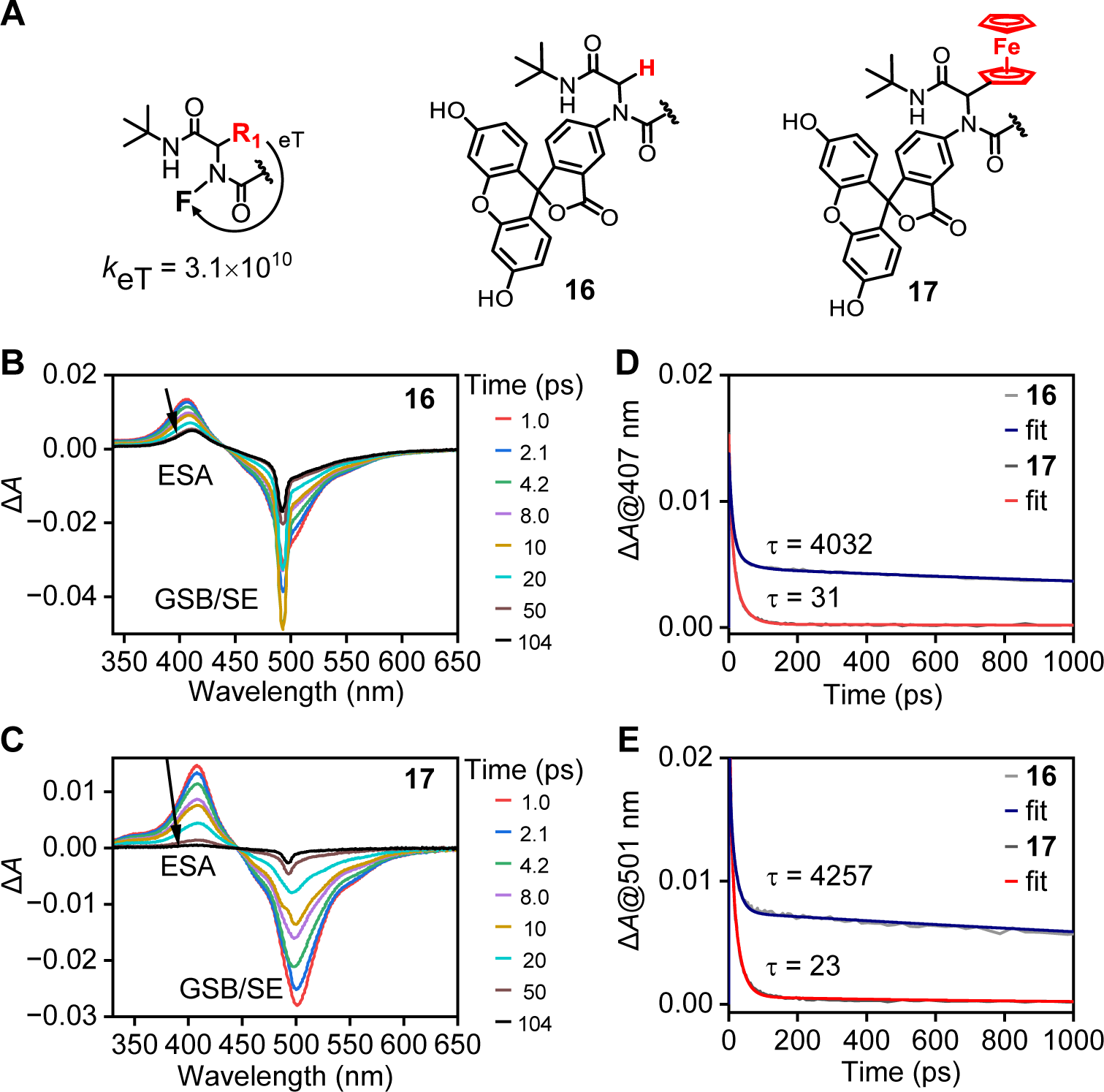
(A) The fluorophore moiety is located at the N-terminal end of the U-4CR framework within the α-acylamino structure including compound **16** and **17**. (B, C) Transient absorption spectra and (D, E) time evolutions and the derived lifetime constants of the transient absorption signals at 412 and 501 nm for **16** and **17**. They were excited at a wavelength of 490 nm with excitation power of 100 μW.

**Figure S43.**
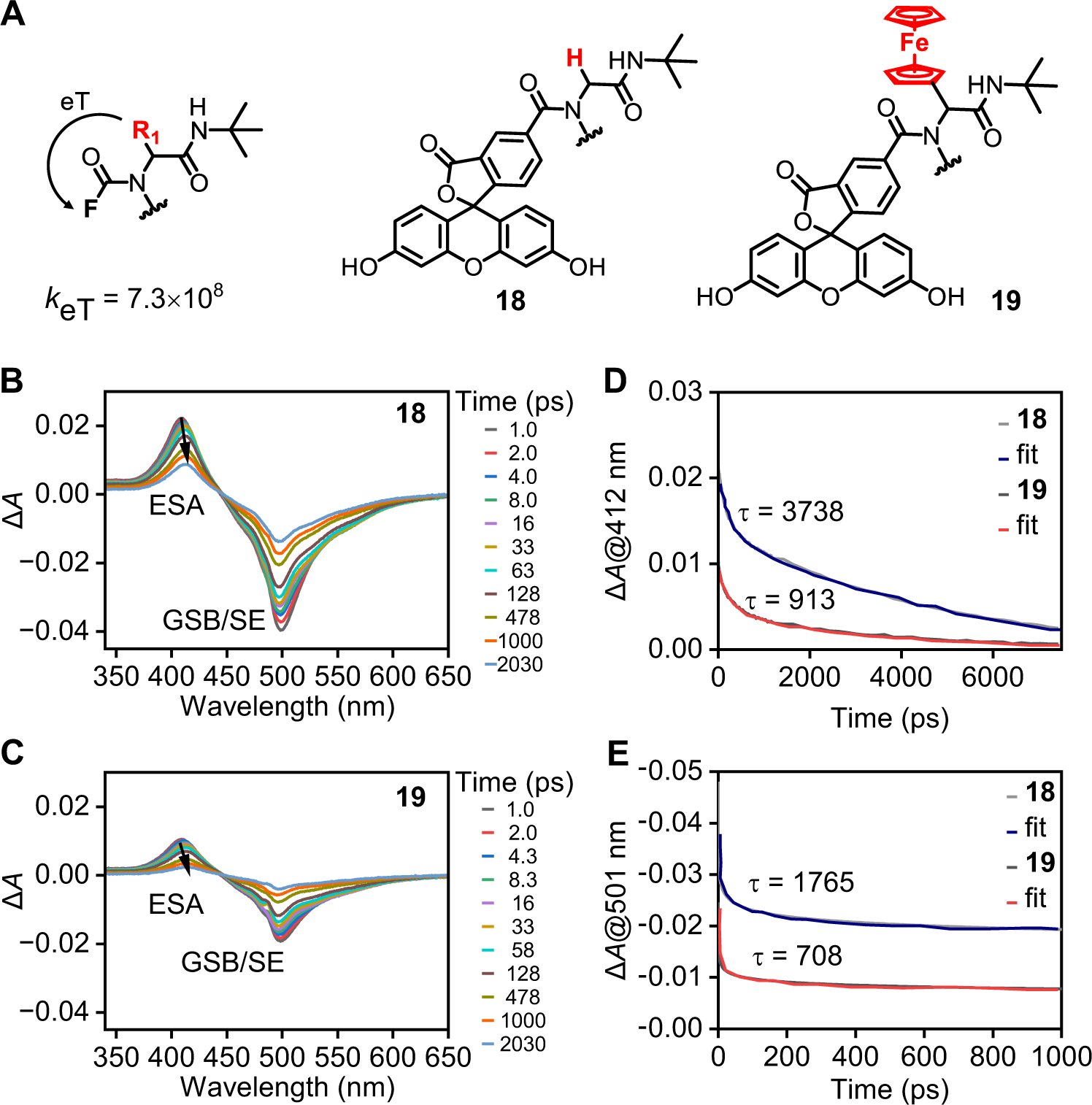
(A) The fluorophore moiety is located at the C-terminal end of the U-4CR framework within the α-acylamino structure including compound **18** and **19**. (B, C) Transient absorption spectra and (D, E) time evolutions and the derived lifetime constants of the transient absorption signals at 412 and 501 nm for **18** and **19**. They were excited at a wavelength of 490 nm with excitation power of 100 μW.

**Figure S44.**
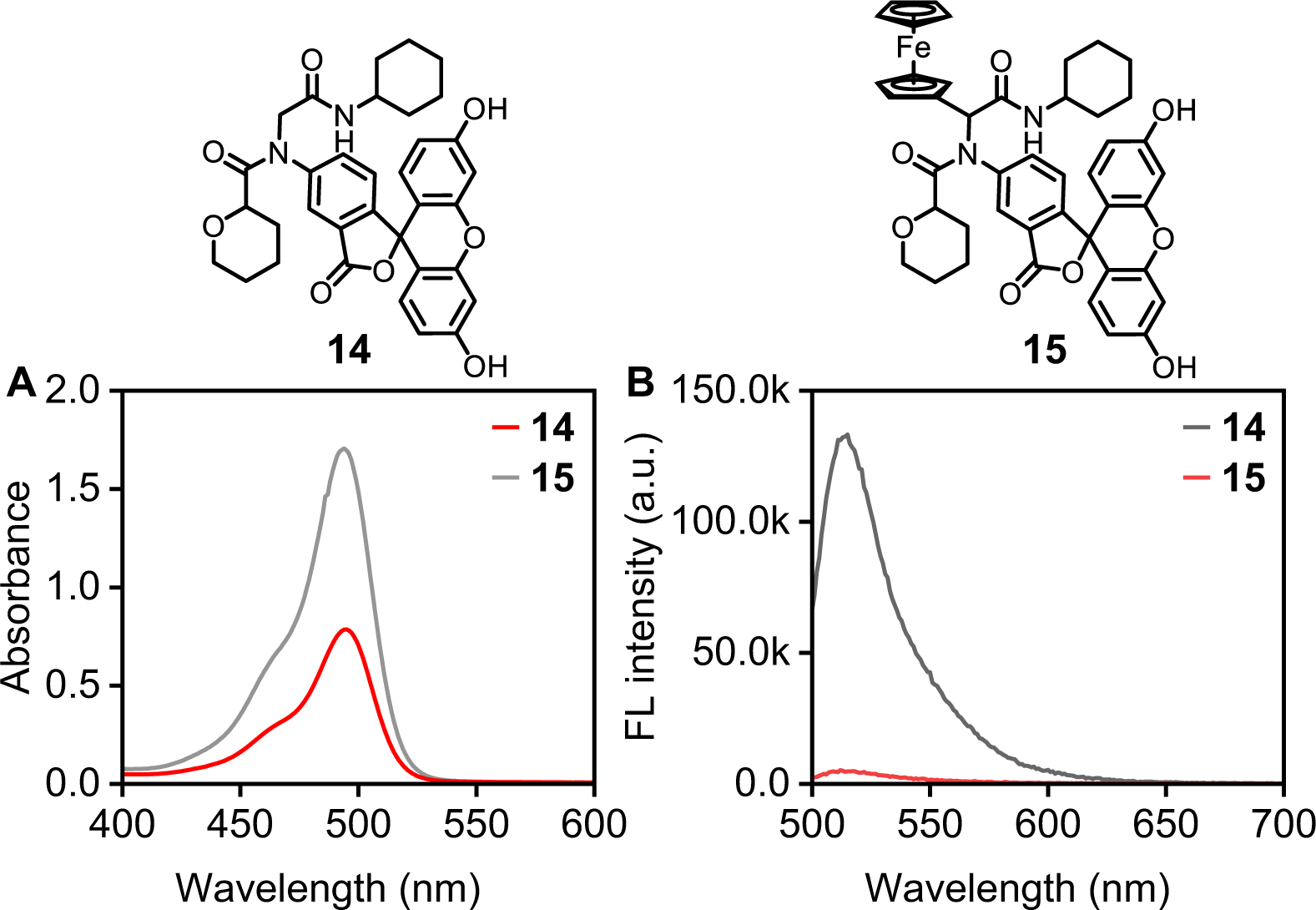
UV-Visible absorption spectra (A) and fluorescence emission spectra (B) of **14** and **15** in pH 7.4 PBS solutions with 1% DMSO. The probe concentrations are 10 μM in (A) and 1.0 μM in (B), respectively.

**Figure S45.**
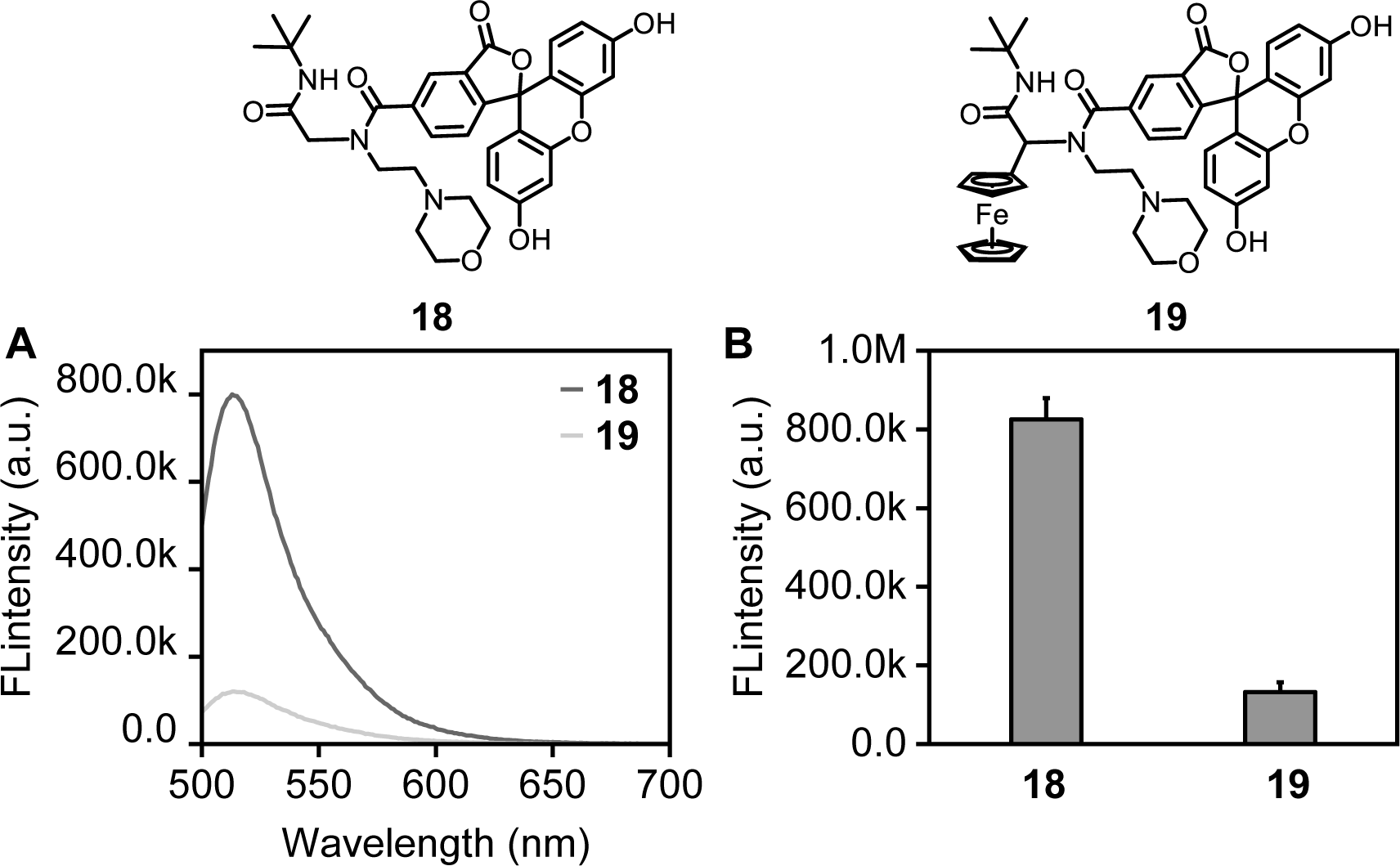
Fluorescence emission spectra (A) and fluorescent intensity at 513 nm (B) of **18** and **19** at a concentration of 10 μM in pH 7.4 PBS solution with 1% DMSO.

**Figure S46.**
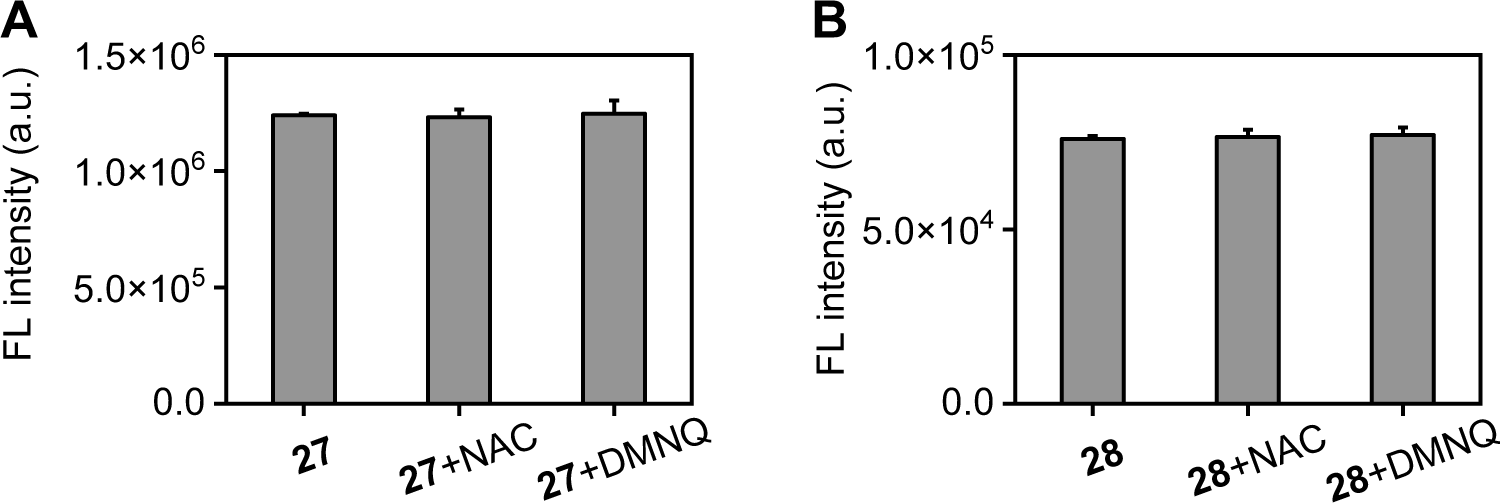
Comparison of the effects of N-acetylcysteine (NAC) and 2,3-dimethoxy-1,4-naphthoquinone (DMNQ) on the fluorescence intensity of compounds **27** and **28** in cell culture medium.

